# Sleep is required for odor exposure to consolidate memory and remodel olfactory synapses

**DOI:** 10.1101/2020.11.24.395228

**Authors:** Rashmi Chandra, Fatima Farah, Fernando Muñoz-Lobato, Anirudh Bokka, Kelli L. Benedetti, Chantal Brueggemann, Fatema Saifuddin, Julia M. Miller, Joy Li, Eric Chang, Aruna Varshney, Vanessa Jimenez, Anjana Baradwaj, Cibelle Nassif, Sara Alladin, Kristine Andersen, Angel J. Garcia, Veronica Bi, Sarah K. Nordquist, Raymond L. Dunn, Kateryna Tokalenko, Emily Soohoo, Vanessa Garcia, Sukhdeep Kaur, Malcolm Harris, Fabiola Briseno, Brandon Fung, Andrew Bykov, Hazel Guillen, Decklin Byrd, Emma Odisho, Bryan Tsujimoto, Alan Tran, Alex Duong, Kevin C. Daigle, Rebekka Paisner, Carlos E. Zuazo, Matthew A. Churgin, Christopher Fang-Yen, Martina Bremer, Saul Kato, Noëlle D. L’Étoile, Miri K. VanHoven

**Author notes:** Indicates co-first authors (in alphabetical order) (Rashmi Chandra, Fatima Farah, Fernando Muñoz-Lobato). Indicates co-second authors (Anirudh Bokka, Kelli L. Benedetti). Indicates co-corresponding authors (in alphabetical order) (Noëlle D. L’Étoile, Miri K. VanHoven).

## Abstract

Animals with complex nervous systems demand sleep for memory consolidation and synaptic remodeling. Here we show that though the *Caenorhabditis elegans* nervous system has a limited number of neurons, sleep is necessary for both processes. In addition, it is unclear in any system if sleep collaborates with experience to alter synapses between specific neurons and whether this ultimately affects behavior. *C. elegans* neurons have defined connections and well-described contributions to behavior. We show that spaced odor-training and post-training sleep induce long-term memory. Memory consolidation, but not acquisition, requires a pair of interneurons, the AIYs, which play a role in odor-seeking behavior. In worms that consolidate memory, both sleep and odor conditioning are required to diminish inhibitory synaptic connections between the AWC chemosensory neurons and the AIYs. Thus, we demonstrate in a living organism that sleep is required for events immediately after training that drive memory consolidation and alter synaptic structures.

## INTRODUCTION

Understanding how sleep promotes the consolidation of a specific memory is one of the foremost challenges in biology (Jenkins and Dallenbach, 1924). Many animals with complex nervous systems require sleep to consolidate memory (Crocker and Sehgal, 2010). For example, humans need sleep to learn skilled motor tasks (Walker et al., 2002). Mice and rats also consolidate memories during sleep after training; they learn to overcome their innate attraction to dark spaces when the dark side of a box is paired with a foot shock, and this learning requires sleep after training (Impey et al., 1998; Stubley- Weatherly et al., 1996). *Drosophila* can learn to avoid their innate attraction to light if it is paired with the noxious stimulus quinine, and this learning is disrupted in animals that have spontaneously fragmented sleep (Le Bourg and Buecher, 2002; Seugnet et al., 2009; Seugnet et al., 2008). Similarly, *Aplysia* can learn that initially appetitive foods are inedible after repeated trainings, and the durability of this learning requires sleep (Vorster and Born, 2017). The nervous systems in these organisms are large, ranging from tens of thousands to hundreds of billions of neurons (Akhmedov et al., 2014; Erö et al., 2018; Lent et al., 2012; Scheffer et al., 2020). It is not known if nervous system complexity dictates the need for sleep in memory formation.

Learning and memory are thought to require modulation of specific synapses in circuits impacted by training, such that functional changes in synaptic signalling that result from training can eventually be transitioned into physical changes in synaptic structures. These physical changes in the structure of synaptic connections are termed synaptic consolidation (Asok et al., 2019). Early studies on memory in *Aplysia californica* demonstrated that single trial training increased both glutamate release from the presynaptic neuron, and glutamate receptor activity in the postsynaptic neuron (Siegelbaum et al., 1982). After repeated training, changes in transcription and translation were shown to drive stable changes in synaptic structure that correlated with memory consolidation (Bailey and Chen, 1983; 1989; Montarolo et al., 1986; Schacher et al., 1988). The transcriptional changes that drive these and many other examples of long-term, consolidated memory require cAMP response element binding protein (CREB) (Bartsch et al., 2000; Dash et al., 1990). Work in mammalian systems and Drosophila established the evolutionary conservation of these mechanisms of synaptic consolidation (Asok *et al*., 2019; Crocker and Sehgal, 2010; Tononi and Cirelli, 2014). However, we lack a clear understanding of how sleep affects the synaptic structures between the specific neurons that are required for a long-term memory.

The best studied processes by which sleep impacts the nervous system are those that reduce synapses across broad regions of the brain and likely promote plasticity. The synaptic homeostasis hypothesis (SHY) proposes that during sleep, large groups of synapses are downscaled to compensate for global increases in synaptic strength during wakefulness as neurons respond to stimuli. This role of sleep in downscaling synapses is thought to maintain synaptic strength within a functional range (Cirelli and Tononi, 2020; Raven et al., 2018). Many studies support this hypothesis (De Vivo et al., 2017; Diering et al., 2017; Huber et al., 2013; Li et al., 2009; Norimoto et al., 2018). For example, the axon-spine interface in mouse motor and sensory cortices decreases approximately 18% after sleep, although large synapses are spared (De Vivo *et al*., 2017). Similarly, the size of most spines in the mouse primary motor cortex is reduced during sleep, although some spines show an increase (Diering *et al*., 2017). In addition, synaptic potentiation has been observed during sleep (Aton et al., 2014; Frank et al., 2001; Seibt and Frank, 2019). Synaptic downscaling is thought to spare synapses that have been strengthened by learning during wake, an important property if memories that result from synaptic strengthening are to be maintained (Cirelli and Tononi, 2021).

Despite these important findings, an understanding of sleep’s function in consolidating long-term memory at the cellular and synaptic levels remains elusive. Specifically, how sleep affects synapses between single cells with known contributions to memory during consolidation remains unknown in any system (Cirelli and Tononi, 2020; Tononi and Cirelli, 2014). Such an understanding would require discovering the exact cells that are required for a memory, which would allow the discovery of how precise connections are modulated by training and sleep. *C. elegans* has a compact nervous system, with ∼0.3% of the number of neurons in *Drosophila*. The *C. elegans* nervous system is also well-characterized, with stereotyped functions for many neurons and the entire synaptic connectome elucidated. Thus, *C. elegans* provides an opportunity to examine how sleep might change connections between the neurons that control behaviors altered by learning and memory.

Though nematodes have not been reported to require sleep for memory or synaptic modulation, memory and sleep have both been studied separately for over a decade in *C. elegans*. Olfactory memory formation has been studied in *C. elegans* as it provides a unique approach to investigating ethologically relevant stimuli. Odors, universally powerful signals for food and its contaminants, are salient cues. Butanone may serve as such a cue for *C. elegans*. This volatile chemical, emitted from both nutritious and infectious bacteria (Labows et al., 1980; Worthy et al., 2018a; Worthy et al., 2018b), could be associated with either positive or negative experiences. Thus, the mechanism for learning to ignore butanone could be an evolutionarily conserved trait to avoid further ingestion of pathogenic bacteria. This may explain why *C. elegans* can be trained to seek butanone (Kauffman et al., 2011; Torayama et al., 2007; Vohra et al., 2017), or avoid it altogether (Tsunozaki et al., 2008).

Previous studies have elucidated the identities of neurons required for butanone chemotaxis (Albrecht and Bargmann, 2011; Chalasani et al., 2007; Gordus et al., 2015; Gray et al., 2005) and learning (Cho et al., 2016). Butanone is sensed by AWC^ON^, one of the two AWC olfactory neurons (Troemel et al., 1999), which primarily form synapses with three pairs of interneurons: the AIYs, AIAs and AIBs (Albrecht and Bargmann, 2011; Chalasani *et al*., 2007; Gordus *et al*., 2015; White et al., 1986; Witvliet et al., 2021). Together, these neurons coordinate movement during chemotaxis. Short-term plasticity was induced after one pairing of butanone with removal from food (Colbert and Bargmann, 1995; L’Etoile and Bargmann, 2000). The AIA interneurons were found to allow *C. elegans* to learn to ignore the odor in this one-cycle training paradigm (Cho *et al*., 2016). In addition to the rapid one-cycle learning, paradigms have been developed to study long-lasting olfactory memory that is induced after multiple rounds of odor pairing with lack of food interspersed with feedings (Kauffman et al., 2010).

Prior studies on sleep have documented that the hallmarks of sleep are conserved across the animal kingdom: periods of quickly reversed immobility, increased arousal threshold, homeostatic compensation, stereotypical posture and broadly altered patterns of neuronal activity (Nichols et al., 2017; Skora et al., 2018; Trojanowski and Raizen, 2016). There are a number of stressors that have been shown to induce sleep in *C. elegans*: development requires a period of lethargus to allow successful growth and molting (Avery, 1993; Driver et al., 2013; Raizen et al., 2006; Raizen et al., 2008), temperature (Van Buskirk and Sternberg, 2007), cellular stressors (Hill et al., 2014), DNA damage (DeBardeleben et al., 2017), prolonged swimming for more than four hours (Schuch et al., 2020), prolonged periods (16 hours) of starvation (Skora *et al*., 2018) as well as being held in a microfluidic device (Gonzales et al., 2019). The period of sleep can occur once the stressor has resolved as in the case of re-feeding after starvation, which results in satiety-induced quiescence (Gallagher and You, 2014; You et al., 2008). Each sleep trigger is likely to engage the salt-induced kinase (KIN-25/SIK), which is responsive to mobilized fat stores (Grubbs et al., 2020). This leads to activation of neurons in a sleep circuit that release somnogenic peptides (Trojanowski and Raizen, 2016). *C. elegans’* ALA interneuron triggers stress-induced sleep by releasing the FMRFamide FLP-13 among other neuropeptides (Nath et al., 2016) and the interneurons RIS and RIA regulate lethargus at least in part by releasing FLP-11 and NLP-22, respectively (Nelson et al., 2014; Turek et al., 2016; Turek et al., 2013). These neuropeptides, conserved in fly and fish, in turn engage GABAergic pro- sleep circuits (Lee et al., 2017; Lenz et al., 2015; Meeusen et al., 2002).

In this study, we asked whether *C. elegans*, like the more complex organisms studied, requires sleep for long-term memory consolidation. Using a training paradigm adapted from Kauffman and colleagues (Kauffman *et al*., 2011),we show that three cycles of training with butanone in the absence of food produces an olfactory memory that makes trained *C. elegans* lose their innate attraction to butanone. Remarkably, we found this olfactory memory is dependent upon sleep. We discovered that animals have increased bouts of sleep for at least six hours after training, and if sleep is disrupted during the initial two hours either by mechanical disturbance or removal from food, the animals do not consolidate their memory. Therefore, when odor training is followed immediately by sleep, we find that *C. elegans* retain the memory for a large fraction of their reproductive lifespan. These results uncover a specific temporal function of sleep that benefits memory.

We next asked how sleep organizes a neural circuit to store memory. We found that AIB or AIY interneurons can compensate for each other during learning. However, AIYs are consistently required for memory consolidation while AIBs have a more variable contribution. We visualized synapses between AWC chemosensory neurons and the AIY interneurons using the split-GFP based trans-synaptic marker Neuroligin-1 (NLG-1) GFP Reconstitution Across Synaptic Partners (GRASP). We find that butanone-trained animals have significantly reduced AWC-AIY connections when compared with their control buffer-trained counterparts 16 hours after training. Interrupting sleep for two hours immediately after training abolishes this synaptic reduction 16 hours post-training. Thus, odor-training and post-training sleep are both required to modify these specific synapses. We sought to understand the dynamics of synaptic remodelling and found that synapses are significantly reduced in both odor and control (buffer-trained) animals by the end of the two-hour period required for sleep to consolidate memory. However, synaptic levels in these groups become distinct from each other during the following fourteen-hour period, so that by 16 hours after training, synaptic levels in butanone-trained animals are lower than in their control buffer-trained counterparts.

Our study reveals that sleep is required for odor memory consolidation in a simple nervous system. This is the first study to report sleep-dependant synaptic structural plasticity in *C. elegans.* This provides a new level of precision and granularity in understanding learning and memory. This work strongly indicates that nervous system complexity is not a requirement for memory consolidation and modification of synaptic connections by sleep.

## RESULTS

### Olfactory conditioning induces sleep and long-lasting memory

In order to understand how long-lasting memories are formed and retained, we adapted a spaced, repeated conditioning paradigm from Kauffman and colleagues (Kauffman *et al*., 2011) (Figure 1A). This paradigm takes advantage of *C. elegans* ability to learn to ignore butanone, an innately attractive odor that is emitted by both nutritious (Worthy *et al*., 2018b) and pathogenic bacteria (Worthy *et al*., 2018a) that are found in *C. elegans* natural environment. In this training paradigm, the negative, unconditioned stimulus is removal from food and is paired with either butanone diluted in buffer (1:10,000 dilution) or, as a control, buffer alone. Learning is defined as the difference in a population of animals’ attraction to butanone after training with butanone as compared to training with buffer (see inset in Figure 1A). Attraction is quantified using a chemotaxis assay developed by Bargmann and colleagues (Bargmann et al., 1993) (Figure 1A). In this assay, animals are placed onto a 10cm diameter petri dish filled with a layer of agar. A point source of 0.1% (11mM) diluted butanone is placed opposite a similar source of diluent (ethanol), and animals are introduced at an origin equidistant from each source. Each point source is supplemented with sodium azide to paralyze the animals once they reach it. After at least 90 minutes of roaming, the position of each animal on the plate is scored. The chemotaxis index is calculated by subtracting the number of animals at ethanol from the number at butanone and dividing by the total number of worms on the plate (not including the origin) (Figure 1B and inset formula).

**Figure 1.**
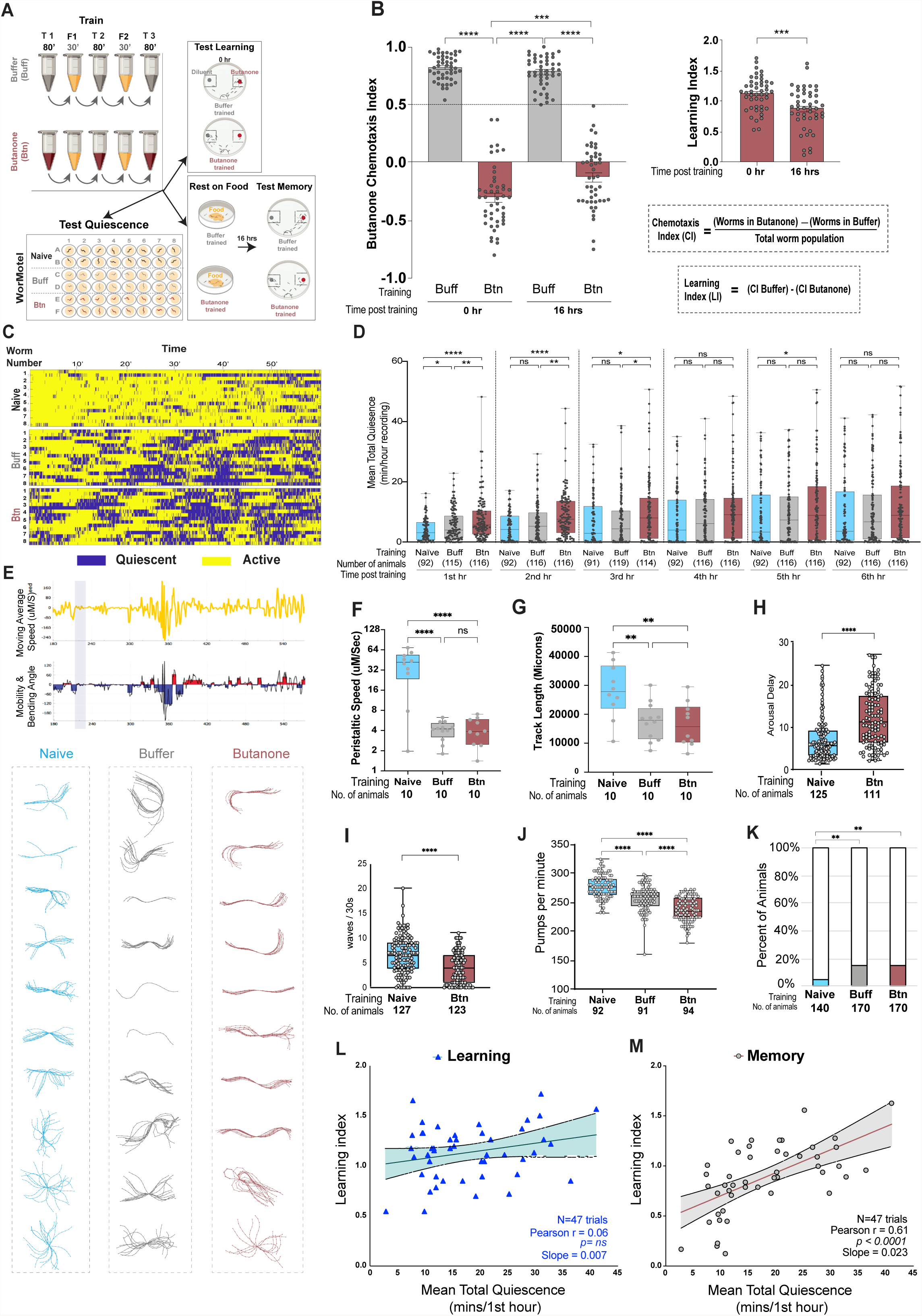
Spaced training paradigm induces quiescence and olfactory memory (A) Training and subsequent analysis. Populations of animals were subjected to repeated, spaced training with either butanone or buffer (control) after which the populations of animals were split into thirds and tested for learning (chemotaxis assays), placed on food (OP50) containing plates for 16 hours then tested for memory (chemotaxis assays) or loaded singly in to individual wells of a WorMotel device that contains food. Movement was imaged at 1frame/3 sec. to measure quiescence. **(B) Learning and 16 hour memory**. Chemotaxis index (CI, see equation below the graph) and learning index (LIs) of each population of wildtype animals performed at the indicated time after training. N = 47 trials. Throughout the paper, all CI and LI data points represent a trial of at least >100 animals and each was performed on a separate day. One way ANOVA was performed on the CI and two-tailed t-test on the LI (*** p<0.005; **** p<0.0005). **(C) Quiescence analysis.** Raster plot showing activity (yellow) and quiescence (blue) of naïve, buffer or butanone trained individuals at the indicated time after training. Each row represents one individual animal. **(D) Quiescence over 6 hours.** Each animal’s mean total quiescence (minutes) over one hour is plotted as a data point for the indicated hour elapsed after training, numbers of animals are indicated below. One-way ANOVA with Bonferroni’s correction, ****P < 0.0001, ***P < 0.001, ** P < 0.01, *P < 0.05, N = 7 trials. **(E) Posture analysis.** Top, (yellow trace) example of one animal’s moving average speed and below, blue and red histogram of midpoint- bending angle in the hour after training. Shaded bar indicates the period with the lowest speed and least bending angle. Bottom, 10 consecutive frames from this period were captured, animals were skeletonized and their skeletons overlayed. The average midpoint bending angle over the 10 frames is indicated for each animal. **(F) Movement speed.** The speed of each animal in **E** is plotted. One-way ANOVA with Bonferroni’s multiple correction (****P<0.0001). **(G) Track length.** The distance each animal in E traveled is plotted. One-way ANOVA with Bonferroni’s multiple correction (****P<0.0001). **(H) Arousal delay.** Time (seconds) before blue LED light flashes and 1.2KHz of vibrations evoked a complete sinusoidal escape wave was recorded and scored. Data show individual animals in three separate trials. **(I) Activity following arousal**. The number of sinusoidal waves completed in 30 seconds after exposure to blue LED light and 1.2 KHz vibrations stimuli. Additional animals from videos in **H** were analyzed. **H** and **I**, the Mann-Whitney µ -test was performed and N = number of animals assayed. **(J) Feeding rate.** Feeding rate: pharyngeal pumps per minute. Each point is one animal, 5 trials. One-way ANOVA with Bonferroni’s multiple correction (****P<0.0001). **(K) Feeding quiescence.** Fraction of animals on food not pumping for 4 seconds (colored bars). Z-test with Hochberg correction (**P<0.005). Total number of animals in 5 independent experiments indicated below. **(L) Learning immediately after training versus mean quiescence.** The LI of each population at t=0 (before recovery) is plotted versus the mean duration of quiescence in the hour after training. N = 47 trials, Pearson’s correlation coefficient is 0.06 P=ns. **(M) Memory 16 hours after training versus mean quiescence.** The LI after 16 hours of recovery on food is plotted versus the mean duration of quiescence in the hour after training. N = 47 trials, Pearson’s correlation coefficient (r) is 0.61, P <0.0001.

The bulk of a buffer-exposed population is attracted to the odor butanone, and their chemotaxis index (CI) is usually between 0.6 and 0.9 while the butanone-exposed population gives rise to a lower, sometimes negative CI between 0.4 and -0.7 (Figure 1B, left). Each point on the graphs represents a CI resulting from an independent day’s population of >50 animals. By subtracting the CI of the buffer-trained population from that of its siblings in the butanone-trained cohort immediately after training, we quantify how much the population has learned (Learning Index, LI). We found that repeated training typically produces LIs from 0.4 to 1.2 (Figure 1B, right, 0 hour after training).

We then asked how long this difference in attraction lasts when animals are placed on food. We found that the difference between the buffer- and the butanone-trained populations’ CIs was significant, even after the animals spent 16 hours on a petri dish with food (Figure 1B, left, time post-training 16 hours). The difference in the buffer- and butanone-trained cohorts’ CIs at this time point 16 hours after training is considered to be a measure of memory. The amount of memory kept over 16 hours, as assessed by the LI after 16 hours on food, ranged from 0.12 to 1.62 with a mean of 0.87. This is similar to the LI of 0.6, that was seen in Kaufmann et. al. 2010, with seven shorter cycles of training and higher concentrations of butanone. We found that this memory persists up to 24 hours (Figure S1A and S1B). This conditioning paradigm did not interfere with odor detection in general, as we found that three cycles of training with butanone did not affect the animals’ ability to sense and track the food-associated odor diacetyl, which is sensed by AWA chemosensory neurons, 16 hours after training.

Similarly, butanone-trained animals sense and track the food-associated odor benzaldehyde as well as their buffer-trained control cohorts (Figure S1C and S1D). Butanone is sensed by the AWC^ON^ cell, while benzaldehyde is sensed by both AWC^ON^ and AWC^OFF^ neurons (Wes and Bargmann, 2001). Thus butanone training does not impair the animal’s general ability to chemotax.

We observed that after training, the animals appeared quiescent when compared to age-matched naïve animals. We asked whether *C. elegans* sleep after conditioning. There are a number of macroscopic metrics by which sleep is assessed in all animals: decreased movement over time, reduced feeding rates and increased arousal latency (Trojanowski and Raizen, 2016). We first assessed individual animals’ movement over time using the WorMotel, a video-based setup adapted from Churgin and colleagues (Churgin et al., 2017) (Figure 1A, bottom, Video S1). Individual animals are placed into wells of the WorMotel, which is a 48-well PDMS device filled with solid nematode growth media that is supplemented with a lawn of bacteria (Video S1). The WorMotel keeps animals separated from one another, and locomotion is monitored using an automated imaging system (Churgin *et al*., 2017). The output of one hour of recording is shown in Figure 1C. We measured the length of time an individual animal remained quiescent using a frame by frame subtraction method and a frame rate of 3 seconds (Churgin *et al*., 2017). If the pixel displacement was zero from one frame to the other and remained zero for 9 consecutive frames (27 seconds), then the animal was considered to be undergoing a quiescence bout, and a blue line marked the bout on the raster plot. Conversely, when animals moved, pixel displacement between the consecutive frames was greater than zero and a yellow mark noted 3 seconds of movement (Figure 1C). By summing the duration of each bout over that period, we traced the bouts of quiescence for six hours after training (Figure 1D) to identify the period when animals exhibited the highest amount of sleep. To compare the amount of sleep among age-matched cohorts, we divided the WorMotel into three groups of animals: naïve, buffer-trained and butanone-trained animals. Each group contained 16 animals in one WorMotel containing 48 chambers.

We found that the animals that underwent training slept significantly more than the naïve animals during the first hour after training (Figure 1D), wherein the butanone-trained population slept the most, followed by the buffer-trained, and then the naïve populations. After the first hour, the differences in the quiescence bouts decreased between trained and untrained animals, as naïve and buffer-trained animals started to sleep more (Figure 1D). As the trained animals exhibited the highest amount of sleep immediately after training, we quantified the quiescent bouts of naïve, buffer- and butanone-trained populations for 47 trials during the first hour after training and found that the mean total quiescence of all butanone-trained animals was greater than that of either the naïve or the buffer-trained cohorts during the first one-hour period after training (mean total quiescence: naïve = 5.63 ± 0.4 minutes, buffer-trained = 10.84 ± 0.6 minutes, butanone-trained = 13.94 ± 0.8 minutes).

We then asked if in addition to reduced movement and bouts of inactivity, the trained animals showed other hallmarks of sleep, namely a stereotypical posture (Iwanir et al., 2013; Lawler et al., 2021). We found that in periods of lowest activity, both butanone and buffer trained animals took on one of two basic postures: a C curve either with a straight tail such as observed previously (Iwanir et al., 2013; Lawler et al., 2021) or a very straight slightly sinusoidal posture (Figure 1E). Next, we asked if they exhibited increased arousal latency, and reduced feeding. We examined arousal latency by asking how long it takes an animal to make an escape maneuver after being exposed to a noxious stimulus.

We stimulated animals with a blue light pulse (a noxious stimulus) in conjunction with mechanical vibrations (1 KHz frequency) from a piezoelectric buzzer and found that it took longer for butanone-trained than untrained (naïve) animals to execute an escape response (Figure 1H: median of 7 seconds for untrained vs 12 for trained). The trained animals also executed fewer body bends after the stimulus is removed (Figure 1I; median of 7 sinusoidal waves for untrained vs 4 for trained). Reduced movement after arousal may reflect a sleep debt incurred by the stimulation or that animals are tired or both.

We next examined feeding by counting pharyngeal pumping rates. These rates were significantly decreased in both buffer- and butanone-trained populations as compared to naive (Figure 1J; median of 276 pumps/min for naïve, 260 buffer- and 236 butanone-trained). When we asked what proportion of animals paused pumping for at least 4 seconds (Hill et al., 2014), we found that 15% of butanone or buffer-trained animals paused while none of the naïve animals paused for even 3 seconds (Figure 1K). Thus, by these four criteria, increased quiescence, stereotypical posture, increased arousal latency and reduced feeding, animals that undergo training either with buffer or butanone exhibit sleep.

Post-training sleep in a population of flies has been shown to benefit memory formation (Ganguly-Fitzgerald et al., 2006), but whether the amount of sleep correlates directly with higher olfactory learning or increased olfactory memory remains elusive. We reasoned that if sleep is important for memory consolidation even in the simple nervous system of *C. elegans*, memory measured 16 hours after training might correlate with sleep duration in the first hour after training. Therefore, we determined if there was a correlation between the amount of learning (LI at 0 hours after training) or memory (LI at 16 hours after training) with the amount of post-training sleep in the first hour after training. We found that the amount a population learns is not correlated with sleep duration in the first hour after training (Pearson r = 0.06, N = 47 trials, slope = + 0.007) (Figure 1L). However, we found that memory 16 hours after training correlated strongly with the amount of sleep a population exhibited (Pearson r = 0.61, N = 47 trials, P < 0.0001) (Figure 1M). Thus, the more a population slept after training, the better the memory consolidation.

### CREB is required for long-term memory after training

CREB, the cyclic AMP response element binding protein, is required for long-term memory formation in flies, *Aplysia,* mice (Ganguly-Fitzgerald et al., 2006; Silva et al., 1998), and in *C. elegans (* Kauffman et al., 2010). Thus, we asked if CREB plays a role in this olfactory learning paradigm. We found that *crh-1(tz2)/CREB* mutants learned as well as wildtypes when they were tested immediately after training (Figure 2A and B compare first pairs of brick wildtype to teal *crh-1(tz2)*), but they fail to keep the memory 16 hours after training. We conclude that though learning after three cycle training does not require CREB, memory at 16 hours does require this transcription factor. Thus, the long-term memory induced by three cycles of training is likely to be transcription-dependent.

**Figure 2.**
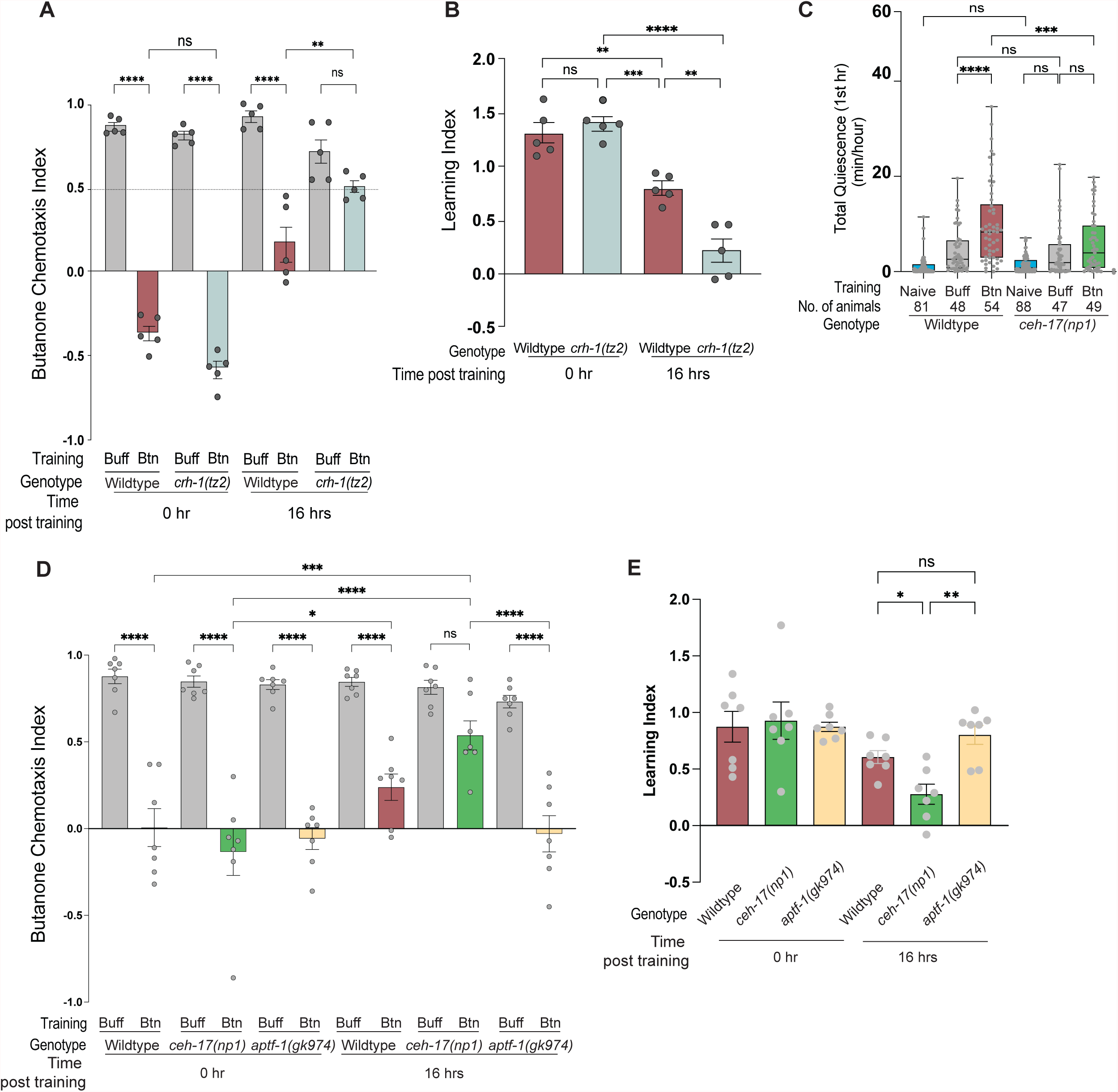
Long-term olfactory memory requires the sleep-promoting ALA neuron. **(A)** CIs and **(B)** LIs for wild type animals and CREB-defective *crh-1(tz2)/CREB* mutants immediately after training and 16 hours after recovery on food. One way ANOVA with Bonferroni’s multiple correction (**** P<0.0001, ***P < 0.001, ** P < 0.005 and (ns) is P>0.05), N = 5 trials. **(C)** Mean total quiescence in the first hour after training. Quiescence in naïve (untrained), buffer and butanone trained wild type and ALA defective *ceh-17(np1)* animals was examined in the WorMotel. One way ANOVA with Bonferroni’s multiple correction. (**** P<0.0001, ***P < 0.001, and (ns) is P > 0.05). N = 5 trials. **(D)** CIs and **(E)** LIs of wild type, *ceh-17(np1)* and *aptf-1(gk974)*. Two-way ANOVA of CIs with Bonferroni’s multiple correction show **** P<0.0001, ***P < 0.001, ** P < 0.005 and (ns) is P>0.05. One way ANOVA of LIs show ** P < 0.005, * P < 0.05 and (ns) is P > 0.05. N = 7 trials.

### Sleep-promoting ALA neuron is required to induce sleep and retain memory after training

The ALA neuron in *C. elegans* produces stress induced sleep (Hill et al., 2014a; Miyazaki et al., 2022; Nath et al., 2016b). Therefore, we tested if sleep produced by our spaced training paradigm requires ALA neuron. *ceh-17* mutants cannot undergo postembryonic differentiation of ALA neuron; therefore, ALA neuron is absent in *ceh-17* mutants. We found that naïve *ceh-17* mutant sleep like wildtype animals (Figure 2C). However, after training *ceh-17* mutants exhibit not only lower quantity of sleep, the difference in the amount of sleep between buffer and butanone trained populations is also absent in *ceh-17* mutants (Figure 2C). This suggests that ALA neuron responds to butanone to induce sleep.

As the amount of sleep is directly proportional to the amount of memory retained at 16 hrs, we tested if *ceh-17* mutants that exhibit sleeplessness retains less memory (Figure 2D and 2E). Indeed, *ceh-17* mutants, which learned equally well as wildtype animals but underwent severe memory loss at 16 hrs. Beside ALA, the RIS interneuron is known to promote sleep (Turek *et al*., 2016). In our training conditions, we found RIS defective *aptf-1* is learning and retaining memory like wildtype animals (Figure 2D and 2E) indicating that RIS interneuron is not participating in memory formation in this spaced training paradigm.

### Sleep is necessary for long-term memory

We asked whether *C. elegans* with its compact nervous system requires sleep after training to consolidate memory. To keep animals from sleeping, we mechanically disturbed them for a period of two hours at different time points after training. We then asked if these animals retained memory 16 hours after training (Figure 3Ai). In order to mechanically disturb the animals, we had to reduce the viscosity of their food, adapted from Driver and colleagues (Driver *et al*., 2013), in order to allow shaking of the plate to sufficiently jostle the worms. To reduce the viscosity of the food, we resuspended OP50 *E. coli* in S Basal to make a slurry, which was added to the worms on the plates. To physically disturb the animals, we shook the plates for one minute out of every 15 (Figure 3Ai). We found that shaking prevented them from sleeping (Video S2). We quantified the number of colony-forming units of GFP-expressing OP50 in the intestines of animals placed in a slurry compared with animals placed on a bacterial lawn of GFP-carrying OP50 (Figure S2A). We determined that animals ate the same amount under either condition. Of note, animals also had similar learning and memory when fed OP50 with or without GFP (Figure S2B-C).

As we found that trained animals exhibit more sleep immediately after training (Figure 1D-E), we reasoned that disrupting sleep immediately after training might hamper memory retention. Therefore, we mechanically disrupted sleep of the trained animals in three two-hour periods after training (Figure 3Ai). We found that disrupting sleep in the first two hours after training blocked memory retention (Figure 3B and C). By contrast, cohorts that had been mechanically disturbed after the first two hours kept the memory (Figure 3B and C). This suggests sleep in the first two hours after training is required for memory, but sleep after this time is not.

**Figure 3.**
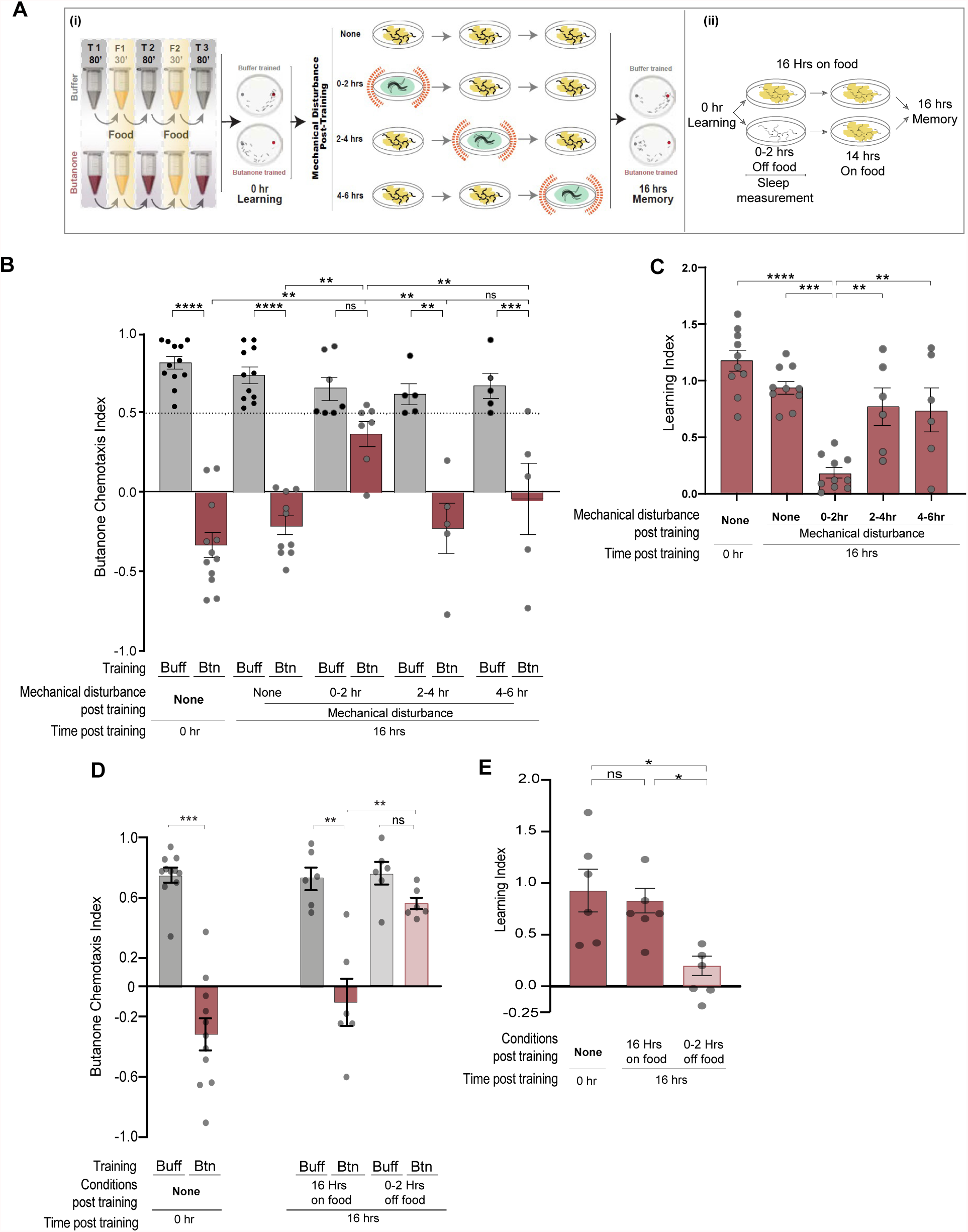
Disturbing animals immediately after training blocks memory. **(A)** Paradigms to disrupt sleep after training. **(i)** Mechanical disturbance: training is followed by shaking (red springs) animals in lower viscosity food every 15 minutes for two hours from 0 to 2, 2 to 4 or 4 to 6 hours after training. Supplemental Figure 3 indicates that animals eat during mechanical disturbance. Videos of sleep disruption are in Supplemental Video 3: Supports Figure 2. After disturbance, animals were allowed to recover on food without shaking until memory was assessed16 hours after training. **(ii)** Metabolic disturbance: training is followed by starvation. Animals are placed on food-less agar petri dishes for two hours immediately after training then moved to food-containing agar petri dishes for 14 hours before being tested for memory. Quiescence was measured during the period of starvation. **(B)** Cis and **(C)** LIs of mechanically disturbed populations. Cis were analyzed with a two-way ANOVA followed by Bonferroni’s multiple correction (****P < 0.0001, ***P < 0.001, ** P < 0.01, *P < 0.05, and (ns) is P > 0.05). N = 5 trials. LIs were analyzed using a one-way ANOVA with Bonferroni’s multiple correction (****P < 0.0001, ***P < 0.001, ** P < 0.01, and (ns) is P > 0.05). N < 5 trials. **(D)** CIs and **(E)** LIs of animals starved 0-2 hrs after training. One-way ANOVA with Bonferroni’s multiple correction (****P < 0.0001, ***P < 0.001, ** P < 0.01, *P < 0.05, and (ns) is P > 0.05). N > 5 trials.

Another way to disturb *C. elegans* sleep is to remove them from their bacterial food source, causing them to roam in search of their next meal (Gallagher and You, 2014; Gray *et al*., 2005; You *et al*., 2008). To further test if sleep is required for memory, we removed animals from food for the first two hours after training. To determine the extent of the sleep disruption, we analyzed a portion of the population’s behavior using the WorMotel. We found that animals placed in WorMotel wells without bacteria slept significantly less than animals placed in wells with bacteria (Video S2, Figure S2I). We found that animals removed from food for two hours after training failed to retain memory 16 hours post-training (Figure 3E and F). Thus, we show using two distinct mechanisms that sleep during the first two hours after training is necessary for long-term memory in *C. elegans*. This, to our knowledge, is the first example of sleep being required for memory consolidation in *C. elegans*.

### Sleep enhances long-term memory of butanone

As the amount of post-training sleep directly correlates with memory retention (Figure 1 I) and disruption of this sleep causes memory loss (Figure 4B and C), we asked if sleep could convert a short-term memory into a long-term memory. Though animals learn to ignore butanone after one cycle of training (as depicted in Figure 4A, bottom row) the memory is not maintained 16 hours later [Figure 4C and (Benedetti et al., 2021)]. We next asked if additional cycles of training with food in the absence of odor would increase quiescence. Swimming is energetically costly (Laranjeiro et al., 2017) and can induce sleep (Grubbs *et al*., 2020). As shown in Figure 4A, we altered our standard three-cycle training protocol (4A, top row, buffer training not shown) to include two cycles in which animals swim in liquid containing food and only one in which they swim in either odor or buffer.

**Figure 4.**
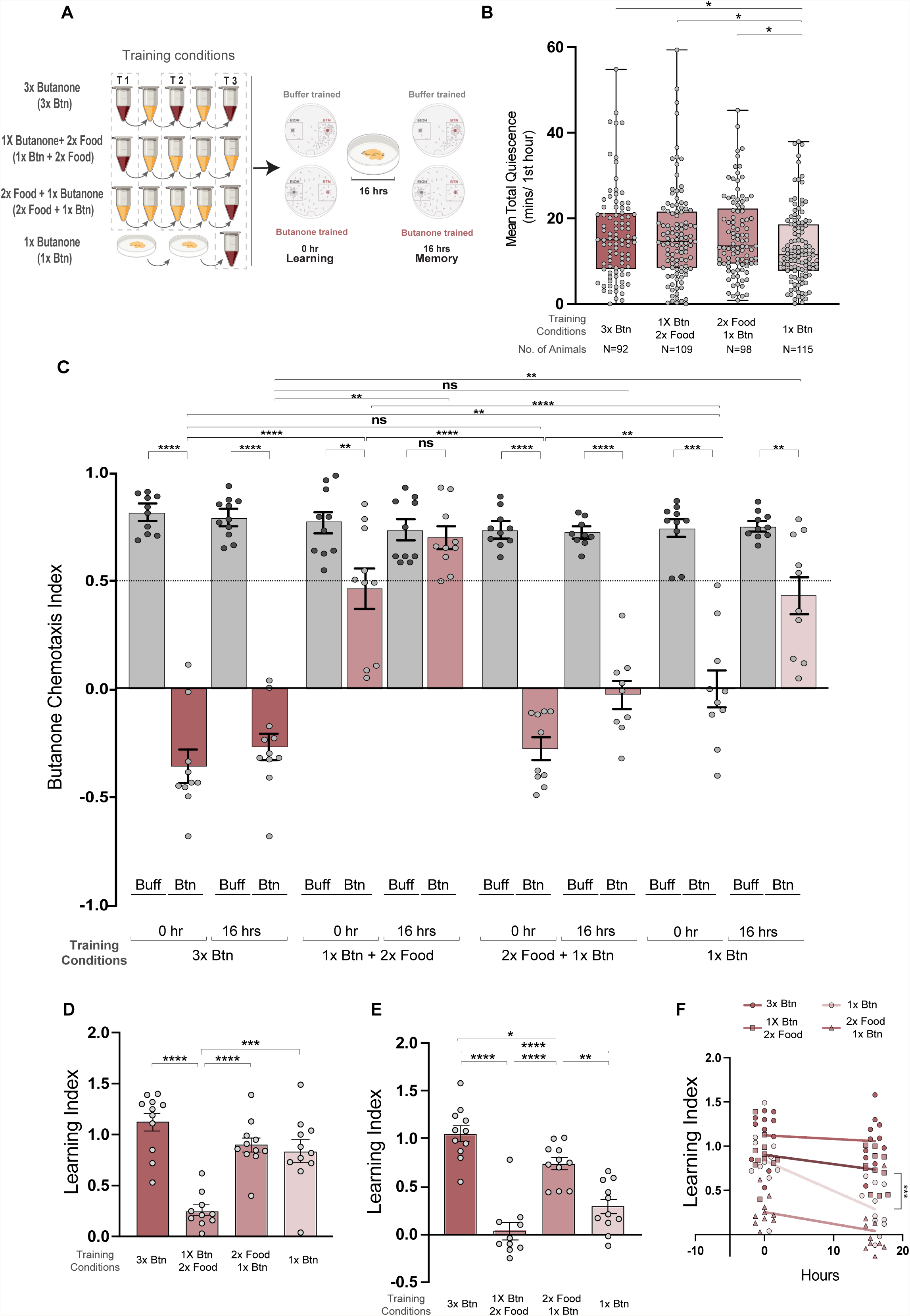
Increasing sleep increases memory. **(A)** Animals were trained as previously (top row, 3XBtn), or exposed to butanone during the first cycle followed by two cycles in buffer and food (1XBtn+2XFood), or two cycles in buffer and food followed by one cycle butanone in the last cycle (2XFood+1XBtn), or one cycle with butanone (bottom row, 1XBtn). After training, as previously, populations were assessed for quiescence (WorMotel), learning with a chemotaxis assay after training or chemotaxis assay after 16 hours on food. **(B)** Mean total quiescence after each training paradigm. Unpaired t-test with Welch’s correction (p < 0.05). N = 5 trials. **(C)** CIs of populations after each training paradigm at 0 hr or 16 hours post training. Two-way ANOVA with Bonferroni’s multiple correction (****P < 0.0001, ***P < 0.001, ** P < 0.01, *P < 0.05, and (ns) is P > 0.05). N = 10 trials**. (D)** LIs immediately after training. **(E)** LIs after 16 hours recovery on food. One-way ANOVA of with Bonferroni’s multiple correction is reported as ****P < 0.0001, ***P < 0.001, ** P < 0.01, *P < 0.05, and (ns) is P > 0.05. N = 10 trials. **(F)** Comparison of the slopes using linear regression show the amount of memory lost between 0 hr and 16 hrs for each training condition.

We found that animals that had only one cycle of swimming in butanone showed the lowest mean total quiescence (median of 13.19, Figure 4B, lightest pink, last bar) and those that had three cycles of swimming either in odor or in food showed more quiescence (median of 16.21, 15.94 and 16.01 minutes, Figure 3B red and medium pink, first three bars). Thus, sleep, as measured by total quiescence, is significantly increased if the number of cycles of training is increased from one to three. We then asked if this would convert the short-term into a long-term memory. We found that the cohorts that had two cycles of food training before one cycle of butanone training (third row in 3A) exhibited more long-term memory (LI = 0.73) than one cycle-trained animals. Thus, inducing sleep after a single cycle of odor training was sufficient to increase memory retention after 16 hours.

Interestingly, the cohorts that were trained with one cycle of butanone before the two cycles of food (row two in 4A) showed little learning, as the median LI at 0 hours after training for that cohort, 0.25, is significantly lower than the median LIs of the other groups that enjoyed three cycles of training, approximately 1.12 for 3 cycles butanone and 0.90 for two cycles of food followed by one of butanone (Figure 4D). The population that was trained first with butanone then with two food cycles had poor memory, as 16 hours post-training its median LI is 0.28 (Figure 4D), which is the smallest median LI observed. The 0-hour and 16-hour median LIs are each less than the corresponding LIs of the one cycle-trained animals, which are 0.255 and 0.016, respectively (Figure 4D and Figure S3). This could be because animals that are not allowed to sleep immediately after training cannot consolidate memory. The memory may also have been extinguished by the food training after butanone.

We next asked if the rate of memory loss was affected by sleep. We saw that in populations that were able to sleep after their last odor training, the rate of decay was significantly less than the one cycle-trained population that slept less (Figure 4E). This indicates that sleep reduces the rate at which memory is lost.

### Long-term memory does not accompany changes in AWC sensory neuron activity

We asked whether changes in the AWC sensory neuron response underlie some or all of the observed memory. One advantage of using the transparent *C. elegans* is that we can examine neuronal activity at the single neuron level in live animals at various time points during the sleep-induced memory stabilization. We used GCaMP3 (Tian et al., 2009) to monitor activity as reflected in calcium transients in AWC chemosensory neurons, the primary sensory neurons in the butanone chemotaxis circuit (Figure 4A) (Gordus *et al*., 2015). We imaged animals immediately after conditioning when they are repulsed by butanone, before the memory is consolidated, and after 16 hours on food when the memory is stable (Figure 5B-E).

**Figure 5.**
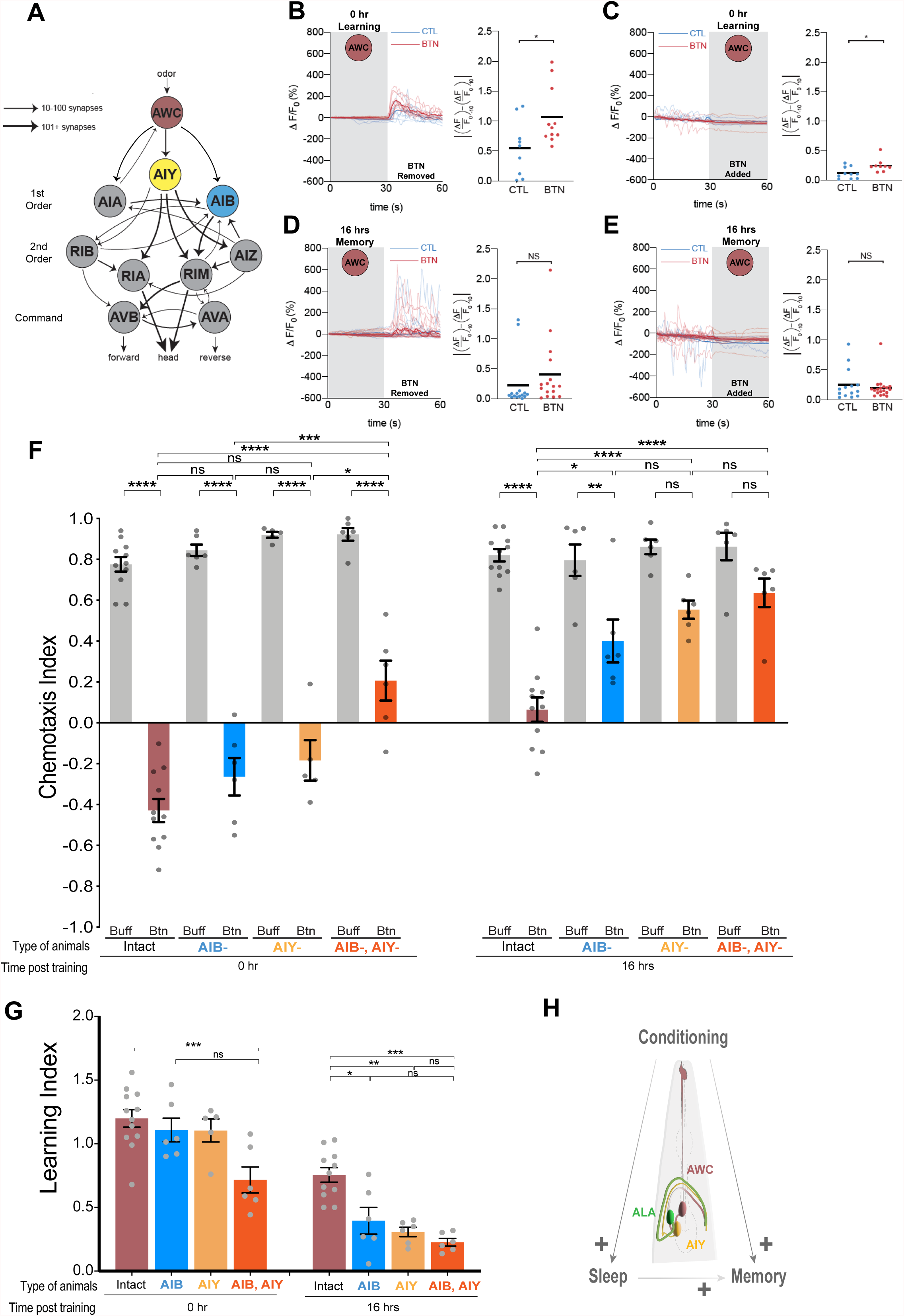
The interneuron AIY is required for sleep-dependent memory. **(A)** The AWC olfactory circuit. Sensory neurons AWC (red) are inhibited by odor. One of the AWC pair is shown. The AIY (yellow) interneuron promotes straight runs and is inhibited by glutamate release from AWC. The AIB (blue) interneuron promotes turns and is activated by glutamate release from AWC. Thus, odor activates AIY and inhibits AIB thereby allowing the animal to run up an odor gradient and reorient if going down the gradient. Smallest arrow indicates 1- 10 synapses between the two neurons, medium arrow, 10-100 synapses, and largest arrow, more than 100 synapses. Only chemical synapses are indicated in the circuit (gap junctions not shown). Figure adapted from Gordus et al., 2015. **(B-E)** Calcium transients (GCaMP3) in the AWC^ON^ of a trapped animal as it is exposed to butanone (grey shaded area) or buffer (white). Blue traces are transients in control trained animals and red those of butanone trained. (**B** and **C**) Transients measured immediately after training or (**D** and **E**) or after 16 hr of recovery on food. The scatter plot panels on the right of each calcium recording show the change in fluorescence immediately before and after the change in stimulus and each point signifies one worm. Paired T-test was performed on all the comparisons. **(F)** The CIs of animals missing: no neurons (brick), AIB (blue), AIY (yellow) or both AIB and AIY (orange) immediately after training (t=0) or after 16 hours of recovery on food. One-way ANOVA with Bonferroni’s correction was performed (****P < 0.0001, ***P < 0.001, ** P < 0.01, *P < 0.05, and (ns) is P > 0.05). N > 5 trials. **(G)** The LIs of animals missing: no neurons (brick), AIB (blue), AIY (yellow) or both AIB and AIY (orange) immediately after training (t=0) or after 16 hours of recovery on food. One-way ANOVA with Bonferroni’s correction (****P < 0.0001, ***P < 0.001, ** P < 0.01, *P < 0.05, and (ns) is P > 0.05). N > 5 trials. **(H)** Model: Spaced olfactory conditioning induces sleep and memory in *C. elegans*. Sleep induced by butanone conditioning is ALA-dependent and benefits memory retention. The signal that butanone has been sensed passes from the AWC neuron to interneurons including AIY and AIB. Memory requires AIY thus we hypothesize that sleep may act on the AIY neuron or the connection between AWC and this interneuron.

AWC calcium levels decrease when animals are exposed to butanone, rise after odor removal, then return to baseline (Chalasani *et al*., 2007; Cho *et al*., 2016). We find that odor removal triggers a small but significantly higher increase in calcium in the AWC neuron immediately after three cycles of butanone-training, compared with animals that were tested immediately after three cycles of buffer-training (Figure 5B, P=0.0264). Likewise, odor onset triggers a small, but significantly greater (Figure 5C, P=0.0172) silencing of the AWC neurons in butanone-trained animals as compared to the buffer-trained controls. The difference between AWC activity in buffer- and butanone-trained animals is thus seen immediately after training while the animals are repulsed by the odor (Figure 5B, C). However, after 16 hours on food, these differences in the AWC response to butanone disappear [Figure 5D (P=0.316) and E (P=0.521)]. Thus, it is unlikely that a change in the sensory response of AWCs is responsible for the long-term memory.

### The long-lasting memory requires the AIY postsynaptic interneurons

We reasoned that the cells responsible for memory might be downstream of AWCs in the chemosensory circuit. Serial electron micrographs (Cook et al., 2019; White *et al*., 1986; Witvliet et al., 2020) indicate that the AWC chemosensory neuron pair forms synapses with three pairs of interneurons, the AIY interneurons (25-34 synapses), the AIB interneurons (22-29 synapses), and the AIA interneurons (22 synapses) (Figure 5A). Since AIA is required for butanone learning after one cycle of training (Cho *et al*., 2016) the learning defects of AIA-ablated strains (C.B. and K.B. personal communication) were not assessed. To inactivate AIY neurons, we employed the *ttx-3(ks5)* mutant allele, which prevents the birth of the AIY neurons (Altun-Gultekin et al., 2001). To kill AIB neurons, we expressed the caspase CED-3 from the *odr-2b* promoter, which is specific for AIBs and kills them during development (Chelur and Chalfie, 2007; Chou et al., 2001).

We found that animals that lack either AIY or AIB neurons exhibit normal chemotaxis and learning (Figure 5F and G, compare the CIs and LIs of brick (intact), yellow (AIY-) and blue (AIB-) at 0 hours after training). However, when AIBs and AIYs are both missing, this learning is reduced from that of wild type (see Figure 5F and G, compare first [brick, intact] and fourth [green, double ablation] pair of bars). This might be explained if another neuron in the circuit is primarily responsible for learning and AIBs and AIYs are redundant or play a smaller role.

16 hours after training, animals lacking AIB neurons are able to retain some memory, but animals missing AIY neurons do not exhibit any significant difference between buffer- and butanone-trained chemotaxis indices (Figure 5F). This suggest that AIY interneurons are required to retain memory. Learning indices of animals at this time indicate that both AIB and AIY interneurons are required for long-term memory, but loss of AIBs leads to more variable deficits (Figure 5G). Animals deficient in both AIB and AIY neurons learn less at 0 hours and retain the least memory at 16 hours (see Figure 5F and G). We observed variable CIs of butanone-trained populations when animals lack either AIBs, AIYs, or both AIBs and AIYs (Figure 5F), thus we calculated the degree of memory loss occurring in each cohort to account for the variability (Figure S3). We found that when animals lack AIY interneurons, the learning indices between 0 hour and 16 hours shows the biggest depreciation of memory retained with least variability (Figure S4C). Thus, we focused on understanding if the mechanism by which long-term memory depends on the interaction between the AWC and AIY neurons (Figure 5H).

### AWC-AIY synapses are visualized with NLG-1 GRASP

To understand the mechanism by which memory is stored, we sought to understand if olfactory synapses are altered in animals that remember their training. We focused on synaptic connections between the AWC chemosensory neurons and the AIY interneurons, as AIY neurons are consistently required for the olfactory memory (Figure 5F and G), and AWCs form the largest number of synapses with AIY neurons (White *et al*., 1986; Witvliet *et al*., 2021). To visualize AWC-AIY synapses, we utilized Neuroligin 1 GFP Reconstituted Across Synaptic Partners (NLG-1 GRASP), a split GFP-based trans-synaptic marker [Figure 6A (Feinberg et al., 2008; Park et al., 2011; Varshney et al., 2018)]. The marker has two complementary GFP fragments, GFP1-10 and GFP11, which can reconstitute and fluoresce when they come in contact (Cabantous et al., 2005). The split GFP fragments are connected via flexible linkers to the transmembrane synaptic protein Neuroligin-1 (NLG-1), which localizes to pre- and postsynaptic sites in *C. elegans* (Feinberg *et al*., 2008).

**Figure 6.**
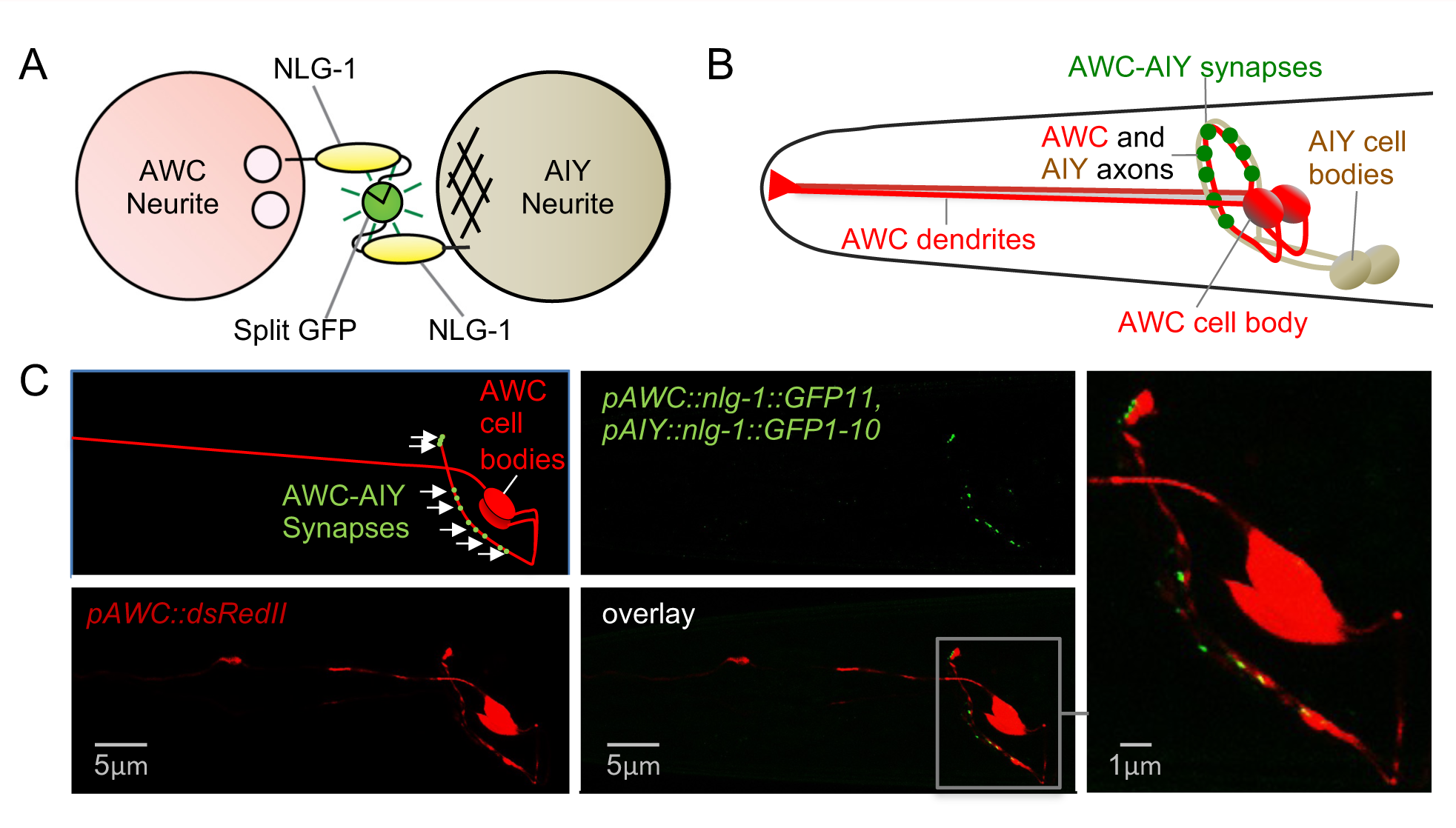
NLG-1 GRASP Visualizes synapses between AWC chemosensory neurons and AIY interneurons. **(A)** Schematic of split GFP-based AWC-AIY NLG-1 GRASP marker. Circles represent cross-sections of the AWC and AIY neurites, and one neurite from each neuron pair is represented for simplicity. Fragments of the split GFP are linked to the pre- and postsynaptically localized protein NLG-1 (Neuroligin 1), and expressed in the AWC and AIY neurons with the selective promoters *_p_odr-1* and *_p_ttx-3*. When synapses form between the neurons, the split GFPs come in contact, reconstitute and fluoresce. Small white circles indicate a presynaptic site, and crosshatching represents a postsynaptic site. **(B)** Schematic of the head of an animal in which NLG-1 GRASP labels synapses between the AWC (red) and AIY (beige) neurites in the nerve ring, which forms an arch in the head of the animal. **(C)** Schematic and micrographs of an animal carrying the AWC-AIY NLG-1 GRASP marker with the AWC neurons labeled in red with the cytosolic mCherry fluorophore. Synaptic fluorescence is observed in a punctate pattern in AWC axons in the nerve ring. The area in the gray box is expanded in the rightmost image.

To visualize AWC-AIY synapses, we generated a construct driving expression of NLG-1::GFP11 in AWC neurons and coinjected it with a construct driving expression of NLG-1::GFP1-10 in AIY neurons. An additional construct drove expression of cytosolic dsREDII in the AWC neurons (Feinberg *et al*., 2008; L’Etoile and Bargmann, 2000) to visualize AWC neurites. We generated transgenic animals carrying these markers, integrated the marker into the genome, and outcrossed background mutations. AWC neurons have dendrites that extend to the nose of the worm, and axons that extend into the nerve ring, which forms an arc in the head of the worm [Figure 6B, (White *et al*., 1986)].

Electron micrograph reconstruction studies indicate that AWC neurons form *en passant* synapses onto the left and right AIY neurons in the nerve ring (White *et al*., 1986; Witvliet *et al*., 2021). We found that AWC-AIY NLG-1 GRASP labeling results in fluorescent green puncta along the AWC axons in the nerve ring (Figure 6B and C). The localization and distribution of AWC-AIY NLG-1 GRASP fluorescent puncta along the nerve ring in the head (Figure 6C) was consistent with previous electron micrograph reconstructions, as has been the case for several other neurons throughout the animal that have been visualized with NLG-1 GRASP (Cook *et al*., 2019; Feinberg *et al*., 2008; Park *et al*., 2011; Varshney *et al*., 2018; White *et al*., 1986; Witvliet *et al*., 2021). NLG-1 GRASP does not indicated directionality of synapses, however EM studies indicate that AWC chemosensory neurons are presynaptic to AIY interneurons (https://nemanode.org).

### AWC-AIY synapses are reduced in animals with the olfactory memory

To determine if AWC-AIY synapses are physically altered in animals that retain the olfactory memory, we compared AWC-AIY synapses in populations of odor-trained animals with the olfactory memory to populations of buffer-trained control animals. Animals carrying the AWC-AIY NLG-1 GRASP marker were trained for three cycles with either butanone or a control buffer (as in Figure 1A). Their synapses were imaged 16 hours after training was completed. Interestingly, we found that AWC-AIY NLG-1 GRASP fluorescence intensity was significantly reduced in populations of butanone-trained animals that held the olfactory memory, when compared with populations trained with the control buffer that were attracted to butanone (Figure 7A, 6B [left two micrographs], S5A, and S5B). We quantified AWC- AIY NLG-1 GRASP intensity in these populations, and found that the synaptic signal in butanone-trained animals was significantly lower than in animals trained with a control buffer (Figure 7C, 7D [left two boxes], S5A, and S5B). These results indicate that training with butanone results in a synaptic reduction between chemosensory neurons and the postsynaptic cells required for olfactory memory. We assessed AWC-AIY synapses in animals immediately before training, and found that levels were not significantly different from buffer-trained animals 16 hours after training (Figure S5C), consistent with synapses being reduced in butanone-trained animals.

**Figure 7.**
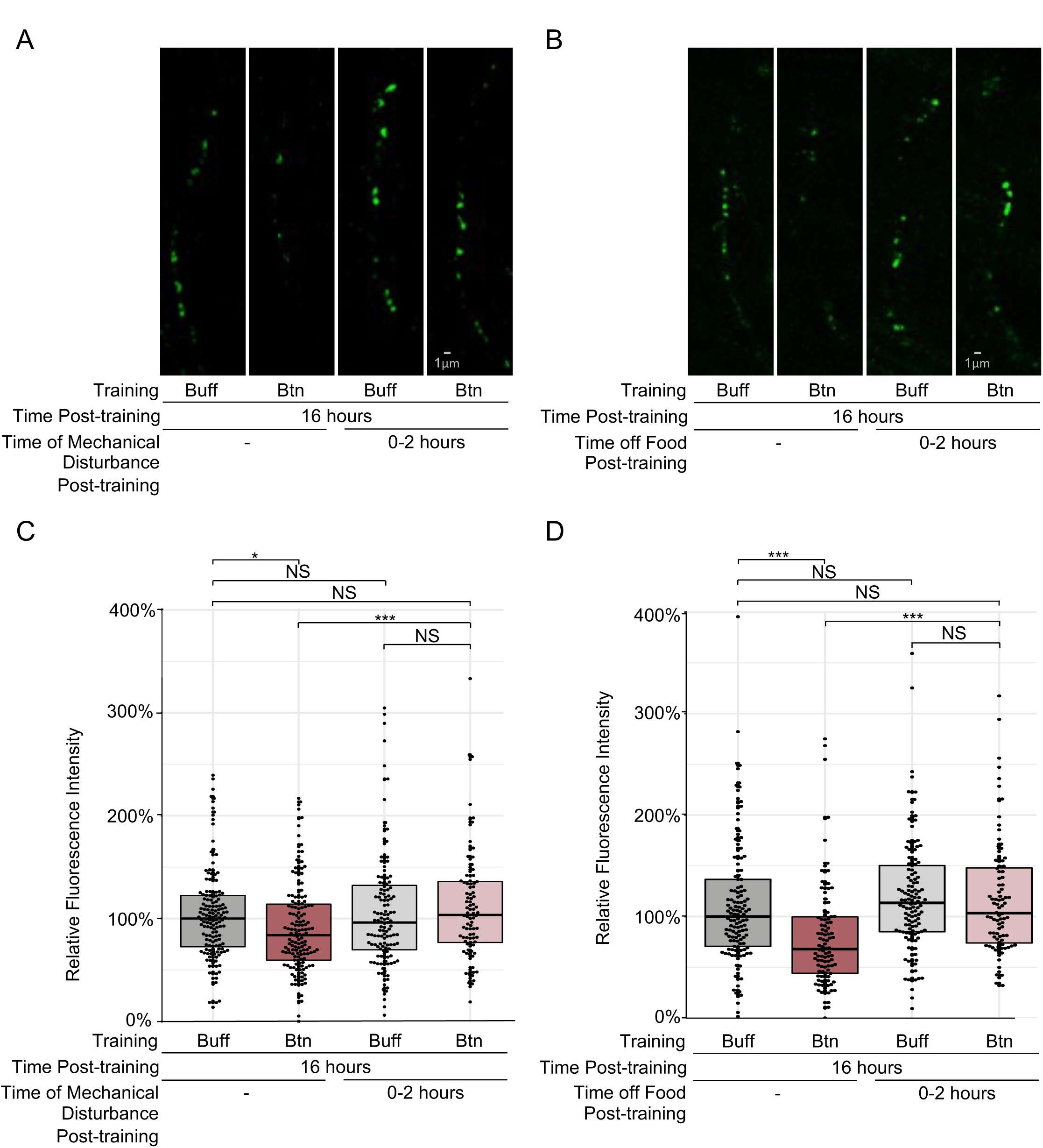
Odor training and sleep result in AWC-AIY synaptic reductions. **(A)** Micrographs of AWC-AIY NLG-1 GRASP fluorescence 16 hours after training with control buffer (Buff) or butanone (Btn) in which sleep was not disrupted after training (left two micrographs), and in which sleep was disrupted by mechanical disturbance for the first two hours after training (right two micrographs). **(B)** Micrographs of AWC-AIY NLG-1 GRASP fluorescence 16 hours after training with control buffer or butanone in which sleep was not disrupted after training (left two micrographs), and in which sleep was disrupted by removal from food for the first two hours after training (right two micrographs). **(C)** Quantification of the reduction in AWC-AIY NLG-1 GRASP fluorescence intensity in animals trained with butanone in which sleep was not disrupted, in comparison with animals whose sleep was disrupted by mechanical disturbance and animals trained with control buffer. n>90 for each box and includes animals trained on four different days. NS P>0.05, * P<0.05, *** P<0.001, Mann-Whitney U-test. P-values were adjusted for multiple comparisons using the Hochberg procedure. **(D)** Quantification of the reduction in AWC-AIY NLG-1 GRASP fluorescence intensity in animals trained with butanone in which sleep was not disrupted, in comparison with animals whose sleep was disrupted by removal from food and animals trained with control buffer. n>90 for each box and includes animals trained on four different days. NS P>0.05, *** P<0.001, Mann-Whitney U-test. P-values were adjusted for multiple comparisons using the Hochberg procedure.

### Sleep is required for AWC-AIY synaptic reductions after odor training

We asked whether the AWC-AIY olfactory synaptic reductions in populations of butanone-trained animals were dependent on the two-hour period of post-training sleep required for olfactory memory. To test if the critical period of sleep was required for synaptic changes observed 16 hours after training, we disrupted sleep by either shaking the worm plates every 15 minutes or removing the animals from food (as in Figure 4) for two hours immediately after training. Animals deprived of sleep for the first two hours by either method were then moved to food plates for 14 hours before synapses were assessed. We found that in populations of animals whose sleep was disrupted during the critical period and whose olfactory memory was perturbed, the synaptic reduction was absent (Figure 7A-D, S5A and S5B).

Specifically, populations of butanone-trained animals deprived of sleep during the critical period (by either method) that lost the olfactory memory had significantly higher NLG-1 GRASP fluorescence intensity than butanone-trained animals that were allowed to sleep and retained the olfactory memory (Figure 7C, 7D, S5A and S5B). Similarly, the NLG-1 GRASP fluorescence intensity in sleep-deprived animals who lost the memory was not significantly different from that of buffer-trained control animals (Figure 7C, 7D, S5A and S5B). These data indicate that the critical period of sleep for olfactory memory is also required for butanone training-induced AWC-AIY synaptic reductions that correlate with memory. Furthermore, 16 hours after training, synapse levels correlate with behavioral responses: lower synaptic levels are found in populations with weaker attraction to butanone, while higher synaptic levels of AWC- AIY synapses are found in populations with a stronger attraction to the odor.

To determine if synaptic changes in response to butanone training are global, or restricted to the butanone chemosensory circuit, we examined the synaptic connections between PHB chemosensory neurons and two of their primary postsynaptic partners, the AVA neurons, using a strain that carries a NLG-1 GRASP marker that labels connections between this pair of neurons (Park *et al*., 2011; Varshney *et al*., 2018). PHB chemosensory neurons sense noxious chemicals, including dodecanoic acid (Tran et al., 2017) and sodium dodecyl sulfate (Hilliard et al., 2002). We treated and selected populations of PHB-AVA NLG-1 GRASP-labeled animals similarly to the populations of AWC-AIY NLG- 1 GRASP-labeled animals above, and found that PHB-AVA connections are not significantly altered by butanone-training or sleep (Figure S6). This indicates that the synaptic changes induced by butanone training and sleep are not global.

### Two temporally distinct processes affect AWC-AIY synapses after training

To understand whether AWC-AIY synapses change during the critical period, we imaged single animals carrying the AWC-AIY NLG-1 GRASP marker immediately after odor training and again after the critical period of sleep. As with most fluorescent synaptic markers, NLG-1 GRASP undergoes photobleaching during imaging, however this should be consistent between buffer- and butanone-trained animals.

Therefore, rather than assess the percent fluorescence intensity reduction in each animal, we determined the proportion of animals with large (≥ 50%) reductions between two time points. Individual animals were imaged from populations of buffer-trained animals that chemotaxed to butanone (CI>0.5) and populations of butanone-trained animals that did not chemotaxis to butanone (CI<0.5) at 0 and 2 hours. During the critical period of sleep (between 0 and 2 hours after training), similar proportions of buffer- and butanone-trained animals had a large reduction in synaptic intensity; 64% of butanone-trained animals and 52% of buffer-trained animals had large synaptic reductions during this period, and these proportions were not significantly different (P=0.21, two-independent sample z-test) (Figure S7A- E). This suggests that synapses are reduced after training independently of whether animals are exposed to odor.

To determine if synapses change during the 14 hours after the critical period, we imaged individual animals from populations of buffer-trained animals that chemotaxed to butanone (CI>0.5) and populations of butanone-trained animals that did not chemotaxis to butanone (CI<0.5) at 2 and 16 hours. We imaged individual animals two hours after training, then at 16 hours, tested each animal for chemotaxis to butanone. Individual buffer-trained animals that were attracted to the odor, and butanone-trained animals that were not attracted to the odor were then imaged again. For animals to pass this behavioral screen, buffer-trained worms needed to move directly towards butanone or stay on the butanone side of the plate the majority of the time, while butanone-trained worms needed to move and not chemotax towards butanone or spend the majority of time on the butanone side of the plate.

Between 2 and 16 hours, 57% of butanone-trained animals had a large synaptic reduction compared to 33% of buffer-trained animals, although these proportions were not significantly different (p=0.061, two-independent sample z-test) (Figure S7A-E).

To further understand how olfactory synapses change over the time course of memory consolidation, we examined AWC-AIY synapses in populations of animals after training. The population assays allowed for larger sample sizes and avoided the issue of photobleaching. We assessed AWC- AIY synapses in butanone-trained populations that chemotaxed to butanone (CI>0.5) and buffer-trained populations that did not chemotax to butanone (CI<0.5) at 0, 2 and 16 hours after training (Figure S7F). We found that immediately after training, both control buffer- and butanone-trained animals begin with higher levels of synapses than those in control buffer-trained animals 16 hours post-training (Figure 8A and B). This indicates that training with butanone alone does not instantly alter synaptic structures.

**Figure 8.**
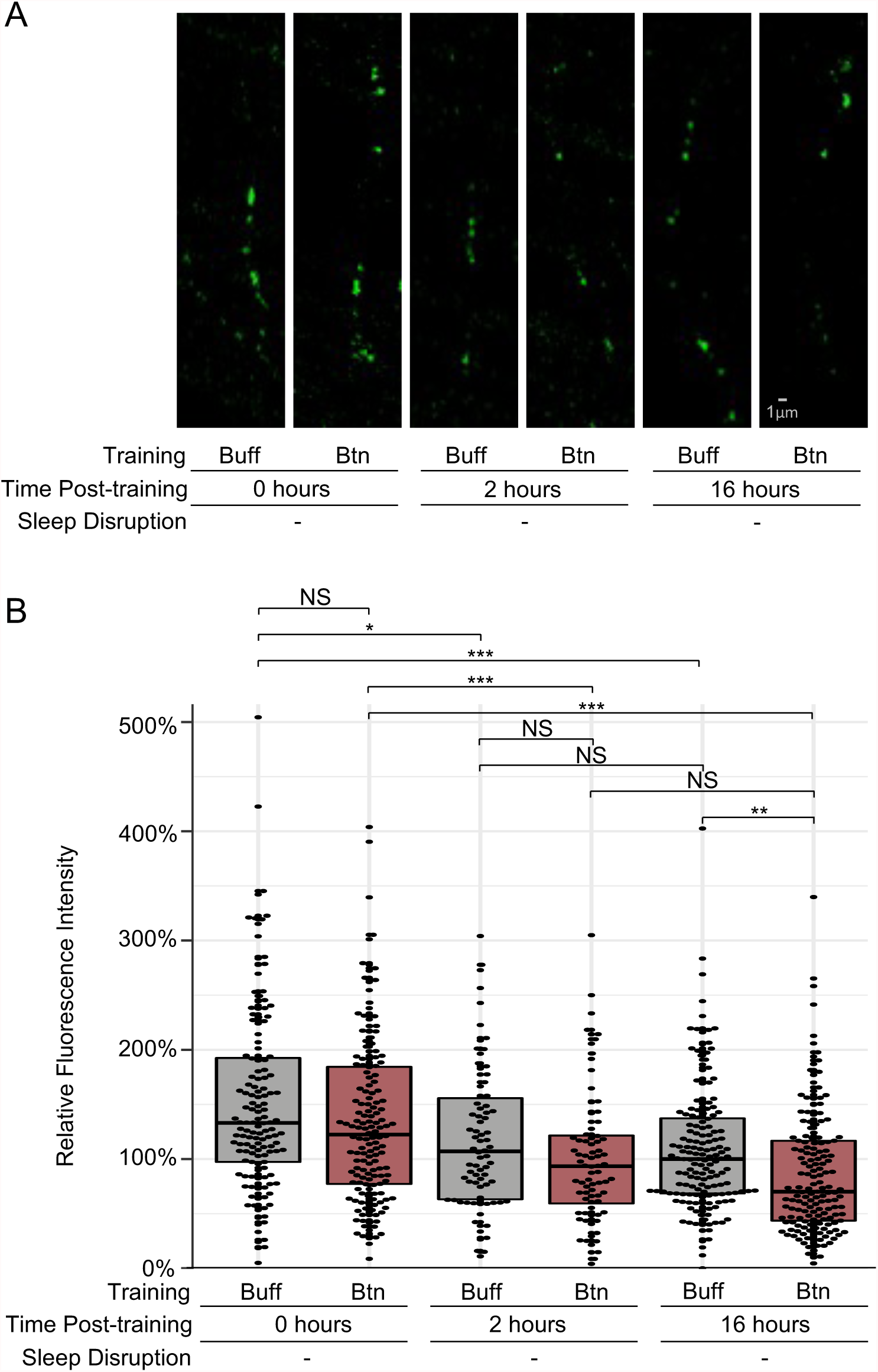
AWC-AIY synapses are altered during and after sleep. **(A)** Micrographs of AWC-AIY NLG-1 GRASP fluorescence in animals trained with butanone (Btn) or control buffer (Buff) at 0 hours, 2 hours, and 16 hours after training without sleep disruption. **(B)** Quantification of AWC-AIY NLG-1 GRASP fluorescence intensity at 0 hours, 2 hours and 16 hours post-training in buffer-trained and butanone-trained animals whose sleep was not disrupted after training. N>75 for each box and includes animals trained on four different days. NS P>0.05, * P<0.05, ** P<0.01 and *** P<0.001, Mann-Whitney U-test. P-values were adjusted for multiple comparisons using the Hochberg procedure.

However, during the two-hour critical period of sleep after training, synapses in both buffer- and butanone-trained worms are similarly reduced (Figure 8A and B). Consistent with our observations in single animals, this indicates that during the first two hours after training, synapses are reduced in an odor training-independent manner.

By 16 hours after training, we observed a significant reduction of AWC-AIY synapses when compared with control buffer-trained animals (Figure 8A and B), similar to that observed in previous assays (Figure 7). This indicates that although synaptic levels are similar in buffer- and butanone-trained animals two hours after training, they become distinct in the two populations after this critical period of sleep. Thus, there are actually two phases of synaptic changes after training: an odor training-independent synaptic reduction during the first two hours post-training, and a process by which synaptic levels in butanone-trained animals become lower than those in buffer-trained animals that takes place during the following 14 hours. Though this second phase of synaptic reduction requires sleep after training (Figure 8A and B), structural changes are not complete until after the critical period.

## DISCUSSION

Sleep is highly conserved, which indicates an evolutionary pressure to retain this mysterious state (Cirelli and Tononi, 2021). It is reasonable that long-term memory is also selected for, as individuals that fail to learn from experience are at a disadvantage (Dissel, 2020; Ganguly-Fitzgerald *et al*., 2006; Glou et al., 2012). The linkage between the processes of sleep and memory has traditionally been investigated using organisms with complex nervous systems containing more than 100,000 neurons. This complexity has hampered understanding how the connections between synaptic partners that are required for memory are affected by sleep. We discovered that sleep is required for long-term CREB-dependent olfactory memory in *C. elegans*. By exploiting the simplicity of the *C. elegans* nervous system, we identified a synaptic partner pair that is required for memory and examined the connections between them as a function of sleep. We show that the structural connections between these two cells are reduced when the animals consolidate memory. This synaptic reduction and long-term memory requires sleep immediately after training. It is surprising that *C. elegans*, which has one of the simplest nervous systems of any metazoan with only 302 neurons, also requires sleep to both consolidate memory and modulate synapses. This suggests that the role of sleep in memory and synaptic modulation is conserved in the vast majority of metazoan species on earth, and is required even in the most compact nervous systems.

### Odor-dependent and independent synaptic modulation during sleep

The odor-independent synaptic reductions that we observe in the first two hours after training are reminiscent of synaptic downscaling seen in many vertebrate systems when the organism sleeps. The synaptic homeostasis hypothesis (SHY) states that the brain resets during sleep by reducing global synaptic strength (Tononi and Cirelli, 2014). Indeed, broad reductions in synaptic strength have been reported in many brain regions during sleep, although studies have also demonstrated widespread increases in synaptic strength in some regions (Durkin and Aton, 2016). Our paradigm may reveal that sleep in *C. elegans* conforms to the synaptic homeostasis hypothesis, as synapses are reduced in the first two hours of sleep after training, regardless of whether animals were trained with odor or butanone. Thus, the reductions occur during sleep, but are not dependent on olfactory experience in the first two hours of sleep.

Sleeping animals show limited neural activity throughout their anterior neuropil (Nichols *et al*., 2017; Skora *et al*., 2018) and this low level of activity may permit the synaptic reductions that we observe. Indeed, synaptic transmission at GABAergic neuromuscular junctions decreased in animals sleeping during developmental lethargus, but UNC-49 GABA receptor immunostaining was not reduced (Dabbish and Raizen, 2011). Studies of unrestrained, sleeping animals showed that the neural dynamics of the AVA backward command neuron is severely blunted, while AWA, an appetitive sensory neuron that has an ON response, rather than an OFF response like AWCs, has a prolonged response to odor when sleeping (Lawler et al., 2021). A similar analysis of the AWC sensory circuit during post-training sleep is required to understand the neural dynamics in the sleeping worm, so that the relationship between neuronal activity and synapse size can be determined.

The odor-dependent synaptic reduction we have observed 16 hours after training shares characteristics with synaptic consolidation reported in vertebrates (Havekes and Abel, 2017). Synaptic consolidation involves the transition from modulation of synaptic strength immediately after learning to more permanent changes in synaptic structures associated with long-term memory. Likewise, the differences in synaptic structures seen 16 hours after training may be preceded by modulation of synaptic strength immediately after training. Our work indicates that a tight temporal link between odor training and sleep is critical for memory. Similarly, the synaptic reduction seen in butanone-trained animals 16 hours after training required two hours of sleep immediately after training. This suggests that odor training-induced changes that mark synapses for reduction are immediately acted on by sleep to promote long-lasting changes. A temporal link between training and sleep has also been demonstrated to be important for memory in vertebrates. For example, a specific three-hour period of sleep is required for some forms of hippocampal synaptic plasticity and memory (Prince et al., 2014).

### AWC-AIY synaptic reductions could contribute to odor memory

When a worm forages on a Petri dish or in the environment, it experiences changes in the level of butanone odor and food odor. These changes resemble removal from butanone or food, which causes calcium increases and presynaptic vesicle release for AWC neurons (Chalasani *et al*., 2007; Cho *et al*., 2016). However, when animals are trained in our paradigm, they are immersed in liquid with butanone without food, which is a condition that reduces calcium influxes in AWCs [Figure 4 and (Cho *et al*., 2016)], and likely results in far less synaptic release. This quiet state followed by post-training sleep may lead to the reduction of synaptic structures. While learning resulting from synaptic increases has been studied far more, learning resulting from synaptic reductions has also been documented (Collingridge et al., 2010). An interesting parallel is the involvement of LTD in some forms of extinction, in which animals are trained to forget a learned behavior (Collingridge *et al*., 2010). Although our paradigm trains animals to stop performing an innate behavior, which is more similar to inhibitory operant conditioning, there may be similarity between the mechanisms.

It is tempting to speculate that reduction of synapses may be an important component of learning and remembering many motor coordination tasks, in which it is as important to not contract unnecessary muscles as to contract the correct ones. Such synaptic reductions might be underappreciated if they are not detected using common methods for visualizing memory engrams, as these are usually associated with building synapses. Further, our findings suggest that in addition to the protective effects of sleep on synapses that have been used more during wake (Cirelli and Tononi, 2021), there may be an increased or extended effect of sleep on synapses that have been used less during wake. This would be consistent with the extended period of synaptic reduction observed in butanone-trained animals allowed to sleep after training.

### Cellular loci for olfactory learning and memory

Here we show that learning to avoid butanone requires either the AIY or AIB interneurons, and the involvement of AIA interneurons has been documented previously (Cho *et al*., 2016). We further show that AIY, and to a lesser extent, AIB interneurons are required for sleep-dependent long-term memory. AIYs were also found to be critical for *C. elegans* to learn to avoid the pathogenic bacteria *Pseudomonas aeruginosa* (PA14) (Zhang et al., 2005). *Pseudomonas aeruginosa* emits butanone in a complex mixture of volatiles (Labows *et al*., 1980) which are the cues by which *C. elegans* decides to avoid or seek out bacteria (Worthy *et al*., 2018a; Worthy *et al*., 2018b; Zhang *et al*., 2005). Their innate attraction to PA14, like that of butanone, can be changed to aversion by PA14 exposures lasting four hours (Ha and O’Toole, 2015; Liu et al., 2022; Zhang *et al*., 2005). Ha et al., 2015 and Liu et al., 2022 showed that the switch from attraction to repulsion correlates with a decrease in the size of the PA14- evoked calcium transients in AIYs. The loss of responsiveness may result from a higher baseline of calcium activity (Liu *et al*., 2022). As AWCs release glutamate, which inhibits AIY neurons via glutamate-gated chloride channels (Chalasani *et al*., 2007), the higher AIY baseline in PA14-trained animals could result from reduced inhibition from AWCs. One way this might occur is if the synapses between AWC and AIY neurons are reduced after PA14 training, similarly to our observation in butanone-trained animals. AIYs receive input from many other neurons and the changes in calcium transients may thus reflect more than just AWC inputs. Still, these findings are consistent with both training paradigms reducing transmission between AWC and AIY neurons.

Our butanone training paradigm does not affect chemotaxis to odors sensed other chemosensory neurons. Since first-order interneurons in the olfactory circuit have inputs from multiple chemosensory neurons, we focused on connections whose modulation would not affect AWA-mediated chemotaxis. However, several pairs of second-order and command interneurons have been implicated in other learning and memory paradigms in *C. elegans.* A pair of downstream interneurons, the RIAs, plays a role in PA14 learning (Ha and O’Toole, 2015; Liu *et al*., 2022). Previous butanone learning paradigms (Lakhina et al., 2015) also found that CREB was required for long-term memory and its expression changed most in another downstream interneuron pair, the AIMs. Studies of the AWA chemosensory circuit in diacetyl olfactory learning also showed increases in the postsynaptic regions of a downstream pair of backward command interneurons, the AVAs (Hadziselimovic et al., 2014). This indicates that the memory of different forms of training might involve changes in both upstream and downstream neurons in the olfactory circuits, and that different forms of training may cause distinct circuit changes.

### The *C. elegans* olfactory circuit depends on sleep for memory

The requirement for sleep to consolidate memory may depend on the circuits that store the memory. Chouhan and colleagues showed that flies that are starved after appetitive training do not need to sleep to consolidate the memory of the appetitive odor (Chouhan et al., 2021). This is distinct from flies that are fed directly after training, as they require sleep to consolidate memory after the same training paradigm. The authors show that starved flies utilize a circuit that does not require sleep for activity. By contrast, fed flies use a circuit that is both active in sleep and promotes sleep after training and feeding. This indicates that perhaps circuits that require sleep for memory consolidation are active during sleep. Chouhan and colleagues further show that feeding and starvation use the feeding-related neuropeptide F to toggle between sleep-dependent and sleep-independent circuits. This study demonstrates that long-term butanone memory requires sleep whether the animals are fed or removed from food after training (Figure 4). The receptor for the *C. elegans* neuropeptide F homolog, neuropeptide Y, is NPR-1. Previous work indicates that NPR-1 is biased to its active state by a mutation in the G alpha binding loop (215V) in the wild type strain we use (N2) (De Bono and Bargmann, 1998) and this may restrict memory formation to a feeding and sleep-dependent circuit.

### Decay of long-term olfactory memory

Many studies demonstrate that forgetting pathways can be engaged after learning in *C. elegans*, so that the memory of their training is significantly reduced within two hours (Hadziselimovic *et al*., 2014; Inoue et al., 2013; Liu *et al*., 2022). For example, olfactory training with diacetyl confers a short-lived increase in GLR-1/GluR1-labeled synapses onto the backward command interneurons in the worm, the AVAs. The memory and size of these synapses decayed within two hours (Hadziselimovic *et al*., 2014), however, the role of sleep in diacetyl memory decay has not been examined. Memory in the butanone learning paradigm described in this work is kept for at least 24 hours if the animals are allowed to sleep after training, although the learning index decreases. These pathways may not be engaged after butanone learning, since the memory does not decay as quickly. The impact of sleep on forgetting pathways will be an interesting avenue for future studies.

Learning to overcome an animal’s innate attraction to butanone may resemble inhibitory operant conditioned memory, in which the strength of a voluntary behavior is modified by punishment. This associative learning task requires the animals to associate an innately attractive odor with the absence of food. Inhibitory operant conditioned memory has been shown to require sleep in invertebrates and vertebrates (Chouhan *et al*., 2021; Rasch and Born, 2013). Interestingly, the extinction of inhibitory operant conditioned memory can be promoted by wakefulness in the presence of the unconditioned stimulus, rather than sleep (Vorster and Born, 2017). Thus, it is possible that if worms were removed from food (unconditioned stimulus) for the first two hours after odor conditioning, it could result in extinction of the butanone memory. However, we found that animals that were maintained on food after conditioning, and whose sleep was mechanically disrupted so that they were not re-exposed to the unconditioned stimulus, also lost the memory 16 hours after training. This indicates that the loss of the butanone long-term memory was due to loss of sleep, rather than extinction.

### Sleep in the butanone conditioning paradigm

This training paradigm, which involves animals swimming for 300 minutes total before being placed onto a solid substrate with food, likely induces sleep as a consequence of mobilization of fat stores after exercise (Grubbs *et al*., 2020), and could also involve satiety-induced quiescence (Gallagher and You, 2014; You *et al*., 2008). While investigating whether the known sleep-promoting peptides and cells are involved, we found that inactivation of the ALA but not RIS impaired memory consolidation (Figure 2).

Different types of sleep (NREM and REM) may be required for different types of memory (Barnes and Wilson, 2014; MacDonald and Cote, 2021). For a similar understanding of sleep architecture in *C. elegans*, it will be useful to observe both muscle and neuronal activity in animals that show the hallmarks of sleep. Lawler and colleagues have begun these studies and verified that sleeping worms show a more flaccid posture, though they twitch the anterior portions of their body during bouts of spontaneous sleep (Lawler *et al*., 2021). Detailed analysis of calcium currents in post-training sleep may allow us to better understand sleep architecture in memory consolidating animals.

## Supplemental Figures, Tables and Videos

**Supplemental Figure 1:**
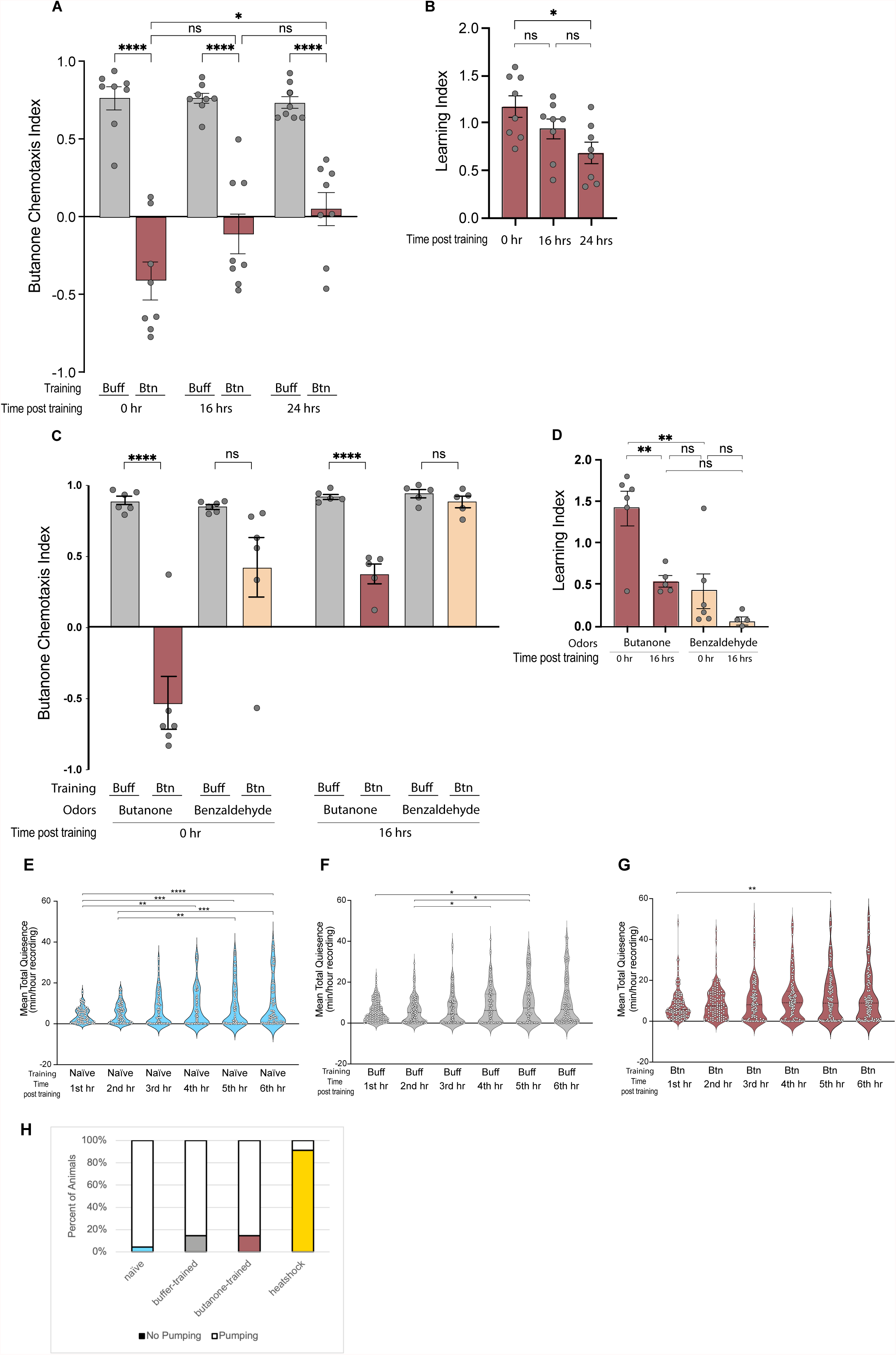
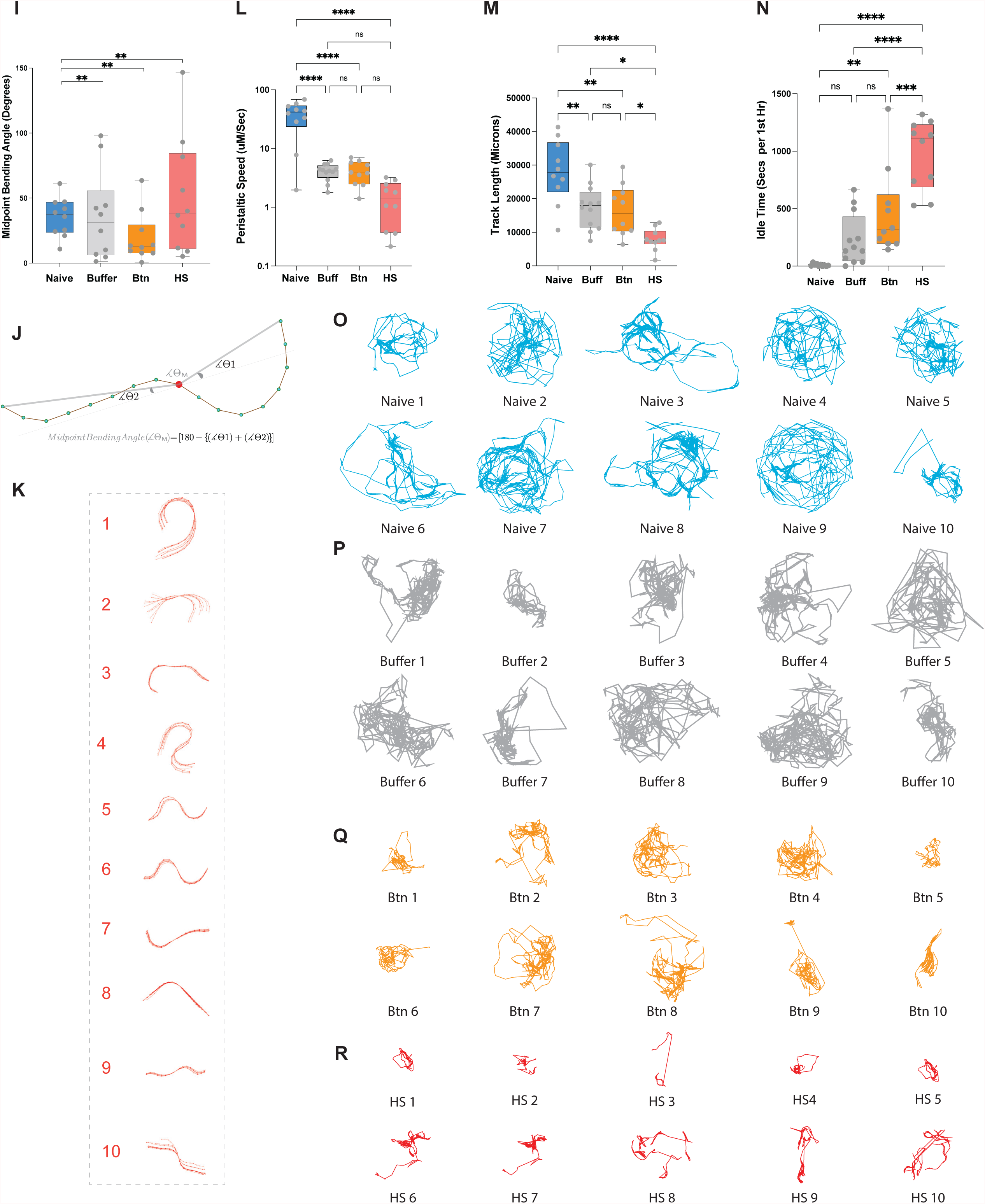
Supports Figure 1. (A) The CIs and (B) the LIs of buffer and butanone trained animals show that memory persists up to 24 hours. One-way ANOVA was performed to compare LIs wherein the P values are reported as ****P < 0.0001, ***P < 0.001, ** P < 0.01, *P < 0.05, and (ns) is P > 0.05. N > 7 trials. (C) and (D) Three cycle butanone training does not affect attraction to benzaldehyde after 16 hours of recovery (showing specificity of butanone sensing). CIs of animals trained to buffer or butanone are tested for attraction to butanone or benzaldehyde at 0 hour and 16 hours The u-test was performed on Buff vs Btn 0 hr and 16 hours before using student’s t-test to confirm the differences in CIs. P values are reported as ****P < 0.0001, ***P < 0.001, ** P < 0.01, *P < 0.05, and (ns) is P > 0.05. N = 5 trials. The LIs show that animals specifically remember to avoid butanone rather than benzaldehyde after butanone conditioning. One-way ANOVA with Bonferroni’s correction (****P < 0.0001, ***P < 0.001, ** P < 0.01, *P < 0.05, and (ns) is P > 0.05). N =5 trials (E) (F) and (G) The analyses of six-hour sleep show naïve and buffer trained animals become quiescent after staying for at least three hours in WorMotel. This suggests that prolonged WorMotel stay induces sleep. However, the butanone trained animals show least increase in the amount of sleep over the course of six hours in WorMotel because they are already sleeping most immediately after training One-way ANOVA was performed, and P values are reported as ****P < 0.0001, ***P < 0.001, ** P < 0.01, *P < 0.05, and (ns) is P > 0.05. Each gray dot represents the number of animals. N =7 trials. (H) With heat shock animals as a positive control, a recovery plate after training contains two kinds of animals pumping (white portion) and no pumping (colored portions). Percent of animals not pumping are significantly higher than naïve. Z-test with Hochberg correction (**P<0.005). (I) With heat shock animals as a positive control, the midpoint bending angle of the trained animals is significantly lower than the untrained animals. One sample t and Wilcoxon signed rank test (**P<0.005), N =10 animals. (J) The equation to measure mid-point bending angle. (K) The midpoint bending angle of the heat shock animals. (L) The peristaltic speed of the trained animals is significantly lower than the untrained animals aand like heat shock animals. One-way ANOVA with Bonferroni’s multiple correction (****P<0.0001), N =10 animals (M) The track length of the heat shock animals is lowest, and the track length of the trained animals is significantly lower than the untrained animals. One-way ANOVA with Bonferroni’s multiple correction (****P<0.0001), N =10 animals (N) The idle time is an independent confirmation of the WormLab software and the matLab script of the WorMotel showing that butanone trained populations sleep more than naïve and the heat shock animals being a positive control for sleep, are most quiescent. One-way ANOVA with Bonferroni’s multiple correction (****P<0.0001), N =10 animals (M) (M) (N) (O) and (P) The track trajectories during 1^st^ hour post training of naïve, buffer, butanone and heat shock animals are shown. The trajectories become small as each animal starts sleeping more.

**Supplemental Figure 2:**
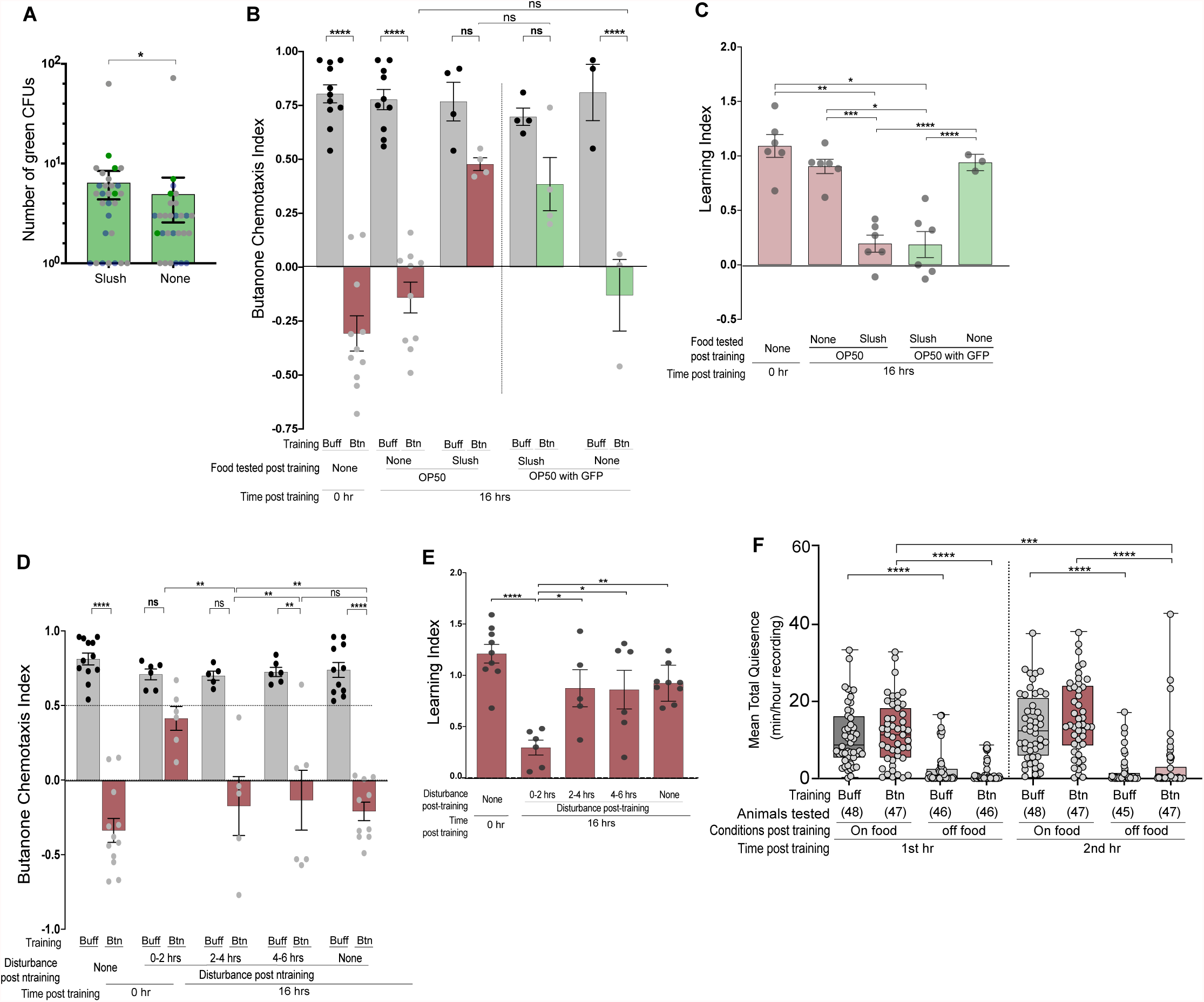
Supports Figure 3. (A) To confirm that animals were eating during mechanical disturbance, we rinsed the worms with 0.5% bleach to kill any OP50 sticking outside the body after sleep disruption, dissected the alimentary canal of the worms, and incubated the alimentary canal for 15 minutes with SOC medium before plating on LB plates to count the colony-forming units (CFU) that were green. As the animals were only fed with green OP50 during the period of mechanical disturbance from 0-2 hours, presence of green CFUs confirmed that the animals ate during the disturbance. Statistical significance is reported as *p<0.05* after paired t-tests. (B and C) The CIs and LIs after recovery with OP50 versus OP50 with GFP show that there no behavioral differences in animals, therefore, the difference in behavior arises due to mechanical disturbances. One-way ANOVA was performed, and P values are reported as ****P < 0.0001, ***P < 0.001, ** P < 0.01, *P < 0.05, and (ns) is P > 0.05. Each gray dot (N) represents the number of independent trials. (D and E) Bar graphs showing the CIs and LIs of animals that were treated with bacterial slush for a two-hour period after training. This data show that changing the food viscosity is sufficient to disrupt sleep and memory. However, to obtain uniformity in sleep disruption, mechanical disturbance in less viscous food was performed (Figure 2B and C). One-way ANOVA was performed, and P values are reported as ****P < 0.0001, ***P < 0.001, ** P < 0.01, *P < 0.05, and (ns) is P > 0.05. Each gray dot (N) represents the number of independent trials. (F) Starvation during the first two hours after training disrupts sleep. One-way ANOVA was performed, and P values are reported as ****P < 0.0001, ***P < 0.001, ** P < 0.01, *P < 0.05, and (ns) is P > 0.05. Each gray dot (N) represents the number of animals.

**Supplemental Figure 3:**
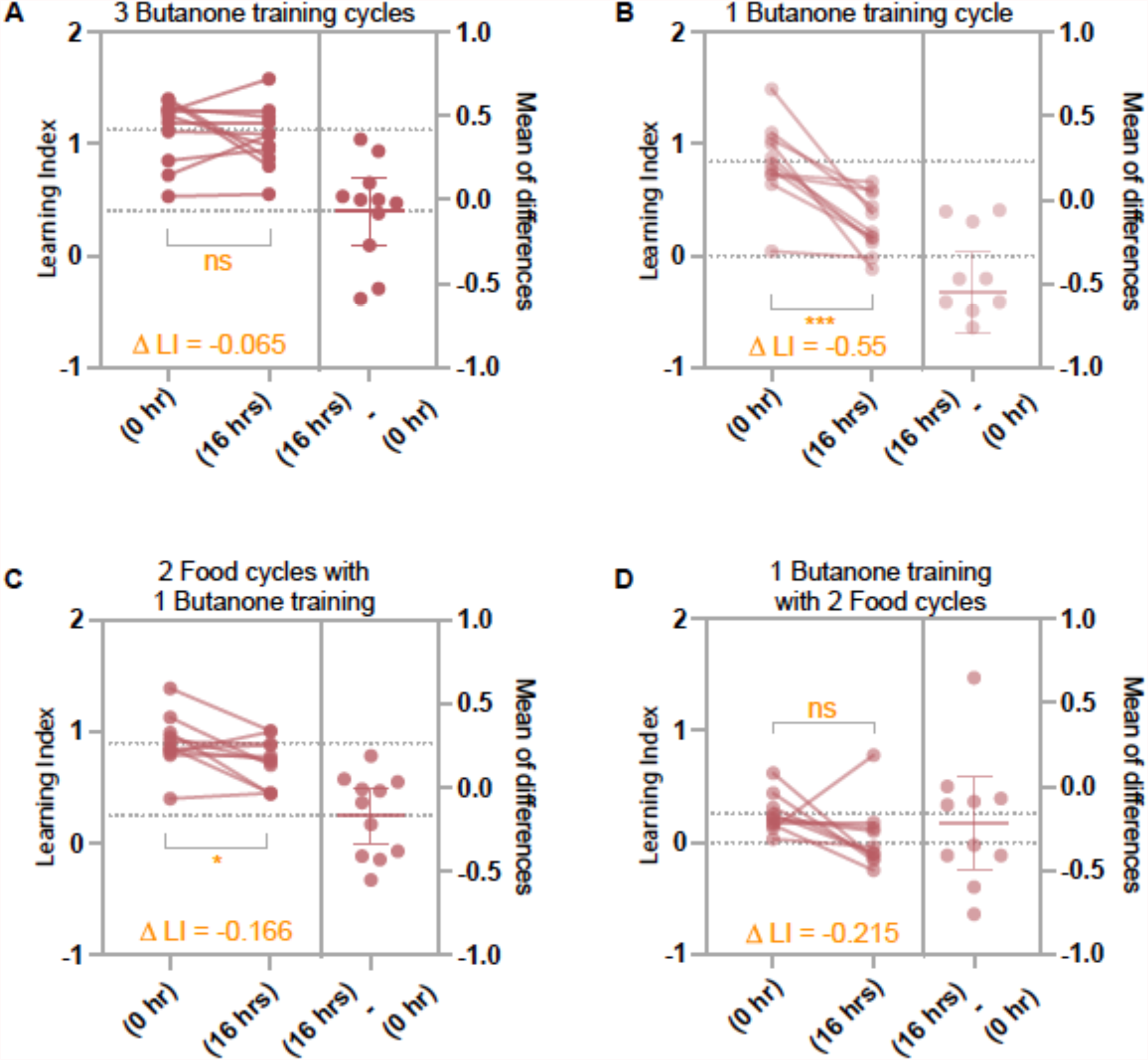
Supports Figure 4D and 4E. (A) One tailed paired t-tests of 3 butanone training cycles show that the LIs remain similar during learning (0 hour) and during memory (16 hours later) suggesting that LI decay is negligible. N= 10 trials (B) One tailed paired t-tests of 1 butanone training cycle show that the LIs fall significantly from learning (0 hour) to memory (16 hours later) suggesting that memory decays after 1 butanone training cycle. N= 9 trials (C) One tailed paired t-tests of 2 food cycles with 1 butanone training cycle show that although the LIs fall significantly from learning (0 hour) to memory (16 hours later) the amount of LI decay (-0.166) is much lower than just 1 butanone training cycle (-0.55). N = 10 trials (D) One tailed paired t-tests of 1 butanone training cycle plus 2 food cycles show that the LIs don’t fall significantly from learning (0 hour) to memory (16 hours later) and the amount of LI decay (-0.215) is negligible. N = 10 trials.

**Supplemental Figure 4:**
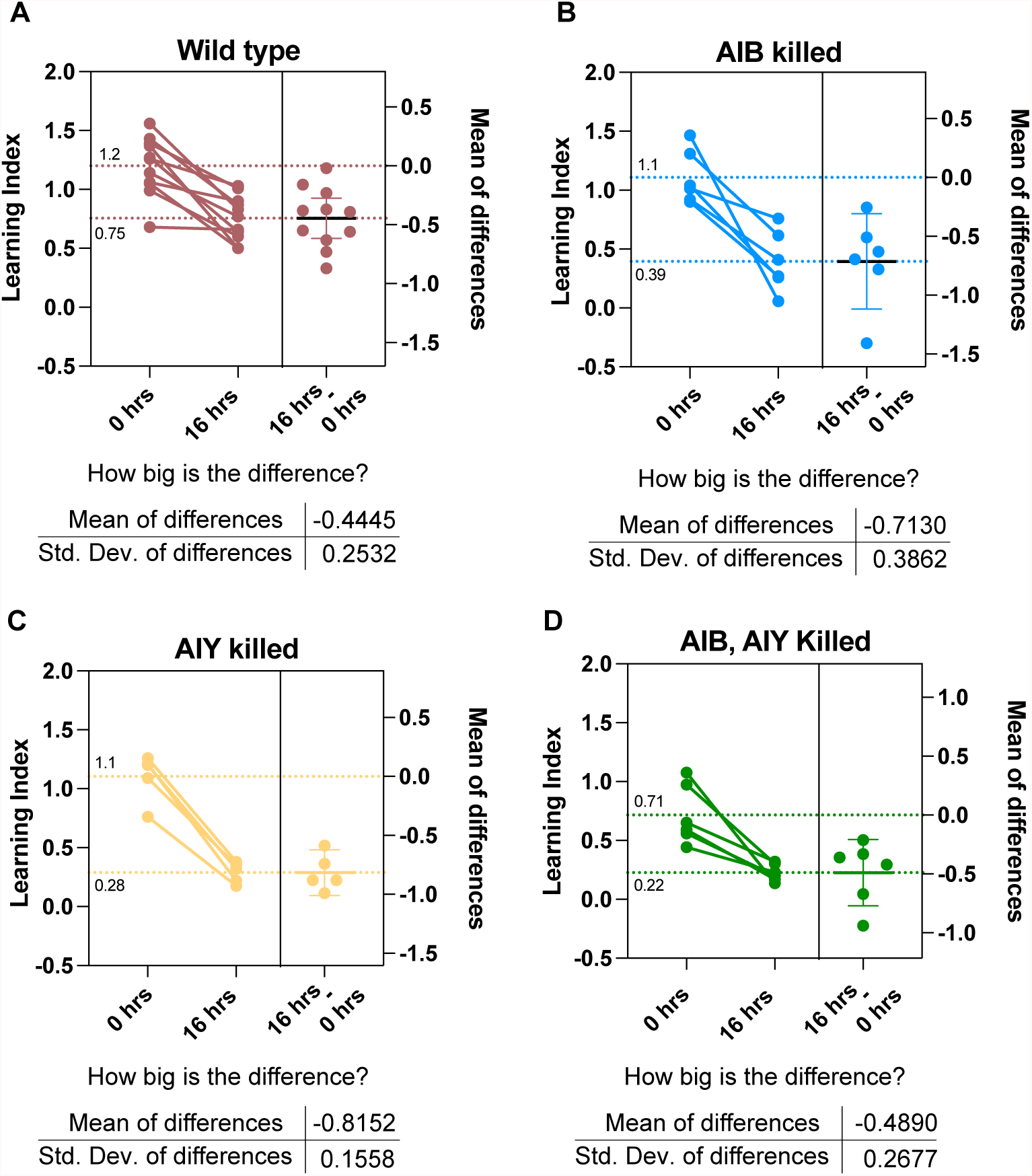
Supports Figure 5F and 5G. (A) The degree of depreciation in the LIs of animals with intact AIB and AIY neurons, and the average of depreciation observed in all the trials are shown. The LIs of wild type animals decreases from an average of 1.2 to 0.75 from 0 to 16 hours. (B) The LI depreciation in animals with AIB killed range from an average of 1.1 to 0.39, however, the range of observed LI differences are higher due to increased variability. (C) The AIY killed animals exhibit the biggest LI depreciation between 0 and 16 hours within a range of average 1.1 LI at 0 hours to 0.28 LI at 16 hours with least variability. (D) The AIB|AIY killed animals exhibit the least LI loss between 0 and 16 hours (average LI from 0.71 to 0.22). This data also show that they learned least, and therefore, AIB|AIY killed animals retained least memory.

**Figure S5.**
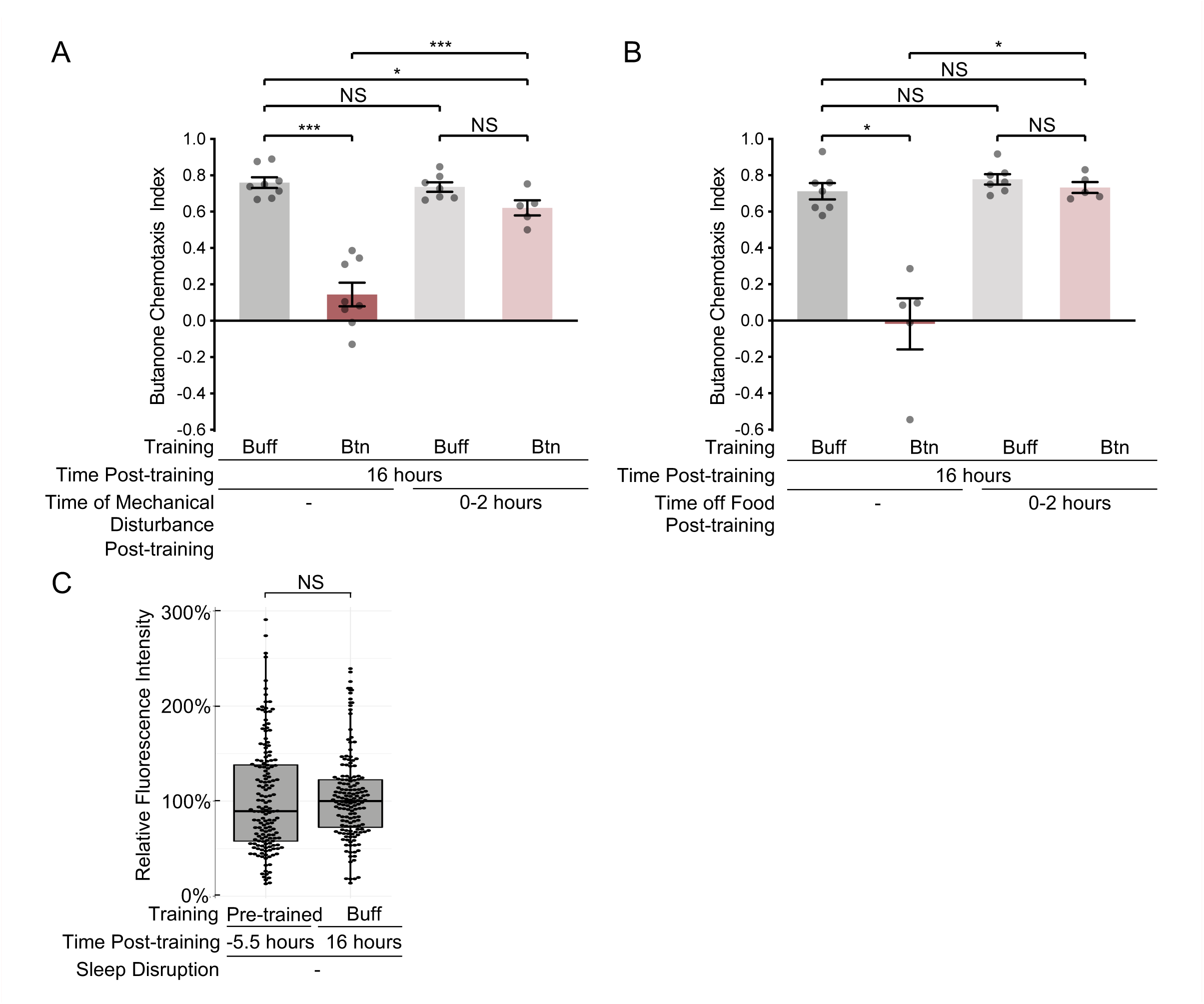
Chemotaxis of AWC-AIY NLG-1 GRASP-carrying animal cohorts assessed for NLG-1. GRASP intensity in sleep disruption experiments, and synaptic intensity of pre-trained animals. (A, B) Chemotaxis indices of AWC-AIY NLG-1 GRASP-carrying animals trained for Figure 6A (A) and Figure 6B (B). Animals were imaged from buffer-trained (Buff) batches and butanone-trained (Btn) batches whose sleep was disrupted that sensed butanone (CI>0.5), and from butanone-trained batches whose sleep was not disrupted that did not sense butanone well (CI<0.5). NS P>0.05, * P<0.05, *** P<0.001, t-test. P-values were adjusted for multiple comparisons using the Hochberg procedure. (C) AWC-AIY NLG-1 GRASP fluorescence intensity in animals immediately before training (5.5 hours before training was complete) was similar to that observed in buffer-trained animals 16 hours after training was complete. NS P>0.05. P-values were adjusted for multiple comparisons using the Hochberg procedure.

**Figure S6.**
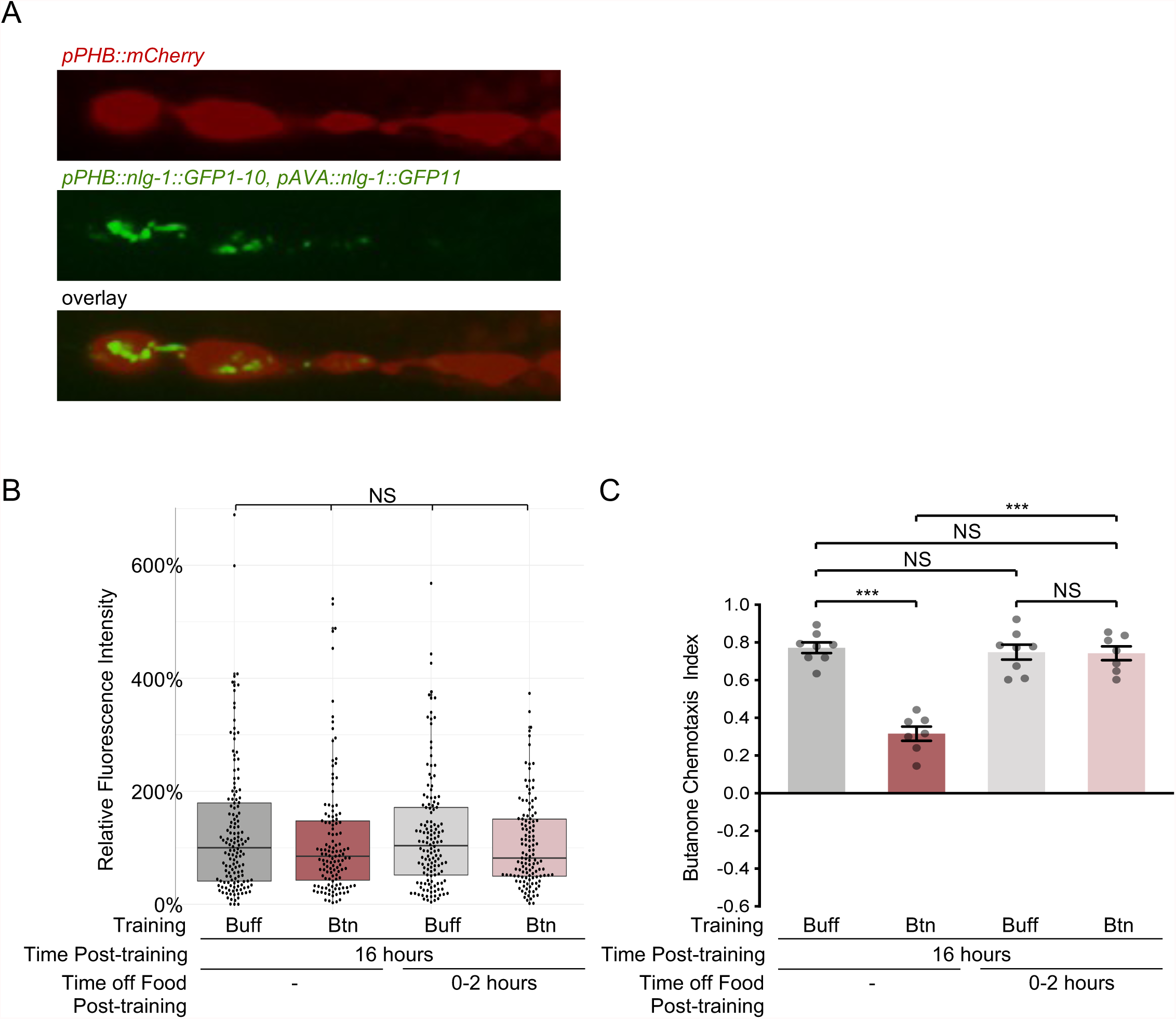
Butanone-training does not affect PHB-AVA synapses. **(A)** NLG-1 GRASP labeling synapses between the left and right PHB chemosensory neurons and the AVA interneurons. PHB neurons are labeled with cytosolic mCherry. **(B)** Quantification of PHB-AVA NLG-1 GRASP fluorescence intensity in animals trained with buffer (Buff) or butanone (Btn) whose sleep was not disrupted (left two boxes), or whose sleep was disrupted by removal from food for two hours immediately after training (right two boxes). There is no significant difference between the four training groups. NS P>0.05, Kruskal-Wallis test. **(C)** Chemotaxis indices of PHB-AVA NLG-1 GRASP-carrying animals trained for panel E. Animals were imaged from buffer-trained batches and butanone-trained batches whose sleep was disrupted that sensed butanone (CI>0.5), and from butanone-trained batches whose sleep was not disrupted that did not sense butanone well (CI<0.5). NS P>0.05, *** P<0.001, t-test. P-values were adjusted for multiple comparisons using the Hochberg procedure.

**Figure S7.**
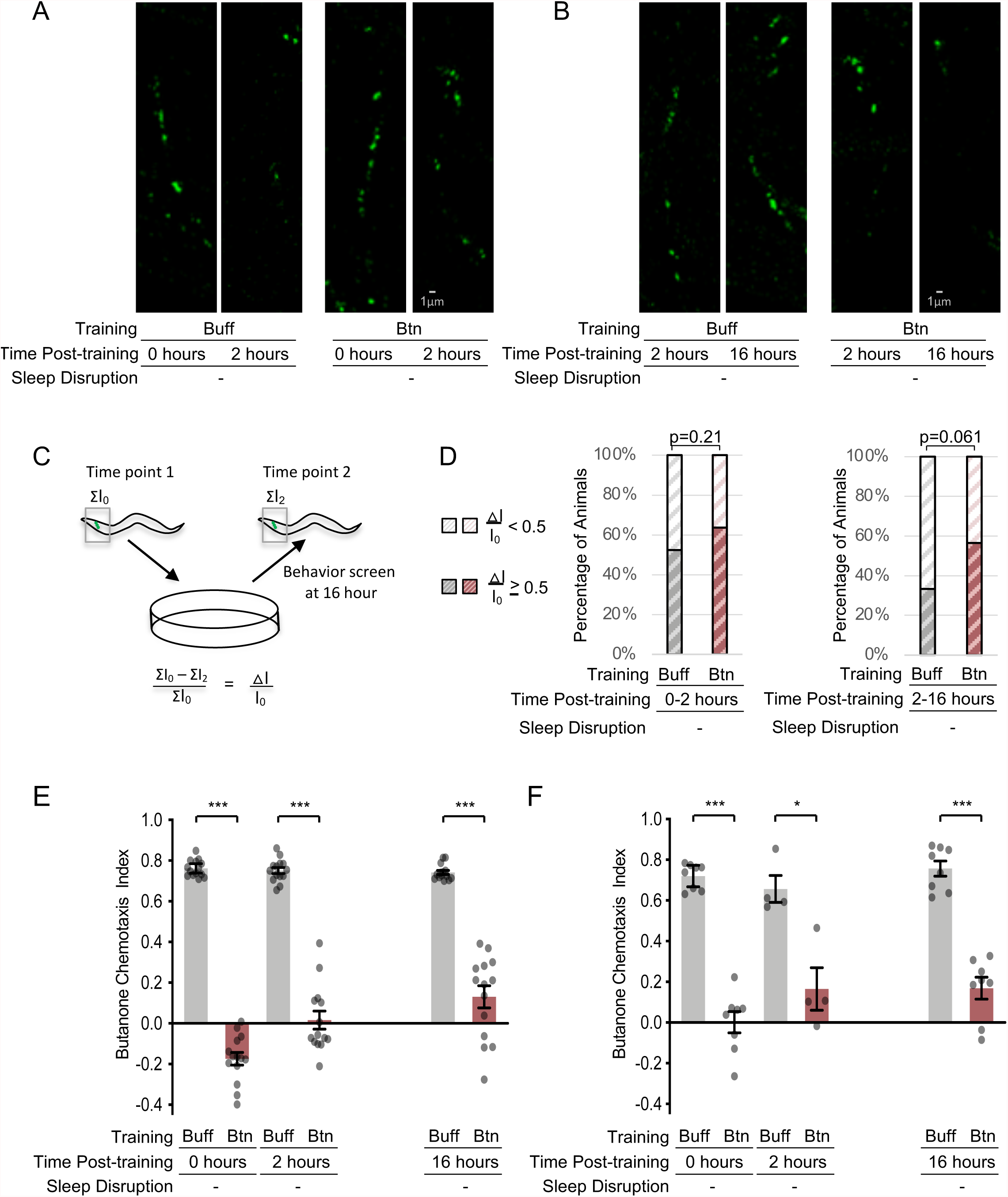
Single-worm synaptic imaging time course studies reveal synaptic reductions over time. **(A)** AWC-AIY NLG-1 GRASP fluorescence in buffer-trained (Buff) animal (left) and butanone-trained (Btn) animal (right), each imaged at 0 and 2 hours after training. **(B)** Buffer-trained (Buff) animal (left) and butanone-trained (Btn) animal (right), each imaged at 2 and 16 hours after training. **(C)** Schematic of procedure for imaging animals and quantifying the change in intensity between the first and second time points, including single-worm behavioral screens performed 16 hours after training. **(D)** Proportions of animals with large reductions (≥ 50%) in AWC-AIY NLG-1 GRASP fluorescence intensity between 0 and 2 hours after training with buffer or butanone (left) and between 2 and 16 hours (right). N>20 for each group. P-values were calculated using the z-test. **(E, F)** Batch chemotaxis indices of AWC-AIY NLG-1 GRASP-carrying animals trained for experiments in A-D (E) and for experiments in Figure 7 (F). Animals were imaged from buffer-trained (Buff) batches and butanone-trained (Btn) batches whose sleep was disrupted that sensed butanone (CI>0.5), and from butanone-trained batches whose sleep was not disrupted that did not sense butanone well (CI<0.5). NS P>0.05, *** P<0.001, * P<0.05, t-test. P-values were adjusted for multiple comparisons using the Hochberg procedure. Note that for the experiments in panel F, only one of each set of two training batches on each day could be tested for chemotaxis two hours after training, given the timing constraints due to training and imaging.

**Supplemental Video 1:**
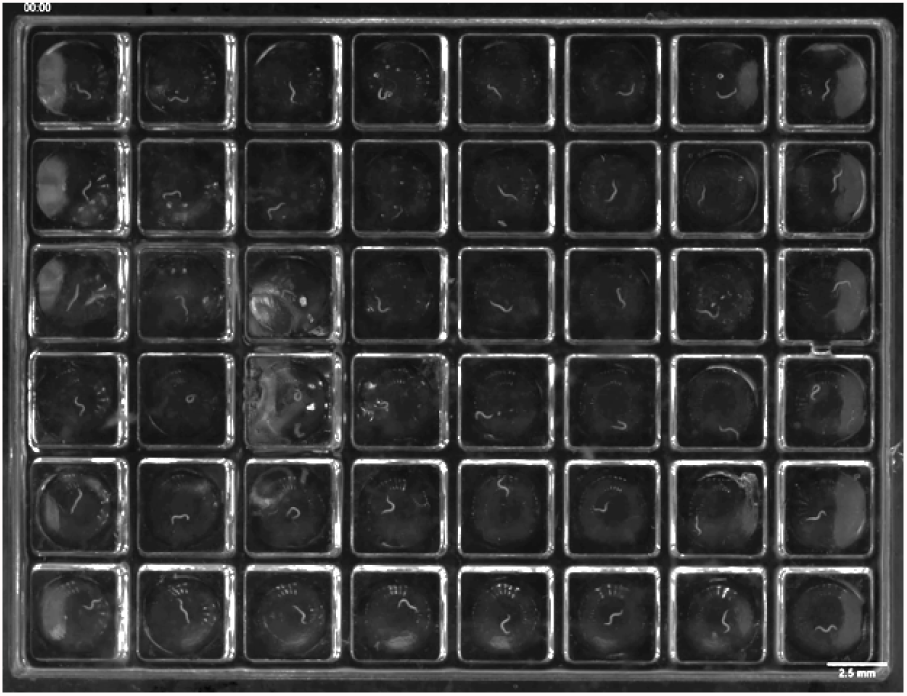
Supports Figure 1.

**Supplemental Video 2:**
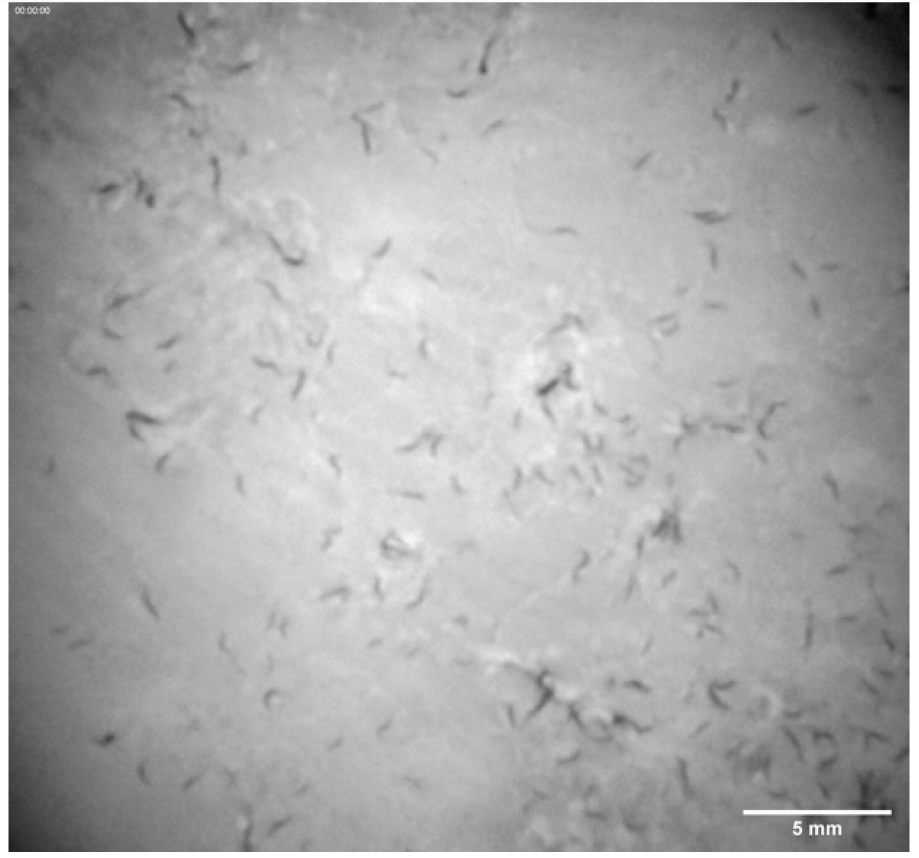
Supports Figure 2.

**Supplemental Video 3:**
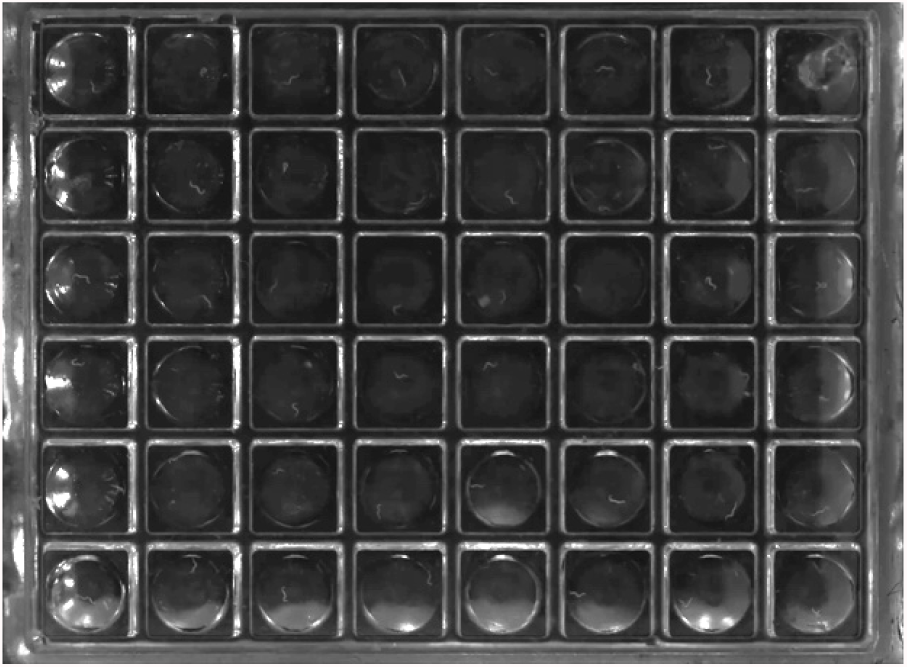
Supports Figure 2.

A single WorMotel setup to observe and quantify sleep disruption induced by removal from food. Top left, buffer-trained in food containing wells, top right butanone-trained animals on food. Bottom left, buffer-trained worms without food, bottom right butanone-trained worms without food. Video sped up 40X.

**Supplemental Table 1:** Supports Figure 2. Multiple comparisons of two-way and one-way ANOVA related to Figure 2B and 2C (mechanically disturbed worms)

| Bonferroni's multiple comparisons test | Mean Diff. | 95.00% CI of diff. | Below threshold? | Summary | Adjusted P-value |
| --- | --- | --- | --- | --- | --- |
| Post training vs. None | 0.2307 | -0.1589 to 0.6203 | No | ns | 0.8538 |
| Post training vs. 0-2hr | 0.992 | 0.6024 to 1.382 | Yes | **** | <0.0001 |
| Post training vs. 2-4hr | 0.404 | -0.04591 to 0.8539 | No | ns | 0.1091 |
| Post training vs. 4-6hr | 0.4357 | -0.01424 to 0.8856 | No | ns | 0.064 |
| None vs. 0-2hr | 0.7613 | 0.3717 to 1.151 | Yes | **** | <0.0001 |
| None vs. 2-4hr | 0.1733 | -0.2766 to 0.6232 | No | ns | >0.9999 |
| None vs. 4-6hr | 0.205 | -0.2449 to 0.6549 | No | ns | >0.9999 |
| 0-2hr vs. 2-4hr | -0.588 | -1.038 to -0.1381 | Yes | ** | 0.0039 |
| 0-2hr vs. 4-6hr | -0.5563 | -1.006 to -0.1064 | Yes | ** | 0.0071 |
| 2-4hr vs. 4-6hr | 0.03167 | -0.4713 to 0.5347 | No | ns | >0.9999 |

| Test details | Mean 1 | Mean 2 | Mean Diff. | SE of diff. | n1 | n2 |
| --- | --- | --- | --- | --- | --- | --- |
| Post training vs. None | 1.174 | 0.9433 | 0.2307 | 0.1305 | 10 | 10 |
| Post training vs. 0-2hr | 1.174 | 0.182 | 0.992 | 0.1305 | 10 | 10 |
| Post training vs. 2-4hr | 1.174 | 0.77 | 0.404 | 0.1507 | 10 | 6 |
| Post training vs. 4-6hr | 1.174 | 0.7383 | 0.4357 | 0.1507 | 10 | 6 |
| None vs. 0-2hr | 0.9433 | 0.182 | 0.7613 | 0.1305 | 10 | 10 |
| None vs. 2-4hr | 0.9433 | 0.77 | 0.1733 | 0.1507 | 10 | 6 |
| None vs. 4-6hr | 0.9433 | 0.7383 | 0.205 | 0.1507 | 10 | 6 |
| 0-2hr vs. 2-4hr | 0.182 | 0.77 | -0.588 | 0.1507 | 10 | 6 |
| 0-2hr vs. 4-6hr | 0.182 | 0.7383 | -0.5563 | 0.1507 | 10 | 6 |
| 2-4hr vs. 4-6hr | 0.77 | 0.7383 | 0.03167 | 0.1685 | 6 | 6 |

**Supplemental Table 2:**
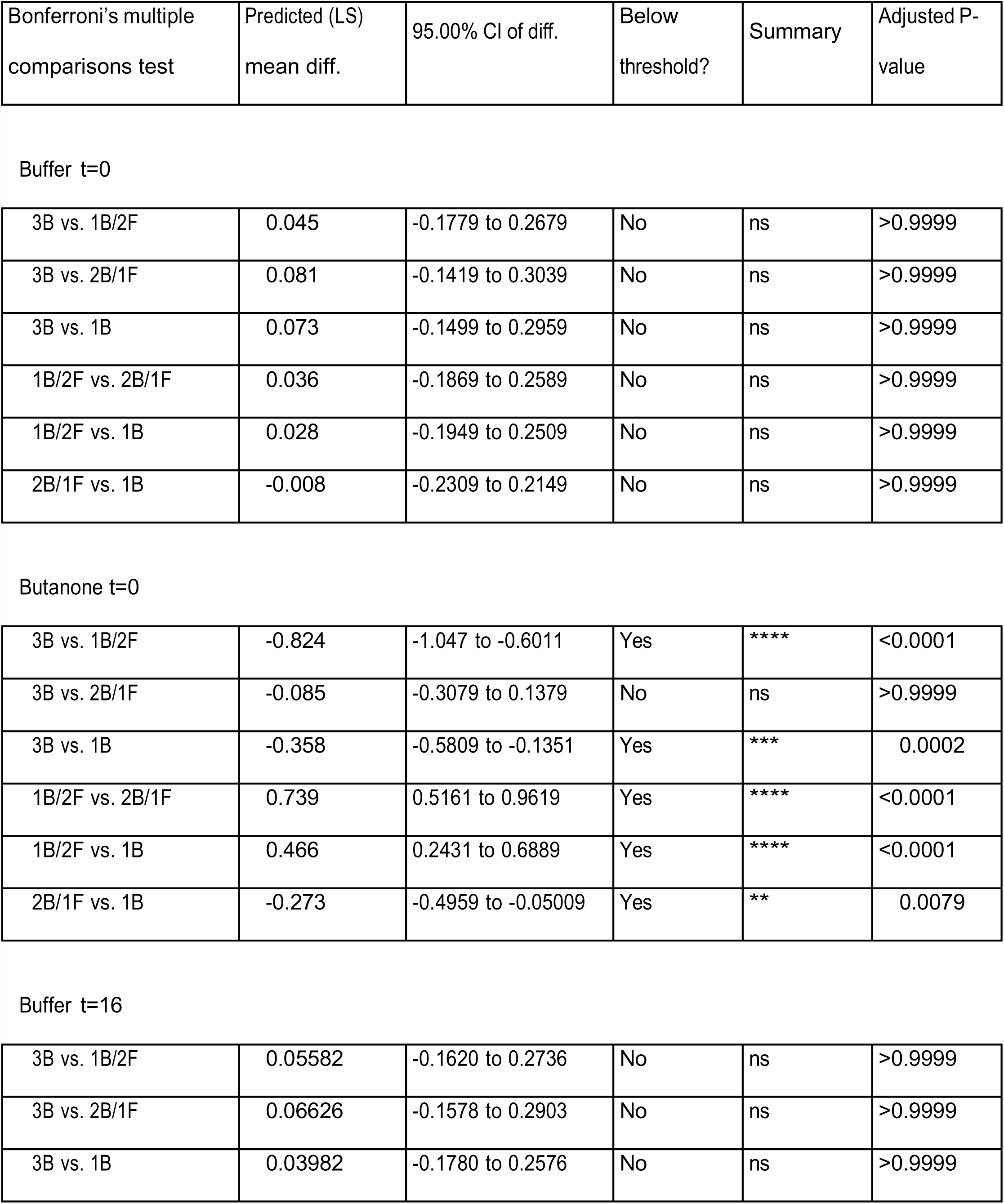
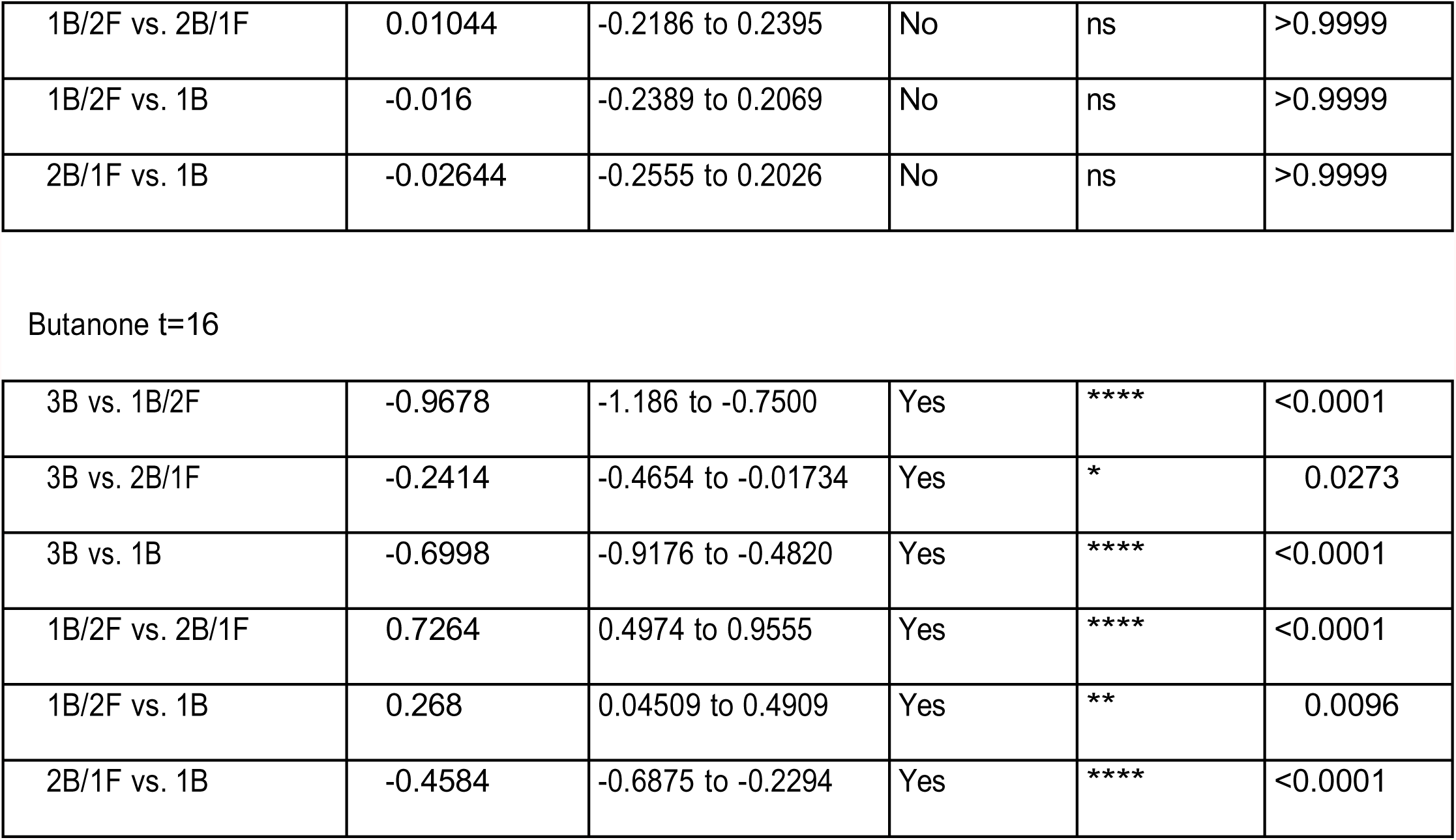
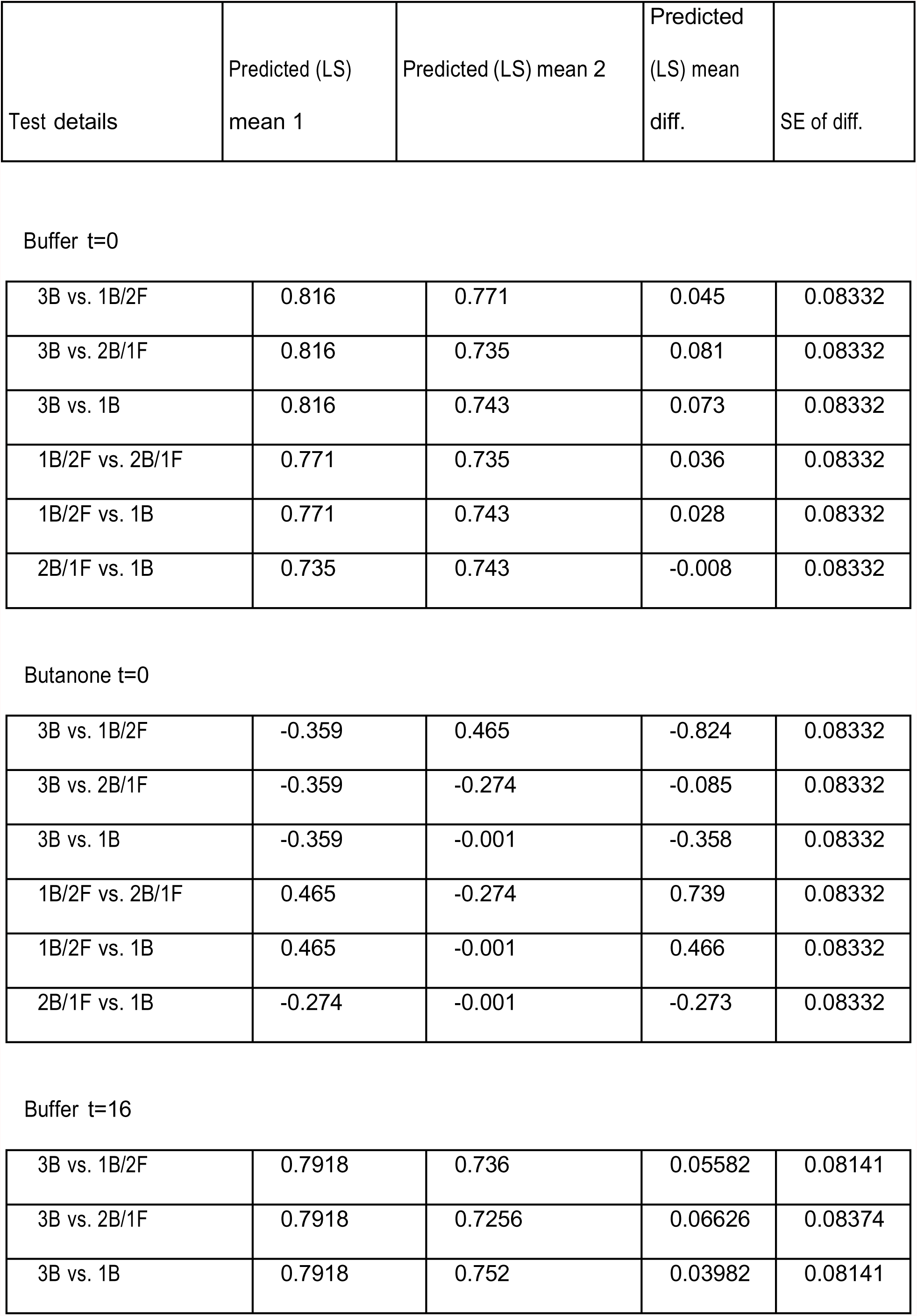
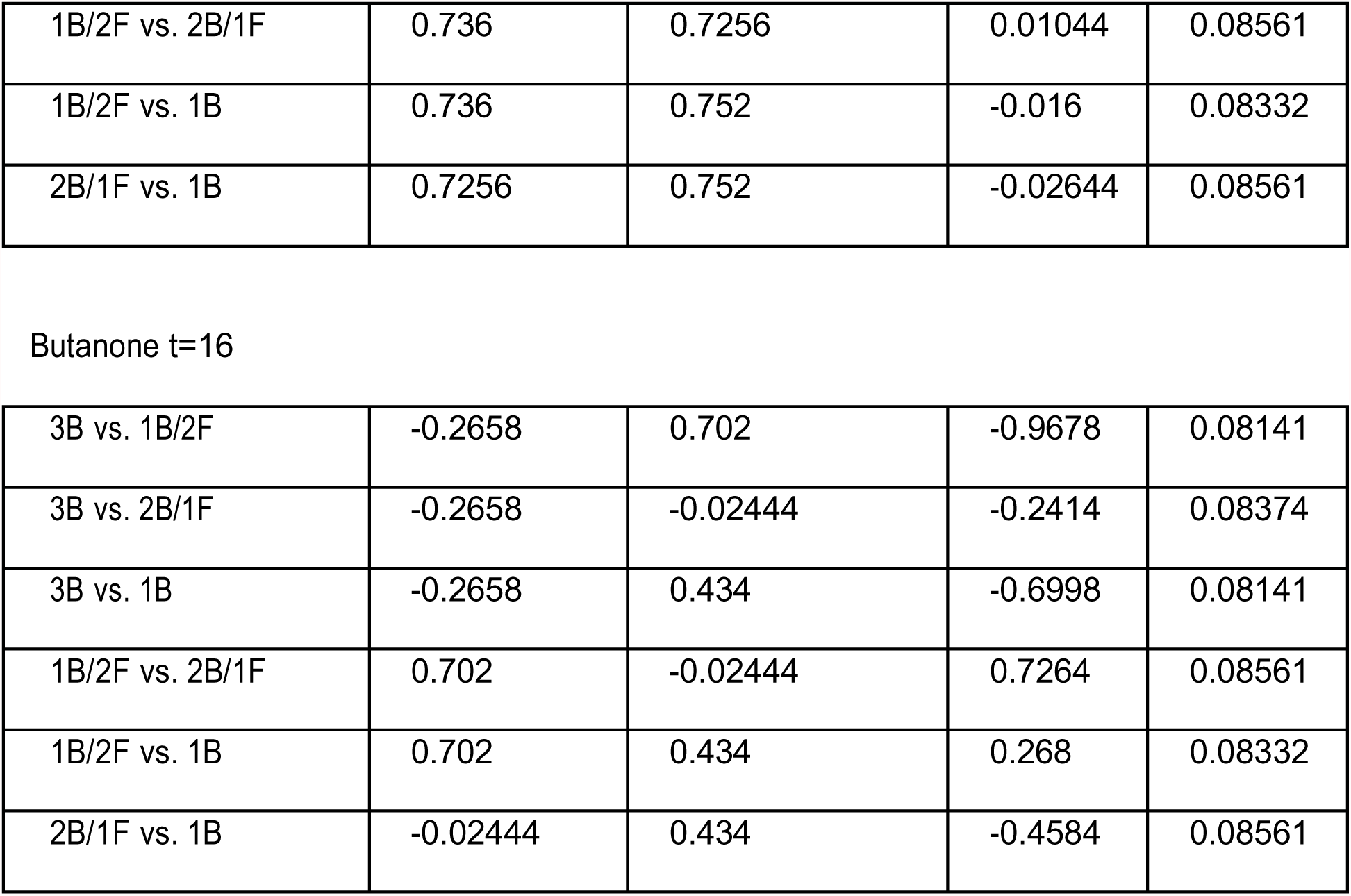
Supports Figure 3.

| Bonferroni's multiple comparisons test | Predicted (LS) mean diff. | 95.00% CI of diff. | Below threshold? | Summary | Adjusted P-value |
| --- | --- | --- | --- | --- | --- |
| Buffer t=0 |  |  |  |  |  |
| 3B vs. 1B/2F | 0.045 | -0.1779 to 0.2679 | No | ns | >0.9999 |
| 3B vs. 2B/1F | 0.081 | -0.1419 to 0.3039 | No | ns | >0.9999 |
| 3B vs. 1B | 0.073 | -0.1499 to 0.2959 | No | ns | >0.9999 |
| 1B/2F vs. 2B/1F | 0.036 | -0.1869 to 0.2589 | No | ns | >0.9999 |
| 1B/2F vs. 1B | 0.028 | -0.1949 to 0.2509 | No | ns | >0.9999 |
| 2B/1F vs. 1B | -0.008 | -0.2309 to 0.2149 | No | ns | >0.9999 |
| Butanone t=0 |  |  |  |  |  |
| 3B vs. 1B/2F | -0.824 | -1.047 to -0.6011 | Yes | **** | <0.0001 |
| 3B vs. 2B/1F | -0.085 | -0.3079 to 0.1379 | No | ns | >0.9999 |
| 3B vs. 1B | -0.358 | -0.5809 to -0.1351 | Yes | *** | 0.0002 |
| 1B/2F vs. 2B/1F | 0.739 | 0.5161 to 0.9619 | Yes | **** | <0.0001 |
| 1B/2F vs. 1B | 0.466 | 0.2431 to 0.6889 | Yes | **** | <0.0001 |
| 2B/1F vs. 1B | -0.273 | -0.4959 to -0.05009 | Yes | ** | 0.0079 |
| Buffer t=16 |  |  |  |  |  |
| 3B vs. 1B/2F | 0.05582 | -0.1620 to 0.2736 | No | ns | >0.9999 |
| 3B vs. 2B/1F | 0.06626 | -0.1578 to 0.2903 | No | ns | >0.9999 |
| 3B vs. 1B | 0.03982 | -0.1780 to 0.2576 | No | ns | >0.9999 |
| 1B/2F vs. 2B/1F | 0.01044 | -0.2186 to 0.2395 | No | ns | >0.9999 |
| 1B/2F vs. 1B | -0.016 | -0.2389 to 0.2069 | No | ns | >0.9999 |
| 2B/1F vs. 1B | -0.02644 | -0.2555 to 0.2026 | No | ns | >0.9999 |
| Butanone t=16 |  |  |  |  |  |
| 3B vs. 1B/2F | -0.9678 | -1.186 to -0.7500 | Yes | **** | <0.0001 |
| 3B vs. 2B/1F | -0.2414 | -0.4654 to -0.01734 | Yes | * | 0.0273 |
| 3B vs. 1B | -0.6998 | -0.9176 to -0.4820 | Yes | **** | <0.0001 |
| 1B/2F vs. 2B/1F | 0.7264 | 0.4974 to 0.9555 | Yes | **** | <0.0001 |
| 1B/2F vs. 1B | 0.268 | 0.04509 to 0.4909 | Yes | ** | 0.0096 |
| 2B/1F vs. 1B | -0.4584 | -0.6875 to -0.2294 | Yes | **** | <0.0001 |

| Test details | Predicted (LS)<br>mean 1 | Predicted (LS) mean 2 | Predicted<br>(LS) mean<br>diff. | SE of diff. |
| --- | --- | --- | --- | --- |

Buffer t=0
|  |  |  |  |  |
| --- | --- | --- | --- | --- |
| 3B vs. 1B/2F | 0.816 | 0.771 | 0.045 | 0.08332 |
| 3B vs. 2B/1F | 0.816 | 0.735 | 0.081 | 0.08332 |
| 3B vs. 1B | 0.816 | 0.743 | 0.073 | 0.08332 |
| 1B/2F vs. 2B/1F | 0.771 | 0.735 | 0.036 | 0.08332 |
| 1B/2F vs. 1B | 0.771 | 0.743 | 0.028 | 0.08332 |
| 2B/1F vs. 1B | 0.735 | 0.743 | -0.008 | 0.08332 |

Butanone t=0
|  |  |  |  |  |
| --- | --- | --- | --- | --- |
| 3B vs. 1B/2F | -0.359 | 0.465 | -0.824 | 0.08332 |
| 3B vs. 2B/1F | -0.359 | -0.274 | -0.085 | 0.08332 |
| 3B vs. 1B | -0.359 | -0.001 | -0.358 | 0.08332 |
| 1B/2F vs. 2B/1F | 0.465 | -0.274 | 0.739 | 0.08332 |
| 1B/2F vs. 1B | 0.465 | -0.001 | 0.466 | 0.08332 |
| 2B/1F vs. 1B | -0.274 | -0.001 | -0.273 | 0.08332 |

Buffer t=16
|  |  |  |  |  |
| --- | --- | --- | --- | --- |
| 3B vs. 1B/2F | 0.7918 | 0.736 | 0.05582 | 0.08141 |
| 3B vs. 2B/1F | 0.7918 | 0.7256 | 0.06626 | 0.08374 |
| 3B vs. 1B | 0.7918 | 0.752 | 0.03982 | 0.08141 |
| 1B/2F vs. 2B/1F | 0.736 | 0.7256 | 0.01044 | 0.08561 |
| 1B/2F vs. 1B | 0.736 | 0.752 | -0.016 | 0.08332 |
| 2B/1F vs. 1B | 0.7256 | 0.752 | -0.02644 | 0.08561 |

Butanone t=16
|  |  |  |  |  |
| --- | --- | --- | --- | --- |
| 3B vs. 1B/2F | -0.2658 | 0.702 | -0.9678 | 0.08141 |
| 3B vs. 2B/1F | -0.2658 | -0.02444 | -0.2414 | 0.08374 |
| 3B vs. 1B | -0.2658 | 0.434 | -0.6998 | 0.08141 |
| 1B/2F vs. 2B/1F | 0.702 | -0.02444 | 0.7264 | 0.08561 |
| 1B/2F vs. 1B | 0.702 | 0.434 | 0.268 | 0.08332 |
| 2B/1F vs. 1B | -0.02444 | 0.434 | -0.4584 | 0.08561 |

**Supplemental Table 3:**
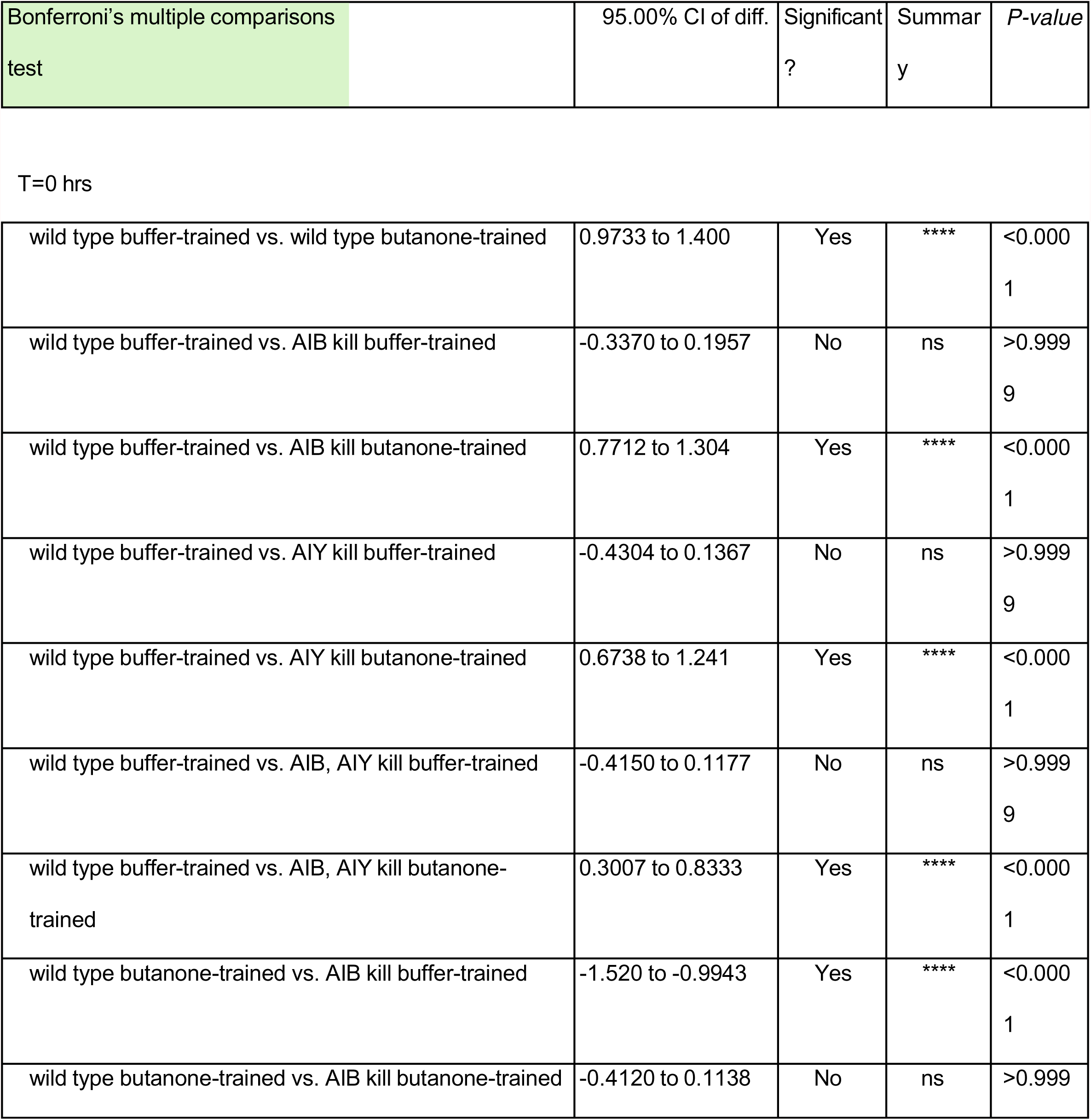
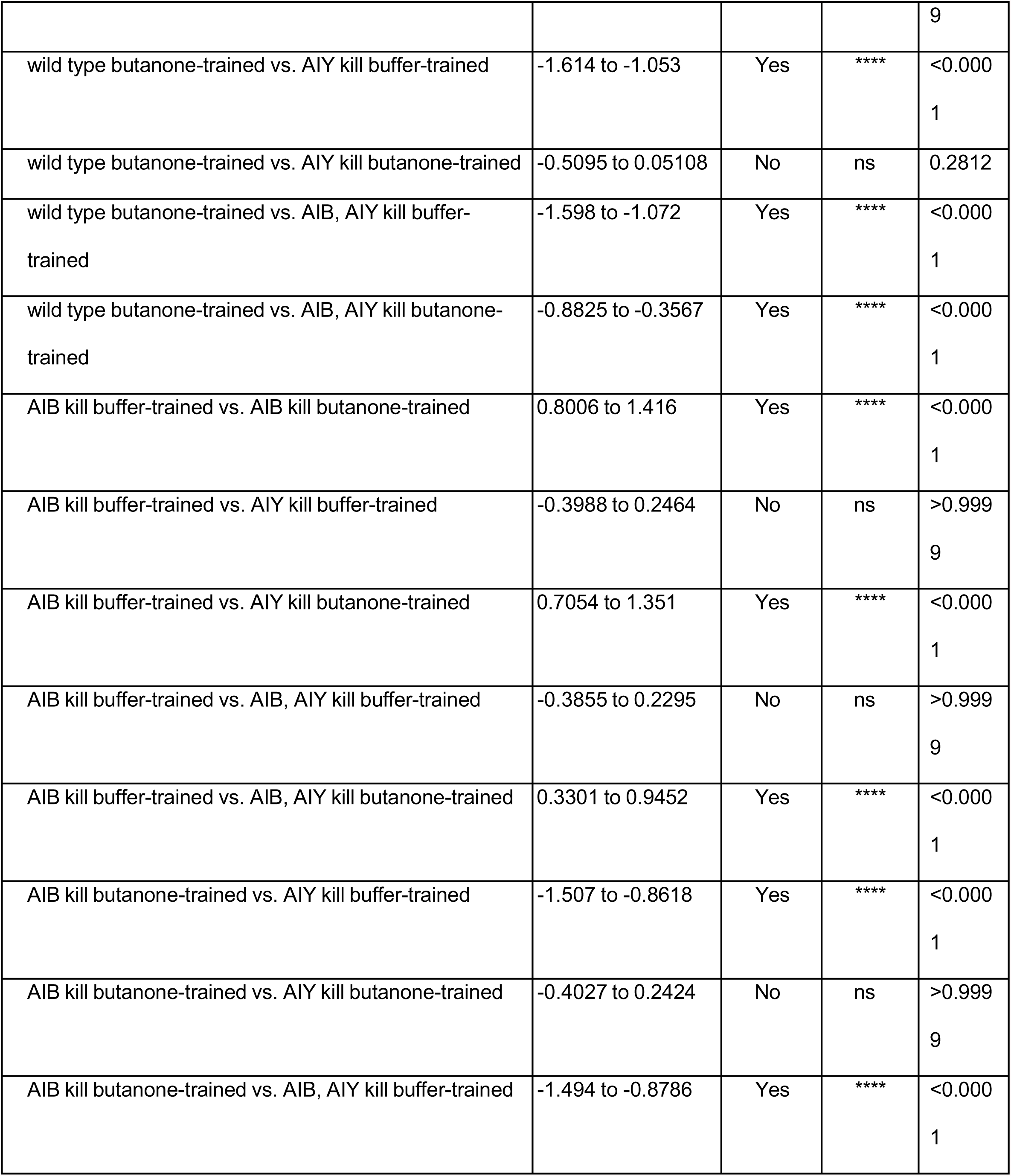
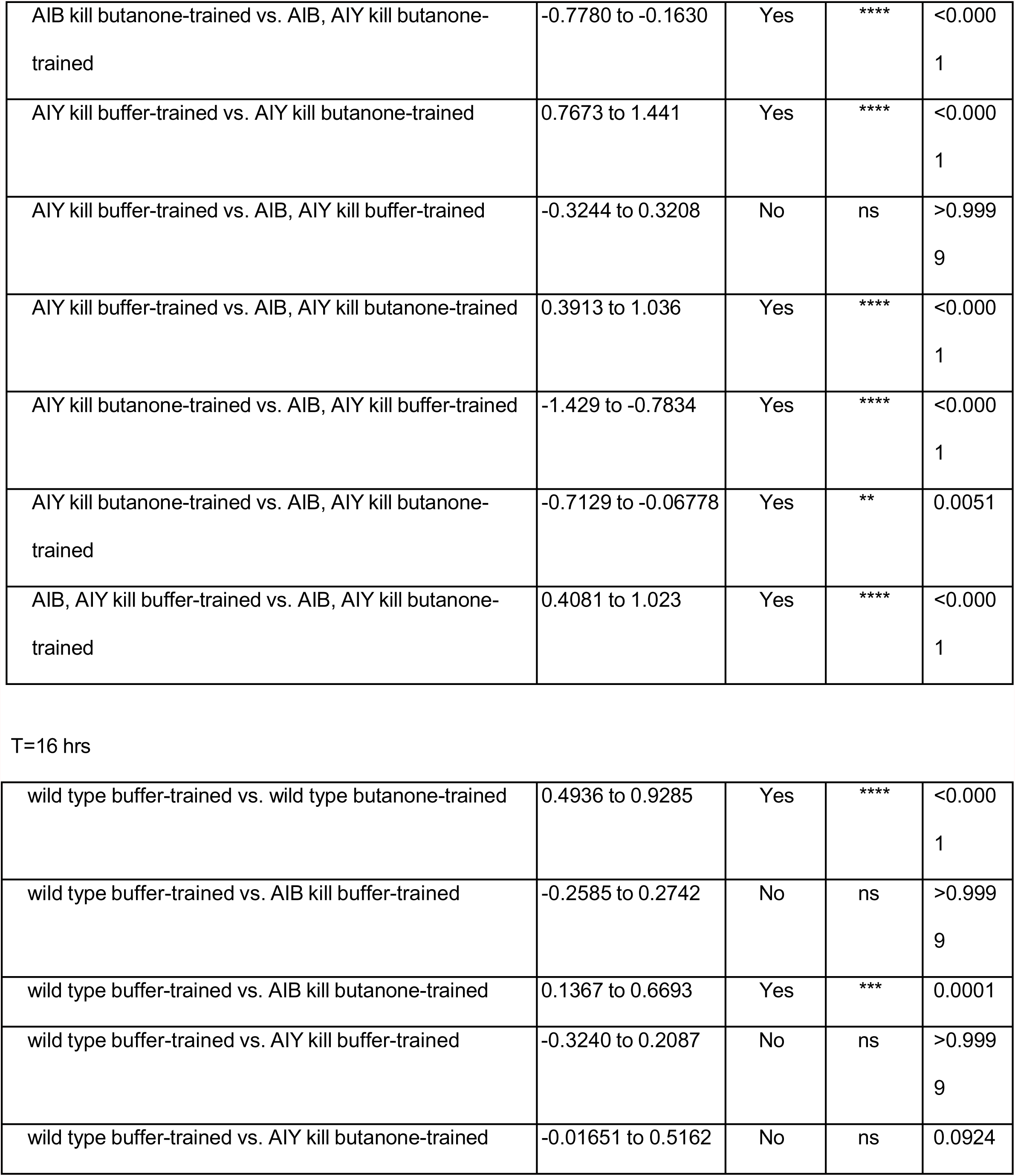
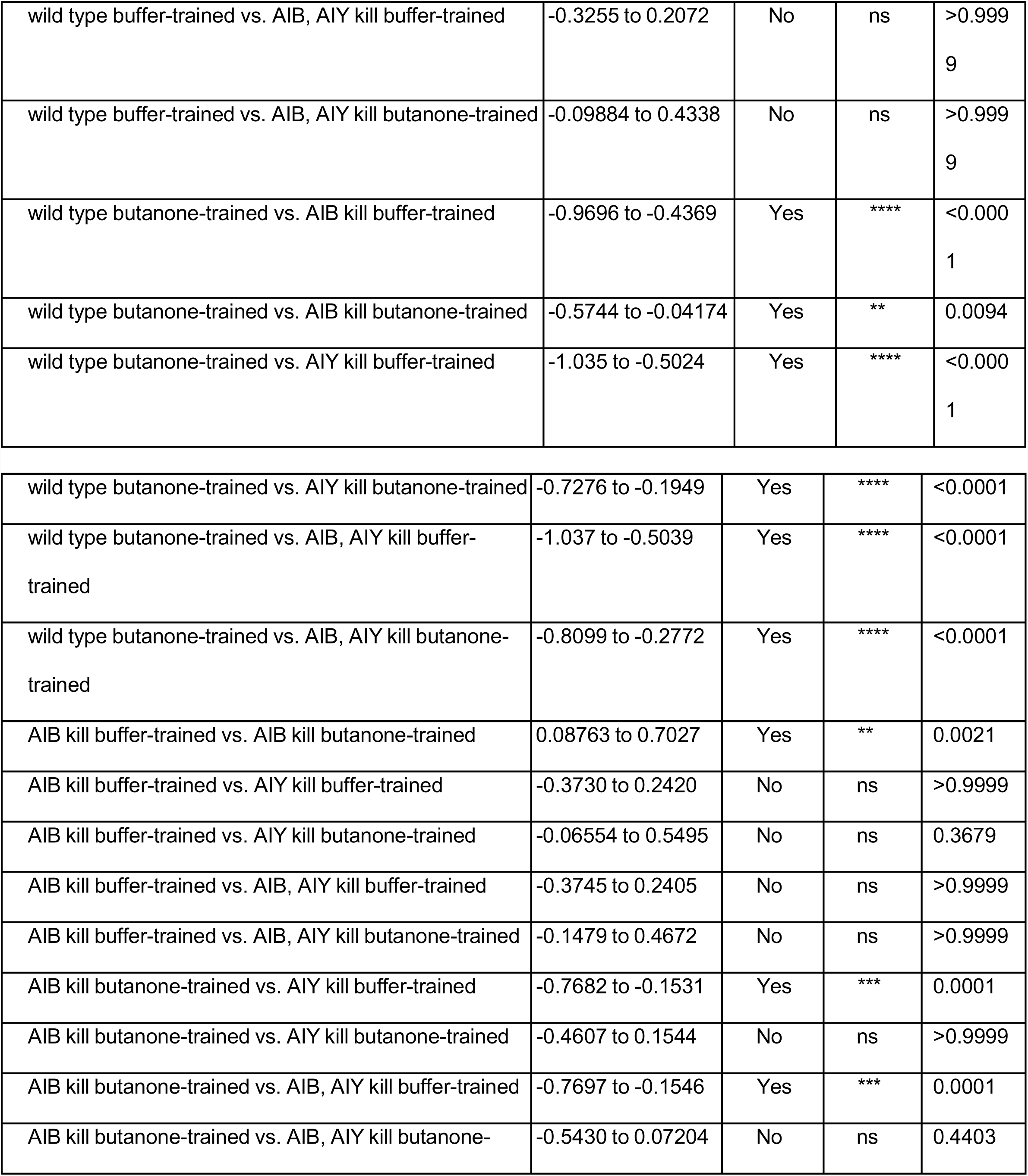
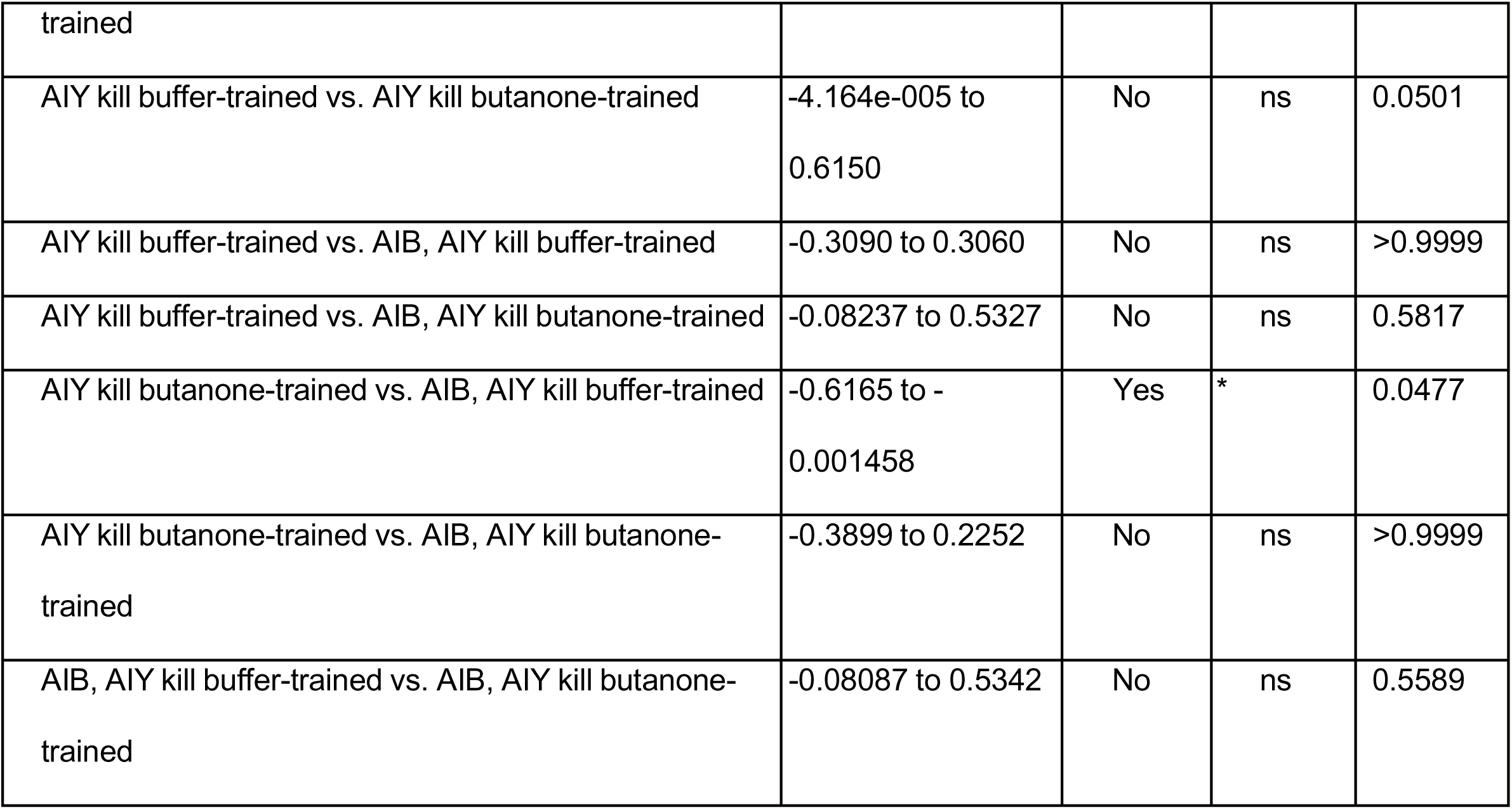
Supports Figure 4.

**Supplemental Table 4.** Supports Figure 6, 7, S6 and S7.

| **Figure panel** | **Comparison** | **Statistical test** | **P-value** |
| --- | --- | --- | --- |
| 6C | All | Kruskal-Wallis | 3.9E-3 |
| 6C | Buff no disturbance to Btn no disturbance | U-test, followed by Hochberg | 1.6E-2 |
| 6C | Buff no disturbance to Buff with disturbance | U-test, followed by Hochberg | 0.83 |
| 6C | Buff no disturbance to Btn with disturbance | U-test, followed by Hochberg | 0.64 |
| 6C | Btn no disturbance to Btn with disturbance | U-test, followed by Hochberg | 7.3E-3 |
| 6C | Buff with disturbance to Btn with disturbance | U-test, followed by Hochberg | 0.64 |
| 6D | All | Kruskal-Wallis | 5.5E-10 |
| 6D | Buff no disturbance to Btn no disturbance | U-test, followed by Hochberg | 5.8E-6 |
| 6D | Buff no disturbance to Buff with disturbance | U-test, followed by Hochberg | 2.1E-1 |
| 6D | Buff no disturbance to Btn with disturbance | U-test, followed by Hochberg | 4.0E-1 |
| 6D | Btn no disturbance to Btn with disturbance | U-test, followed by Hochberg | 1.0E-6 |
| 6D | Buff with disturbance to Btn with disturbance | U-test, followed by Hochberg | 3.4E-1 |
| 7B | All | Kruskal-Wallis | 2.0E-15 |
| 7B | 0 Buff to 0 Btn | U-test, followed by Hochberg | 1.8E-1 |
| 7B | 0 Buff to 2 Buff | U-test, followed by Hochberg | 1.2E-2 |
| 7B | 0 Buff to 16 Buff | U-test, followed by Hochberg | 5.6E-6 |
| 7B | 0 Btn to 2 Btn | U-test, followed by Hochberg | 4.6E-3 |
| 7B | 0 Btn to 16 Btn | U-test, followed by Hochberg | 8.9E-9 |
| 7B | 2 Buff to 2 Btn | U-test, followed by Hochberg | 1.8E-1 |
| 7B | 2 Buff to 16 Buff | U-test, followed by Hochberg | 3.6E-1 |
| 7B | 2 Btn to 16 Btn | U-test, followed by Hochberg | 1.8E-1 |
| 7B | 16 Buff to 16 Btn | U-test, followed by Hochberg | 4.1E-4 |
| S5A | All | One-way ANOVA | 2.6E-10 |
| S5A | Buff no disturbance to Btn no disturbance | t-test, followed by Hochberg | 1.1E-4 |
| S5A | Buff no disturbance to Buff disturbance | t-test, followed by Hochberg | 0.81 |
| S5A | Buff no disturbance to Btn disturbance | t-test, followed by Hochberg | 7.5E-3 |
| S5A | Btn no disturbance to Btn no disturbance | t-test, followed by Hochberg | 4.1E-4 |
| S5A | Buff disturbance to Btn disturbance | t-test, followed by Hochberg | 0.63 |
| S5B | All | One-way ANOVA | 1.9E-7 |
| S5B | Buff no disturbance to Btn no disturbance | t-test, followed by Hochberg | 2.0E-2 |
| S5B | Buff no disturbance to Buff disturbance | t-test, followed by Hochberg | 0.61 |
| S5B | Buff no disturbance to Btn disturbance | t-test, followed by Hochberg | 0.71 |
| S5B | Buff disturbance to Btn disturbance | t-test, followed by Hochberg | 0.61 |
| S5B | Btn no disturbance to Btn disturbance | t-test, followed by Hochberg | 2.0E-2 |
| S5C | Naïve pre-trained worms to 16 Buff | U-test, followed by Hochberg | 0.66 |
| S6B | All | Kruskal-Wallace | 0.58 |
| S6C | All | One-way ANOVA | 2.2E-9 |
| S6C | Buff no disturbance to Btn no disturbance | t-test, followed by Hochberg | 3.7E-6 |
| S6C | Buff no disturbance to Buff disturbance | t-test, followed by Hochberg | 9.1E-1 |
| S6C | Buff no disturbance to Btn disturbance | t-test, followed by Hochberg | 9.1E-1 |
| S6C | Btn no disturbance to Btn no disturbance | t-test, followed by Hochberg | 9.1E-1 |
| S6C | Btn no disturbance to Btn disturbance | t-test, followed by Hochberg | 1.4E-5 |
| S7D | 0-2 Buff to Btn | z-test | 0.21 |
| S7D | 2-16 Buff to Btn | z-test | 0.061 |
| S7E | All | One-way ANOVA | 2.7E-10 |
| S7E | 0 Buff to 0 Btn | t-test, followed by Hochberg | 2.6E-18 |
| S7E | 2 Buff to 2 Btn | t-test, followed by Hochberg | 6.9E-16 |
| S7E | 16 Buff to 16 Btn | t-test, followed by Hochberg | 5.2E-12 |
| S7F | All | One-way ANOVA | 4.4E-10 |
| S7F | 0 Buff to 0 Btn | t-test, followed by Hochberg | 3.6E-6 |
| S7F | 2 Buff to 2 Btn | t-test, followed by Hochberg | 6.6E-2 |
| S7F | 16 Buff to 16 Btn | t-test, followed by Hochberg | 7.8E-6 |

## STAR METHODS

### KEY RESOURCES TABLE

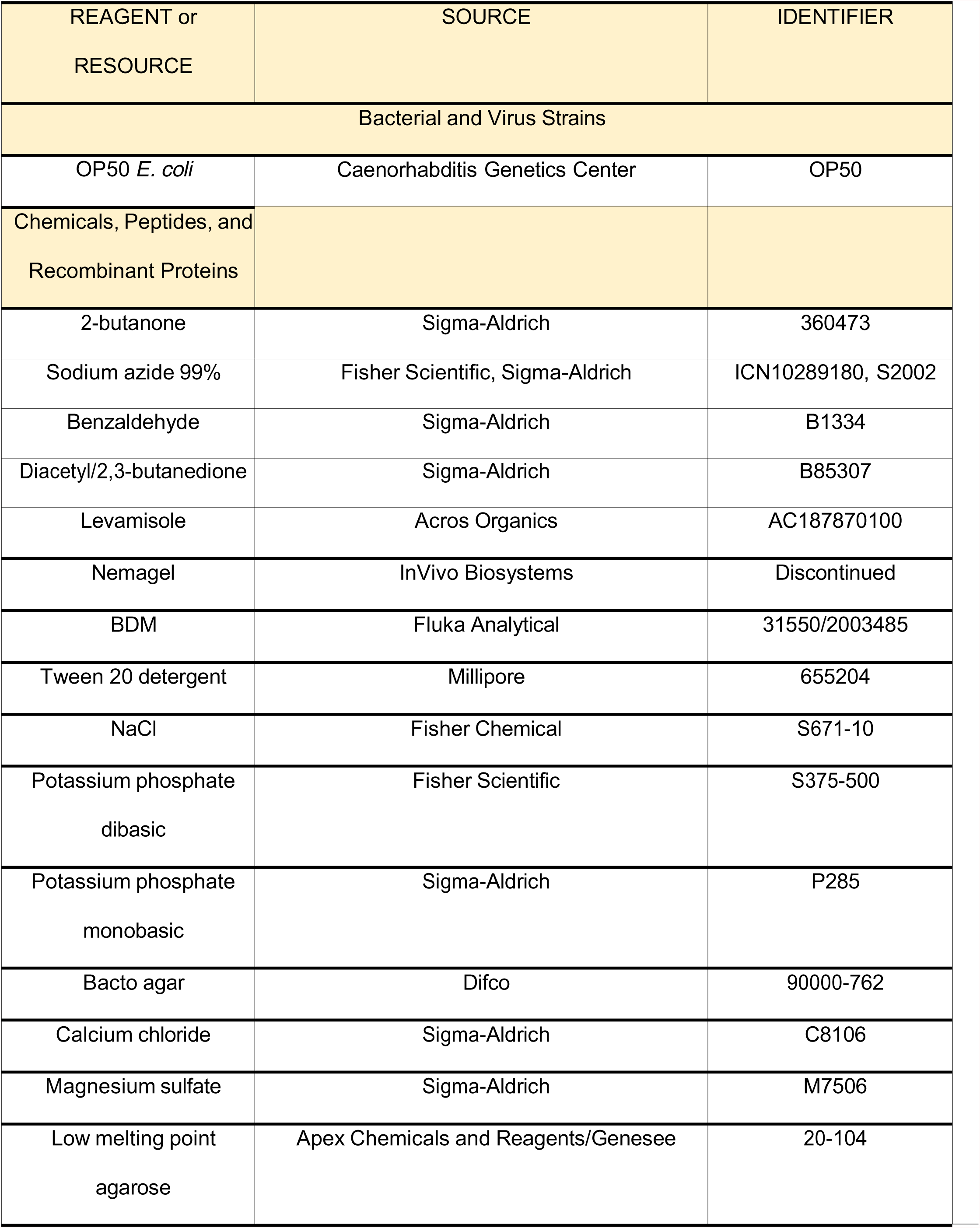
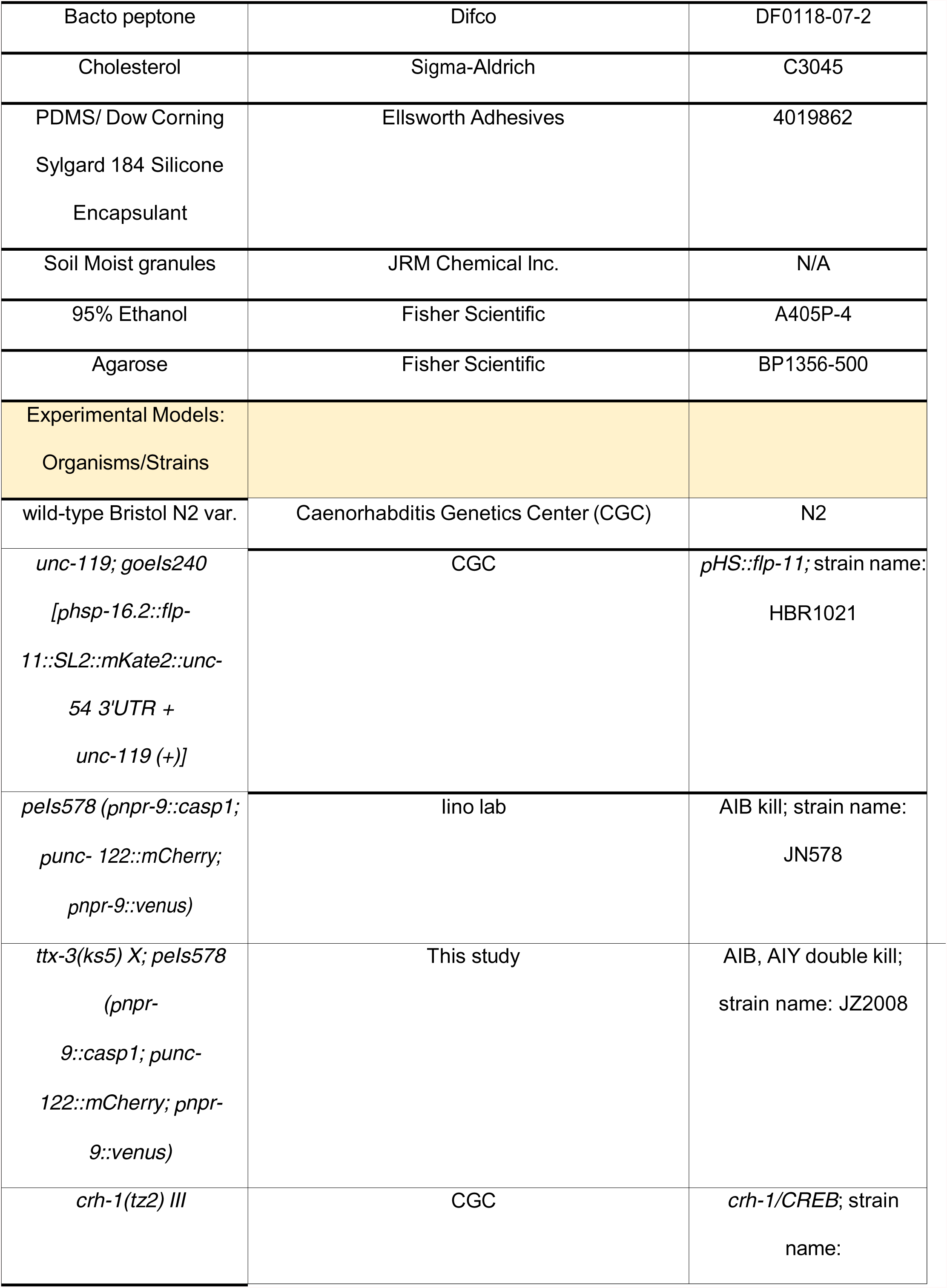
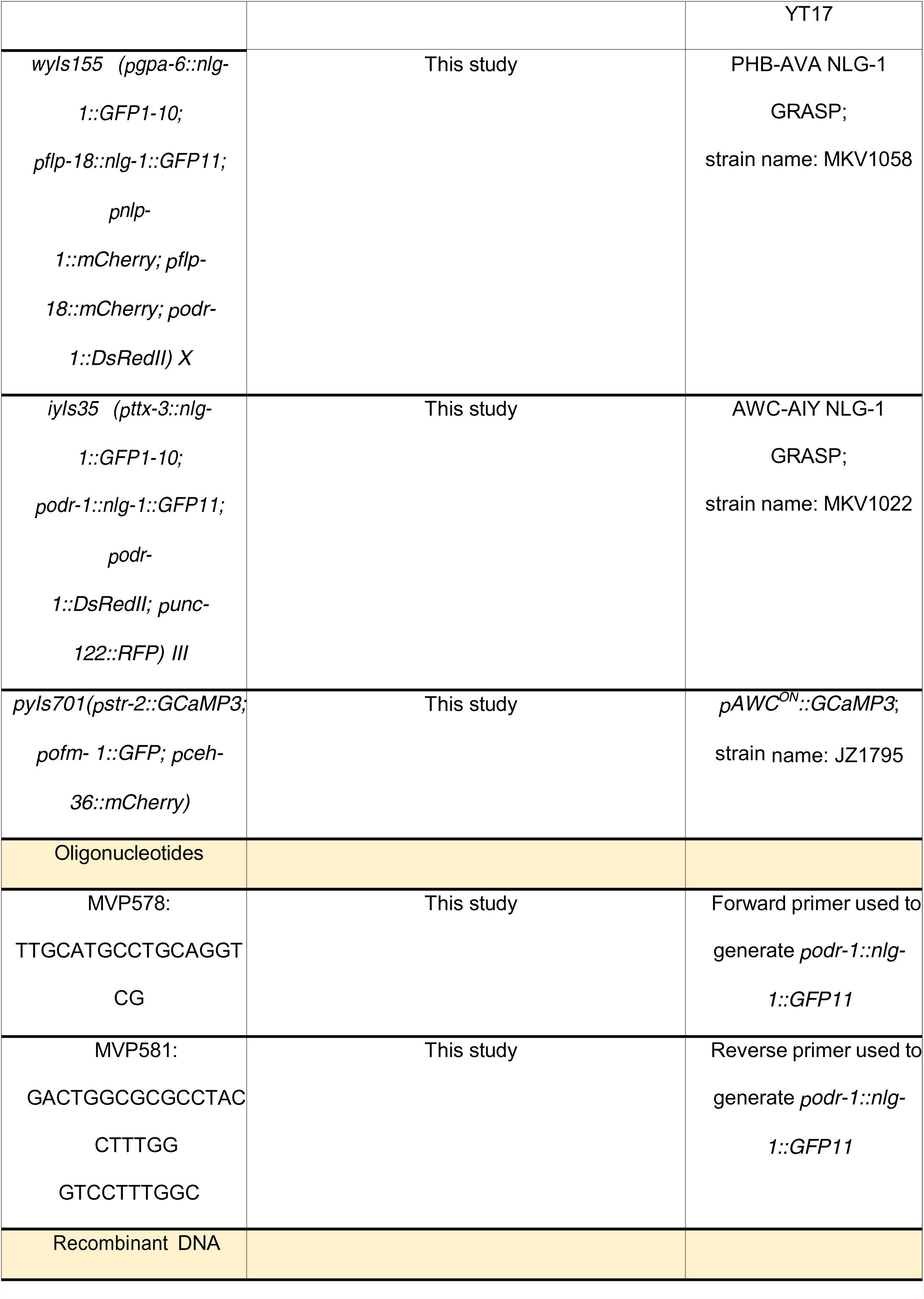
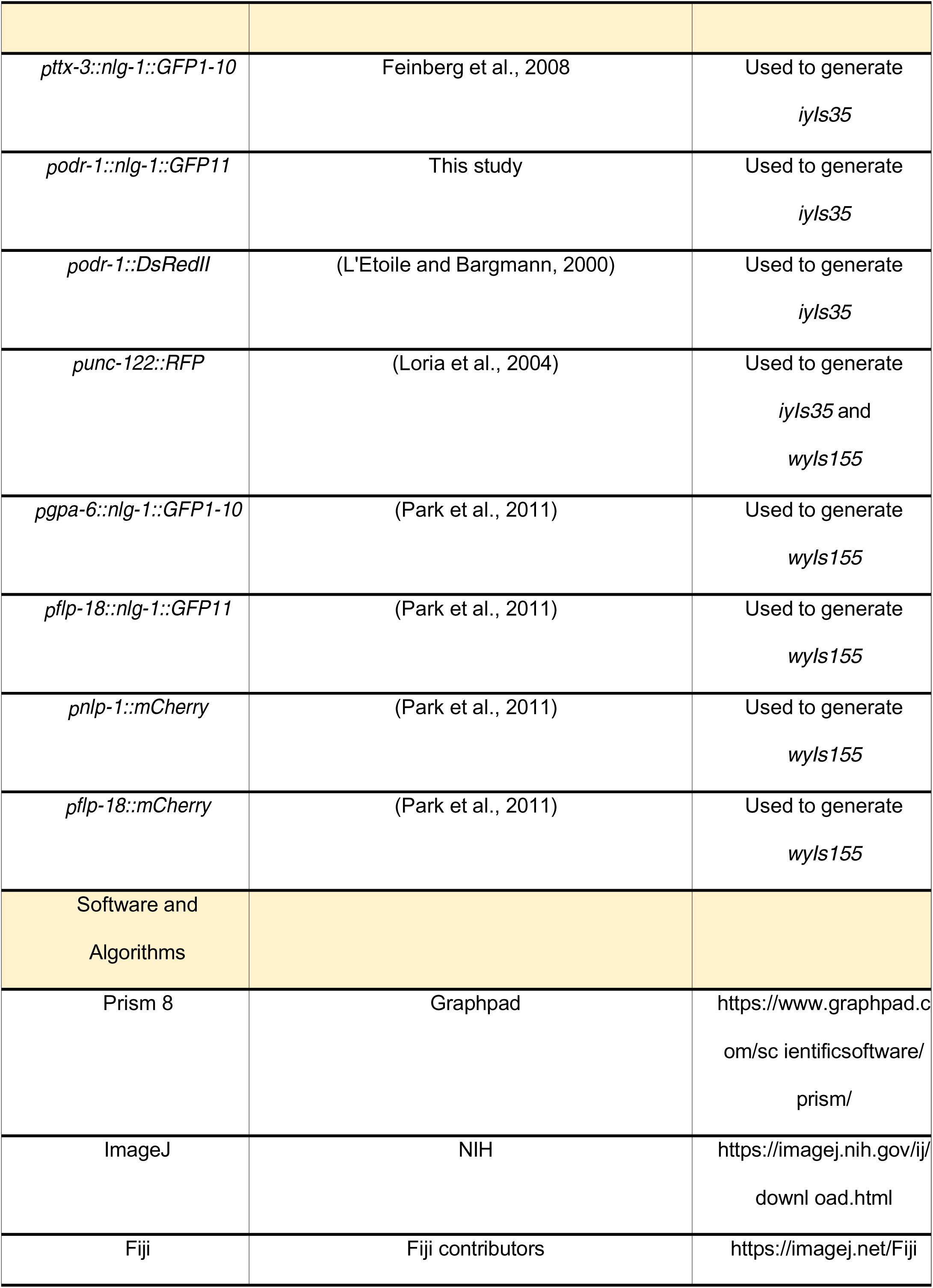
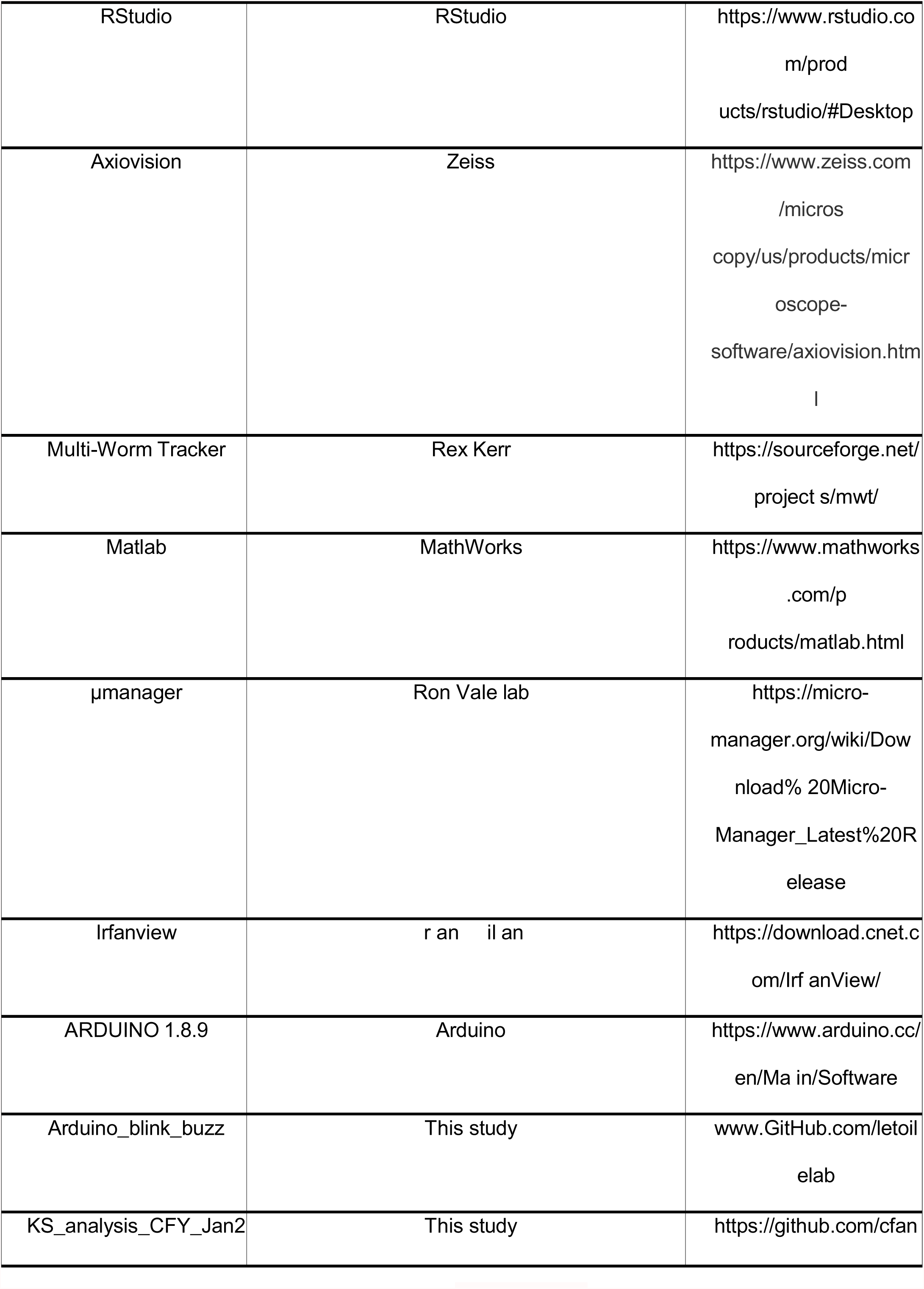
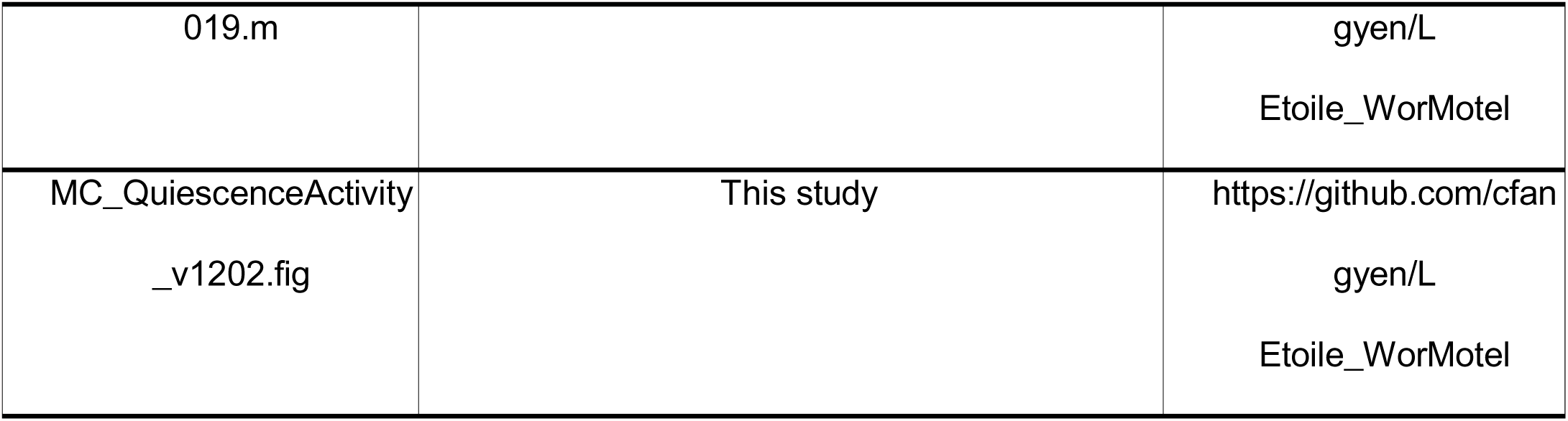

### RESOURCE AVAILABILITY

#### Lead Contact

Further information and requests for resources and reagents should be directed to and will be fulfilled by the Lead Contacts: Noelle L’Etoile for reagents in Figures 1-4 or Miri VanHoven for reagents in Figures 5-7.

### EXPERIMENTAL MODEL AND SUBJECT DETAILS

#### *C. elegans* strain cultivation

All *C. elegans* worms were reared according to standard protocols (Stiernagle, 2006). All strains were raised and tested at 20°C. Animals were raised on 10cm Nematode Growth Media (NGM) plates with an OP50 *E. coli* lawn. All assays were started when animals were day one gravid adults. All strains used in this study are listed in the Key Resources Table. The JZ2008 AIB and AIY double kill strain (*ttx-3(ks5) X; peIs578 (_p_npr-9::casp1; _p_unc-122::mCherry; _p_npr-9::venus)* was made by mating the *FK134/ttx-3(ks5) X* strain with the *peIs578 (_p_npr-9::casp1; _p_unc-122::mCherry; _p_npr-9::venus)* strain from the CGC and the Iino lab, respectively. Other transgenic strains generated for this study are *iyIs35 (_p_ttx-3::nlg-1::GFP1-10* (Feinberg *et al*., 2008) (70ng/μl)*, podr-1::nlg-1::GFP11* (see below for generation) (40ng/μl)*, _p_odr-1::DsRedII* (L’Etoile and Bargmann, 2000) (5ng/μl) and *_p_unc-122::RFP* (Loria et al., 2004) (20ng/μl)) and *wyIs155 (_p_gpa-6::nlg-1::GFP1-10* (Park et al., 2011) (60 ng/μl), *_p_flp-18::nlg-1::GFP11* (Park *et al*., 2011) (30 ng/μl)*, _p_nlp-1::mCherry* (Park et al., 2011) (10 ng/μl), *_p_flp-18::mCherry* (Park et al., 2011) (5 ng/μl) and *podr-1::DsRedII* (20 ng/μl)). Constructs were generated using standard molecular techniques. To generate *_p_odr-1::nlg-1::GFP11,* the *odr-1* promoter was amplified from *_p_odr-1::DsRedII* (L’Etoile and Bargmann, 2000) using *_p_odr-1-*specific primers (MVP578: TTGCATGCCTGCAGGTCG, which has an internal SphI site and MVP581: GACTGGCGCGCCTACCTTTGGGTCCTTTGGC, which introduces an AscI site). Then, the *_p_odr-1* fragment was subcloned into the SphI-AscI fragment from *nlg-1::GFP11* (Park *et al*., 2011)

### METHOD DETAILS

#### LTM assay (Figures 1-4)

The odor training protocol contains three 80-minute cycles of training with odor or a control buffer interspersed with two 30-minute periods of feeding with OP50 *E. coli* bacteria. One day old adult worms were washed with S Basal buffer (0.1M NaCI, 0.05M K_3_PO_4_, pH 6.0) off of 10 cm NGM plates and into microfuge tubes, where they were washed three times with S Basal buffer. The animals were split in two groups and one group was added to a microfuge tube of S Basal and the other group was added to a microfuge tube of 1:10,000 dilution of butanone in S Basal. The microfuge tubes were then rotated on a rotisserie for 80 minutes.

To make the concentrated OP50 for the 30-minute feeding cycles, 100 mL of LB was seeded with OP50 and was shaken overnight at 37°C at approximately 250 RPM until it reached an OD of approximately 10 and then centrifuged at 4000 RPM for 15-30 minutes or allowed to settle for 1 hour, and the pellet was resuspended in 32 mL of S Basal. Animals were washed three times with S Basal and then added to a microfuge tube with 750-1000 µL of concentrated OP50, and rotated for 30 minutes. Animals are then washed three times with S Basal, allowing the animals to settle five minutes each time, before they begin their next 80- minute cycle of training.

After the third treatment cycle, worms were washed three times with S Basal and placed on NGM plates with OP50. When indicated in figures, the animals were split into different groups and assayed for chemotaxis (see below), quiescence or NLG-1 GRASP fluorescence intensity. Long-term memory was assessed 16 hours after training was completed.

For single-cycle odor training (performed in Figure 3), the adult worms were washed with S Basal from the 10 cm NGM plates and trained for 80 minutes. Variations of the LTM assay are described in Figure 3.

#### Chemotaxis assay

Chemotaxis plates were made using these materials (for 100 mL of media): 100 mL ddH_2_O with 1.6 g of Difco agar, agar agar, or bacto agar, 500 µL of 1M K_3_PO_4_ solution (pH 6.0), 100 µL 1M CaCl_2_ and 100 µL 1M MgSO_4_. The agar was boiled or autoclaved to dissolve uniformly in 100 mL ddH_2_O and cooled to 57-53°C (to avoid precipitation) before adding the salt solutions. 10 mL of media was poured into 10 cm plastic petri dishes and let it cool to solidify. We drew assay plate guides as shown in Figure 1A. 1 µL of (1 M) NaN_3_ was pipetted onto the middle of the odor and diluent (control) arenas. We then added 1 µL of 200 proof ethanol to the diluent arena. To the odor arena, we added 1 µL of 1:1000 butanone, 1 µL of 1:200 benzaldehyde or 1 µL of 1:1000 diacetyl.

One day old adults grown at 20°C on 10cm NGM plates with OP50 *E. coli* lawns were trained and used for chemotaxis assays. It is critical that the strains be completely clean with no fungal and bacterial contamination of any kind. When plating animals after three cycles of training, we did two washes with S Basal buffer and then a third wash with S Basal or ddH_2_O. We plated 50-400 animals at the origin of the plate (bottom) and wicked away excess moisture with a Kim Wipe, being careful not to cause any gouges in the agar, which can cause burrowing. Worms were allowed to roam at least 90 minutes.

To calculate a chemotaxis index (CI), we counted how many worms were in each arena, and how many total worms there were on the plate outside the origin. We subtracted the number of worms in the diluent arena from the number in the odor arena, and then divided that by the total number of worms on the plate that were not at the origin. We censored the assays in which the buffer-trained populations exhibited a CI of less than 0.5, as this indicates that the worms were unable to chemotax to the odor for some reason.

#### Sleep analysis

To make the WorMotel (Churgin *et al*., 2017), we used a PDMS chip with 48 total wells made according to the paper or the online resource (http://fangyenlab.seas.upenn.edu/links.html). Next, we made 100mL of NGM by adding together 1.8g low melting-point agarose, 0.3g NaCl, 0.25g bacto peptone, and 1µL tween 20 (to keep a flat agar surface). We boiled the media in the microwave, cooled it down to ∼50-58°C, then added 100µL CaCl_2_, 100 µL cholesterol dissolved in EtOH, 100 µL MgSO_4_, and 2.5 mL K_3_PO_4_. We next added 17 µL/well of chip and let it cool. The solidified WorMotel was placed in a transparent Petri dish with 10mg of gel soil (Soil Moist granules) soaked in 100 mL of water to prevent cracking of wells. Four clay balls were used to prop the lid open uniformly for 1-8 hours or as long as the worms were assayed. Food from NGM/OP50 plate was scooped and smeared on top of the wells to prevent the worms from being starved.

Teledyne Dalsa PT-21-04M30 Camera Link Camera (Dalsa Proprietary Sensor 2352x1728 Resolution Area Scan Camera) attached with a Linos Rodagon Modular-Focus lens (f=60mm) was used to image the entire WorMotel. To obtain a focused working distance, four metal posts with a plastic stage was built. A T175 tissue culture flask filled entirely with water was added to make a cooling chamber as well as a light diffuser (water diffracts light). We used the Multiple-Worm Tracker (MWT 1.3.0r1041) made by Rex Kerr to automate image capture and record worm movement every 3 seconds for the entire duration of the experiment. Irfanview (developed by Irfan Škiljan) was used to re-index the images for sequence verification before quantifying the movements in MATLAB. Once indexed, the images were batch processed in MATLAB for thresholding each worm uniformly to quantify quiescence using a graphic user interface (GUI) created by MC and CFY and available at https://github.com/LEtoileLab/Sleep_2022.git. After thresholding, the animals were quantified for quiescence and activity using another MATLAB code available at https://github.com/LEtoileLab/Sleep_2022.git

#### Statistical analysis

For Figures 1-4 and S1-S4, statistics were performed using GraphPad Prism 8 and Rstudio. *P*-values are used for the statistical readouts, with the following notations: NS P>0.05, * P<0.05, ** P<0.01, *** P<0.001, and **** P<0.0001. All data included in the same graph were analyzed for type of data distribution with the Shapiro-Wilk normality test. If datasets were normally distributed, then one-way ANOVA was used for multiple comparisons, followed by Bonferroni’s multiple correction. If the data were found to be non-normal, then the non-parametric Kruskal-Wallis test was used to analyze differences in means, followed by pairwise comparisons using the Mann-Whitney two-tailed U-test. Then, to correct for Type I error, the Hochberg test was run on p-values compared in the same graph to adjust the p-values for multiple comparisons, which often conservatively increases p-values to avoid incorrectly rejecting the null hypothesis. For graphs with only two datasets to compare, the Shapiro-Wilk test was performed, followed by the t-test or U-test, depending on the distribution of the datasets. For correlation data, Pearson’s correlation test was used. The standard error of the mean (SEM) was calculated and shown on each graph. Specific statistical tests used for each graph in the manuscript are included in the figure legends.

For Figures 5-7 and S6-7, statistics were performed using R and Excel. *P*-values are similarly used for the statistical readouts. For normally distributed data, one-way ANOVA was used followed by pair-wise t-tests. For nonparametric data, the Kruskal-Wallis test was used, followed by pair-wise U-tests. If more than one t-test or U-test was conducted, the resulting *p*-values were adjusted for multiple comparisons using the Hochberg method, which is standardly applied to adjust for the tendency to incorrectly reject a null hypothesis when multiple comparisons are made, and can only conservatively increase *p*-values.

#### Movement analyses

We assessed animal responsiveness to a stimulus which has been previously shown to disrupt quiescence in *C. elegans* (Nagy et al., 2014). Specifically, 3.5 cm or 5.5 cm diameter NGM plates with OP50 lawns with 20- 30 worms were placed on a 50mm piezo 1.2 KHZ piezo buzzer elements (Digikey #668- 1190-ND). Piezo elements were supplied with 5V with a 50% pulse-width modulated duty cycle using an Arduino-style microcontroller and its accompanying software, using the code named “Arduino_blink_buzz” accessible at www.GitHub.com/letoilelab. Stimulus onset was synchronized with video recording by flashing a blue LED, used at the maximum light intensity (we used Digikey #1528-2334-ND) at a distance of about 10cm during video recording. In cases where animals were exposed to prolonged stimulation, animals were subjected to blocks of stimulation for 5 minutes with the blue light flashing for 1 second every 20 seconds, followed by no stimulation for 5 minutes. Videos were recorded on an Imaging Source DMK 23GP031 camera using Micromanager software (Edelstein et al., 2014)

#### Pharyngeal Pumping Assay

This assay was performed by standard methods (Raizen et al., 2012).To perform the assay, we watched the pharynx of a worm under a stereomicroscope at 100-200X magnification and once the grinder in the terminal bulb does one complete contraction and relaxation, or “pump”, we counted that as one pump, using a counter to count every time they complete a full pump for 15 seconds. Then, we disposed of the worm to prevent re- counting of the same animal. We took the number of pumps completed and multiplied that by 6 to find the pharyngeal pumping rate in pumps per minute.

#### Quiescence disruption

Post-training quiescence was disrupted by either mechanical disruption or removal from food for two hours. For mechanical disruption, after training the animals were divided into four different groups in addition to performing sleep and chemotaxis assays. The first group was not disturbed for 16 hours, while the second, third and fourth group were mechanically disturbed by plating the worms in a dense bacterial slurry, which was shaken every 15 minutes to ensure uniform movements of the worms. The bacterial slurry was made by centrifuging an overnight culture of OP50 (at 37°C with 250 RPM, OD=10) at 4000 RPM for 15-30 minutes to resuspend the pellet(s) in 5 mL of S Basal. The slurry plates were made by pouring 1 mL of dense bacterial slurry on NGM plates approximately 30 minutes before plating the worms. After mechanically disturbing the worms, the worms were rinsed three times with S Basal before moving them to NGM plates with OP50 bacterial lawns. The third and the fourth group of animals were initially placed on NGM plates with OP50 bacterial lawns for two or four hours, and then washed with S Basal three times before putting them on slurry plates. For removal from food assays, worms were placed on NGM plates without OP50 lawns for the first two hours after training before plating them on NGM plates with OP50 lawns for 14 hours.

To load the WorMotel with worms that did not have food, we placed the worms on NGM plates without food and loaded individual worms into the WorMotel using a pick. This WorMotel was divided into four groups to compare the amount of sleep, where one group of buffer- and butanone-trained animals received food after training and the other group of trained animals did not.

#### CFU measurements

After the 0-2 hours period of mechanical disturbance with green OP50, worms were washed three times with S Basal before treating 15 minutes with 200 μL 0.5% bleach solution in a 96-well plate. After 15 minutes of incubation at room temperature (20°C), the worms were washed three times with S Basal. The alimentary canal of the worms was dissected out using two 22 gauge hypodermic needles under the dissecting microscope and transferred to the other wells containing 200 μL SOC media, which was incubated at 37°C for 15 minutes before plating the 200 μL of SOC media on LB plates. As a negative control, worms were treated with a bleach solution but left undissected, and treated with SOC media like the dissected worms, which were then plated to confirm that there were no green colonies.

#### Heat shock assays

*C. elegans* animals were heat-shocked at 37°C for 5 minutes in a water bath while on 5.5cm unseeded NGM plates covered in Parafilm. After the heat shock, animals were put at 20°C until behavior was assessed by the chemotaxis assay.

#### Calcium imaging

Calcium imaging of the AWC^ON^ neuron was performed on lines expressing the genetically-encoded calcium indicator GcaMP3 (Tian *et al*., 2009) under the *str-2* promoter (*JZ1795/pyIs701(_p_str-2::GcaMP3; _p_ofm-1::GFP; _p_ceh-36::mCherry)*). One-day-old adult worms were conditioned to either buffer or 1.23mM butanone diluted in S Basal buffer (the same concentration as used for the butanone conditioning mentioned in “Chemotaxis assay”) during three, 80-minute training cycles (interspersed) with feeding (described in “LTM chemotaxis assay”). Immediately after the end of the third training cycle or after a 16-hour overnight period on food, worms were rinsed three times in S Basal buffer and loaded into a custom, polymer polydimethylsiloxane (PDMS) microfluidic device (Chronis et al., 2007). The nose of the animal was exposed to liquid streams of either S Basal buffer or 1.23mM butanone. A manual switch attached to a solenoid valve was used to direct the buffer or odor stream across the nose of the worm. The stimulation protocol consisted of exposing worms to S Basal buffer for 30 seconds followed by a 30-second exposure to 1.23mM butanone (odor on) or by exposing worms to 1.23mM butanone for 30 seconds followed by a 30 second exposure to S Basal buffer (odor off).

Fluorescence was monitored with a Zeiss 40X air objective on an inverted microscope (Zeiss Axiovert 200). Images were taken at 2 frames per second with a blue light exposure time of 100ms using an ORCA-Flash 2.8 camera (Hamamatsu).

#### Calcium imaging analysis

Fiji software was used with the Multi Measure plugin to analyze the images. In animals expressing the GcaMP3 reporter in the AWC^ON^ neuron (*JZ1795/pyIs701(_p_str-2::GcaMP3; _p_ofm- 1::GFP; _p_ceh-36::mCherry)*), the ROI was established at the center of the AWC cell body. A background ROI was also taken, just outside of the animals. Then, the mean fluorescence intensity at the background ROI was subtracted from the mean fluorescence intensity at the cell body ROI and that serves as the “F” values. The fluorescence intensity of the GcaMP3 reporter in the first three images is defined as F_0_.ΔF is the F_0_ value subtracted from each F value.

For every worm imaged, the mean of the ΔF_0_/F (%) values is taken from 10 seconds before and after the butanone is turned on or off (i.e. the means at 20-30 seconds and 30.5-40.5 seconds are taken). The delta is then taken between the two means and the absolute value is taken of that number for comparisons between the datasets (e.g. buffer vs butanone-trained cohorts) taken on the same day.

#### LTM NLG-1 Synaptic Imaging Assays

The LTM training paradigm described above was modified to accommodate NLG-1 GRASP imaging. Approximately 30 NGM plates of day one gravid adult *iyIs35* worms were prepared to allow enough worms for multiple batches and imaging. Worms were divided into four batches that began training 40 minutes apart.

Batches one and three were trained in a control buffer (S Basal), while batches two and four were trained in the odor solution. For all LTM NLG-1 GRASP synaptic imaging assays, images of animals were only analyzed for NLG-1 GRASP intensity from training batches that passed the behavioral batch chemotaxis tests.

Specifically, butanone-trained batches allowed to sleep were considered to pass the behavioral test if they were not attracted to butanone (CI<0.5), while butanone-trained batches with disrupted sleep and buffer-trained batches were considered to pass the behavioral test if they were attracted to butanone (CI>0.5).

For all NLG-1 GRASP LTM assays, A Zeiss Axio Imager.A1 compound fluorescent microscope (Figures 6C, 6D, 7B, S5C, S6A-B, and S7D) and a Zeiss LSM710 confocal microscope (Figure 5C, 6A, 6B, 7A, and S7A-B) were used to capture images of live *C*. *elegans* under 630X magnification. For batch assays (all assays except the single-worm time course assays), images were taken of approximately 20 animals from each batch within approximately 20 minutes of the time point.

LTM NLG-1 GRASP 16-hour mechanical disturbance assays: After three cycles of training (described above), half the worms from each batch were placed on NGM plates with an OP50 *E. coli* lawn. The other half were placed on plates with 1 mL of OP50 suspended in S Basal buffer (as described above) and tapped for one minute out of every 15 minutes for a two-hour period. Worms on these plates were then transferred to NGM plates with OP50 lawns after the two-hour period. All plates were incubated at 20°C until 16 hours after training, when worms were washed (as described above). 50-400 worms from each of the eight batches underwent the butanone chemotaxis assay. Worms were anesthetized for imaging on 2% agarose pads using a 2:1 ratio of 0.3 M 2,3-butanedione monoxime (BDM) and 10 mM levamisole in M9 buffer. All micrographs taken were of one-day old and two-day old gravid adults.

LTM NLG-1 GRASP 16-hour removal from food assays: After three cycles of training (described above), half the worms from each batch were placed on NGM plates with an OP50 *E. coli* lawn, and half were placed NGM plates without bacteria, then transferred to NGM plates with an OP50 *E. coli* lawn after two hours. All plates were incubated at 20°C until 16 hours after training, when worms were washed (as described above). Approximately twenty worms from each of the eight batches were anesthetized and imaged using a Zeiss Axio Imager.A1 compound fluorescent microscope and Axiovision software, as described above, and 50- 400 worms from each of the eight batches underwent the butanone chemotaxis assay.

LTM NLG-1 GRASP batch 0-hour, 2-hour, and 16-hour assays: After three cycles of training, worms from batches one to four were each divided into three groups so that imaging and chemotaxis experiments could be performed at three timepoints: zero hours after training, two hours after training, and 16 hours after training. 0-hour Imaging and Chemotaxis: After training, worms from all four batches were washed (as described above). From each batch, ∼20 worms were separated, anesthetized, and imaged under the Zeiss Axio Imager.A1 compound fluorescent microscope and Axiovision software (see “Synapse Imaging and Analysis” above), and 50 to 400 worms were assessed for butanone chemotaxis. Two-hour Imaging and Chemotaxis: After training, worms from batches one and two were each placed on NGM plates with an OP50 lawn. After two hours, animals were washed as described above. Approximately twenty animals from each of these batches were anesthetized and imaged, and 50 to 400 worms from each of these batches were assessed for butanone chemotaxis. 16-Hour Imaging and Chemotaxis: After training, worms from batches one, two, three, and four were each placed on NGM plates with OP50 lawns. After 16 hours, animals were washed as described above. Approximately twenty animals from each of these batches were anesthetized and imaged, and 50 to 400 worms from each of these batches were assessed for butanone chemotaxis.

LTM Single worm time course NLG-1 GRASP assays: The training paradigm detailed in the LTM NLG- 1 GRASP assays were repeated as described above with the following modifications following the three cycles of training. All worms from each batch were placed on NGM plates with OP50 lawns and gravid adults with only one row of eggs were selected for the single worm time course assay. Each animal was either imaged at 0 and 2 hours after training, or at 2 and 16 hours after training. Animals were anesthetized with 10mM tetramisole in a 1:1 ratio with M9 buffer or a 2:1 ratio of 0.3 M 2,3-butanedione monoxime (BDM) to 10mM tetramisole on agarose pads or NemaGel. After imaging at the first timepoint, each animal was picked from the agarose pad into an approximately 50 µL drop of M9 buffer on an NGM plate with an OP50 lawn. The animal was then transferred to a drop of M9 buffer on a second NGM plate with an OP50 lawn to aid the recovery from the anesthetic. The worms spent roughly 30 minutes total in M9 buffer. These plates were incubated at 20°C until the second timepoint.

For animals imaged at 0 and 2 hours after training, animals were again anesthetized and imaged as described above for the first time point. For animals imaged at 2 and 16 hours after training, at the 16-hour timepoint, each animal that was mobile (as demonstrated by movement tracks on the bacterial lawn) was tested for chemotaxis to butanone before imaging. Individual buffer-trained animals that were attracted to the odor, and butanone-trained animals that were not attracted to the odor were then imaged again as described above. Single animals were washed for five minutes each in a drop of S Basal buffer and a drop of ddH_2_O. Single animals were then placed on a single worm chemotaxis plate with the origin at the center of the plate. The single worm chemotaxis plates were made as previously described (see “Chemotaxis Assay” section above) but with the origin of the single worm placement in the center of the plate. The worm was allowed to roam on the plate for at least 10 minutes. For animals to pass this single-worm behavioral screen, buffer-trained worms needed to move directly towards butanone or stay on the butanone side of the plate the majority of the time, while butanone-trained worms needed to move and not make directed movement towards butanone nor spend the majority of time on the butanone side of the plate. Only animals from assays with two or more successful buffer- and butanone-trained animals that passed the single-worm behavioral screen at the 16-hour timepoint were included. Additionally, only micrographs of animals with their head approximately on its side at both timepoints were analyzed.

#### Synapse imaging analysis

NIH ImageJ software (Abramoff et al., 2003) was used to analyze all Zeiss Axio Imager.A1 compound fluorescent micrographs taken for AWC-AIY NLG-1 GRASP phenotypic quantification, as previously described (Park *et al*., 2011; Varshney *et al*., 2018). In brief, AWC-AIY NLG-1 GRASP intensity was quantified by measuring (Collingridge *et al*., 2010) fluorescence intensity through circling punctal clusters. In Figures 6C, 6D, 7B, S5C, and S6B median intensity values for each treatment were normalized to fluorescence intensity levels for buffer-trained animals placed on food for 16 hours on the same day. In Figure S7D, for the single-worm assays, the median intensity of each worm at the first timepoint was normalized to 100% and the difference in intensity between the first and second timepoint was quantified for each animal. Since animals that are imaged twice can undergo photobleaching, we assessed the proportion of animals with greater than or equal to 50% reduction in fluorescence intensity between the two timepoints.

PHB-AVA NLG-1 GRASP intensity in Figure S6B was also measured by outlining clusters of puncta. Background fluorescence intensity near the gut was also taken into account by calculating the minimum intensity value in the area directly surrounding the puncta. This background intensity value was subtracted from the intensity for each pixel in the punctal cluster, and the adjusted values were added as previously described (Park *et al*., 2011; Varshney *et al*., 2018).

All team members performing image analysis were blind to the animals’ prior conditioning. For batch LTM NLG-1 GRASP statistical analysis in Figures 6C, 6D, 7B, and S6B, the Kruskal-Wallis test was used to analyze variance between treatments. If the p-value was less than 0.05, then the Mann-Whitney U-test was used to compare the medians of each pair of groups, followed by the Hochberg multiple comparison procedure. For single worm LTM NLG-1 GRASP statistical analysis in Figure S7D, we performed two-independent sample Z-tests.

## ACKNOWLEDGMENTS

We would like to thank all of the L’Etoile, VanHoven, Kato, and Fang-Yen lab members for all of their helpful discussions, especially Katie Mellman, AJ Ablaza, Mary Futey, Sarah Woldemariam, Bo Zhang, Trang Duong, and Aarati Asundi. We would also like to thank Sara Alladin, Nebat Ali, Anjana Baradwaj, Claudia Echeverria, Jordan Mitchell, and Idan Siman-Tov for help with synapse experiments. We would like to thank Veronica Valdez for making the reagents to support the experiments in this manuscript. We are thankful to Andrei Goga, Maria Gallegos, Torsten Wittmann, Matt Gruner, Stephen Nurrish for all for all of their advice on experiments in this manuscript. We give a big thanks to the *Caenorhabditis* Genetics Center (CGC), supported by the NIH Office of Research Infrastructure Programs (P40 OD010440), for the strains they provided to us and for their support of the greater *C. elegans* community. We would also like to give another big thanks to the Sternberg lab and everyone else behind Wormbase.org (NIH grant #U24 HG002223) and the Hall lab for Wormatlas.org (NIH grant #OD010943 to DHH), both for their immensely helpful resources. Thanks to BioRender for their artwork which was used in Figure 1 and S1. This work was funded by the NIH, specifically R01DC005991 (N.L.D., M.K.V., C.F.Y.), R15NS109803 (M.K.V.), R01NS087544 (N.L.D., M.K.V.), R35GM124735 (S.K.), F31DC014921 (K.L.B.), and R01NS084835 (M.A.C.).

---

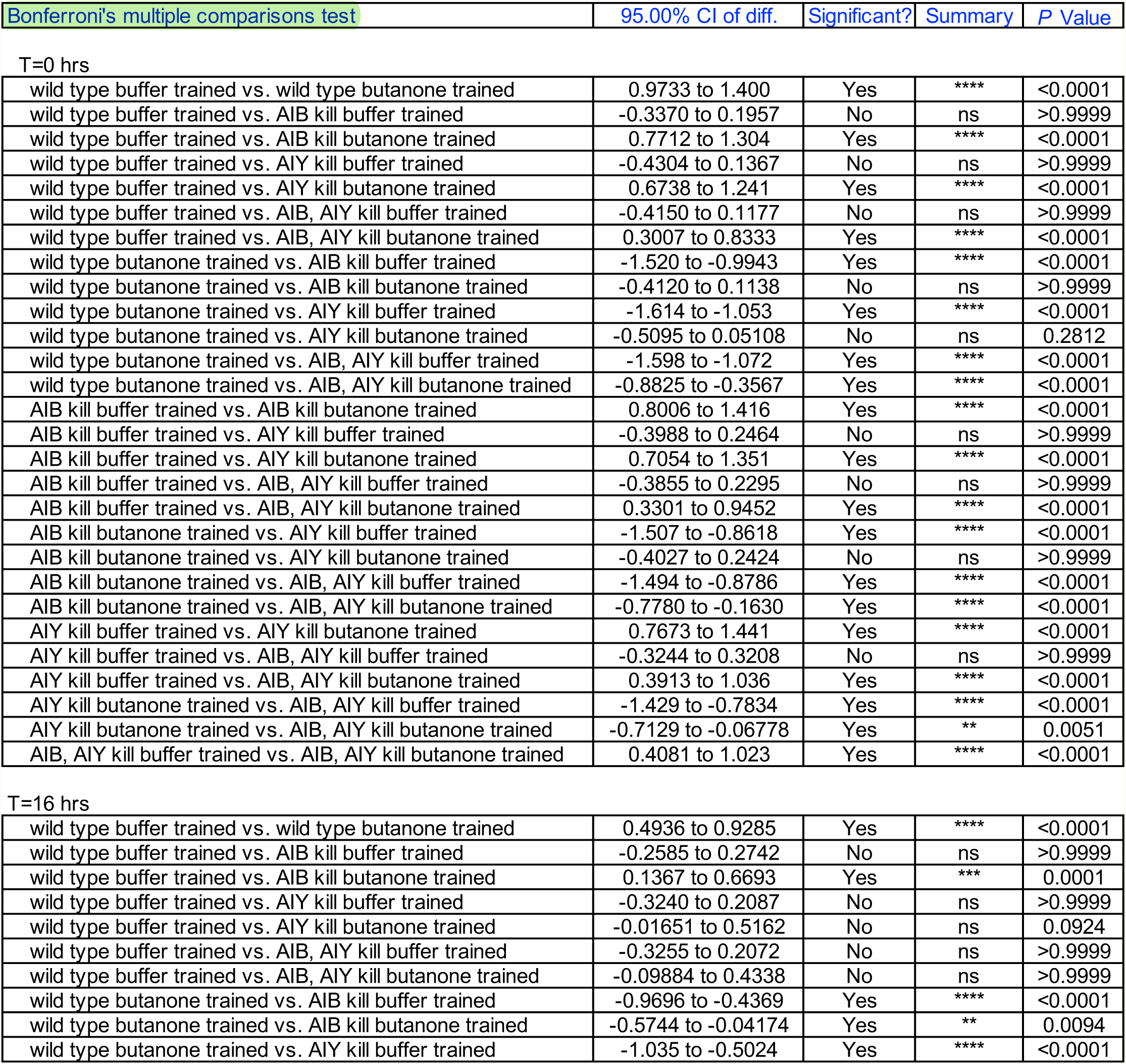
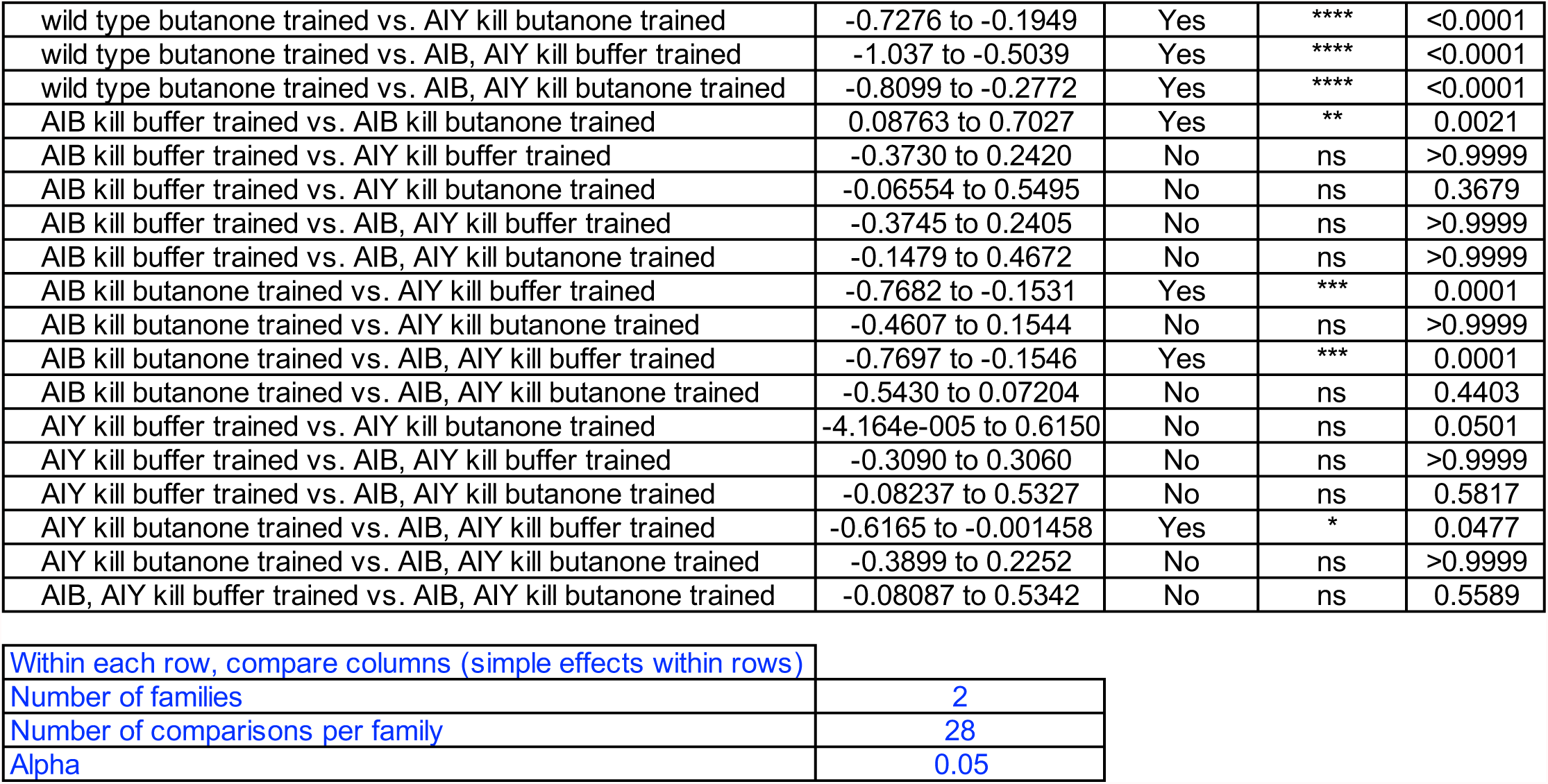

| Bonferroni's multiple comparisons test | 95.00% CI of diff. | Significant? | Summary | P Value |
| --- | --- | --- | --- | --- |
| T=0 hrs |  |  |  |  |
| wild type buffer trained vs. wild type butanone trained | 0.9733 to 1.400 | Yes | **** | <0.0001 |
| wild type buffer trained vs. AIB kill buffer trained | -0.3370 to 0.1957 | No | ns | >0.9999 |
| wild type buffer trained vs. AIB kill butanone trained | 0.7712 to 1.304 | Yes | **** | <0.0001 |
| wild type buffer trained vs. AIY kill buffer trained | -0.4304 to 0.1367 | No | ns | >0.9999 |
| wild type buffer trained vs. AIY kill butanone trained | 0.6738 to 1.241 | Yes | **** | <0.0001 |
| wild type buffer trained vs. AIB, AIY kill buffer trained | -0.4150 to 0.1177 | No | ns | >0.9999 |
| wild type buffer trained vs. AIB, AIY kill butanone trained | 0.3007 to 0.8333 | Yes | **** | <0.0001 |
| wild type butanone trained vs. AIB kill buffer trained | -1.520 to -0.9943 | Yes | **** | <0.0001 |
| wild type butanone trained vs. AIB kill butanone trained | -0.4120 to 0.1138 | No | ns | >0.9999 |
| wild type butanone trained vs. AIY kill buffer trained | -1.614 to -1.053 | Yes | **** | <0.0001 |
| wild type butanone trained vs. AIY kill butanone trained | -0.5095 to 0.05108 | No | ns | 0.2812 |
| wild type butanone trained vs. AIB, AIY kill buffer trained | -1.598 to -1.072 | Yes | **** | <0.0001 |
| wild type butanone trained vs. AIB, AIY kill butanone trained | -0.8825 to -0.3567 | Yes | **** | <0.0001 |
| AIB kill buffer trained vs. AIB kill butanone trained | 0.8006 to 1.416 | Yes | **** | <0.0001 |
| AIB kill buffer trained vs. AIY kill buffer trained | -0.3988 to 0.2464 | No | ns | >0.9999 |
| AIB kill buffer trained vs. AIY kill butanone trained | 0.7054 to 1.351 | Yes | **** | <0.0001 |
| AIB kill buffer trained vs. AIB, AIY kill buffer trained | -0.3855 to 0.2295 | No | ns | >0.9999 |
| AIB kill buffer trained vs. AIB, AIY kill butanone trained | 0.3301 to 0.9452 | Yes | **** | <0.0001 |
| AIB kill butanone trained vs. AIY kill buffer trained | -1.507 to -0.8618 | Yes | **** | <0.0001 |
| AIB kill butanone trained vs. AIY kill butanone trained | -0.4027 to 0.2424 | No | ns | >0.9999 |
| AIB kill butanone trained vs. AIB, AIY kill buffer trained | -1.494 to -0.8786 | Yes | **** | <0.0001 |
| AIB kill butanone trained vs. AIB, AIY kill butanone trained | -0.7780 to -0.1630 | Yes | **** | <0.0001 |
| AIY kill buffer trained vs. AIY kill butanone trained | 0.7673 to 1.441 | Yes | **** | <0.0001 |
| AIY kill buffer trained vs. AIB, AIY kill buffer trained | -0.3244 to 0.3208 | No | ns | >0.9999 |
| AIY kill buffer trained vs. AIB, AIY kill butanone trained | 0.3913 to 1.036 | Yes | **** | <0.0001 |
| AIY kill butanone trained vs. AIB, AIY kill buffer trained | -1.429 to -0.7834 | Yes | **** | <0.0001 |
| AIY kill butanone trained vs. AIB, AIY kill butanone trained | -0.7129 to -0.06778 | Yes | ** | 0.0051 |
| AIB, AIY kill buffer trained vs. AIB, AIY kill butanone trained | 0.4081 to 1.023 | Yes | **** | <0.0001 |
| T=16 hrs |  |  |  |  |
| wild type buffer trained vs. wild type butanone trained | 0.4936 to 0.9285 | Yes | **** | <0.0001 |
| wild type buffer trained vs. AIB kill buffer trained | -0.2585 to 0.2742 | No | ns | >0.9999 |
| wild type buffer trained vs. AIB kill butanone trained | 0.1367 to 0.6693 | Yes | *** | 0.0001 |
| wild type buffer trained vs. AIY kill buffer trained | -0.3240 to 0.2087 | No | ns | >0.9999 |
| wild type buffer trained vs. AIY kill butanone trained | -0.01651 to 0.5162 | No | ns | 0.0924 |
| wild type buffer trained vs. AIB, AIY kill buffer trained | -0.3255 to 0.2072 | No | ns | >0.9999 |
| wild type buffer trained vs. AIB, AIY kill butanone trained | -0.09884 to 0.4338 | No | ns | >0.9999 |
| wild type butanone trained vs. AIB kill buffer trained | -0.9696 to -0.4369 | Yes | **** | <0.0001 |
| wild type butanone trained vs. AIB kill butanone trained | -0.5744 to -0.04174 | Yes | ** | 0.0094 |
| wild type butanone trained vs. AIY kill buffer trained | -1.035 to -0.5024 | Yes | **** | <0.0001 |
| wild type butanone trained vs. AIY kill butanone trained | -0.7276 to -0.1949 | Yes | **** | <0.0001 |
| wild type butanone trained vs. AIB, AIY kill buffer trained | -1.037 to -0.5039 | Yes | **** | <0.0001 |
| wild type butanone trained vs. AIB, AIY kill butanone trained | -0.8099 to -0.2772 | Yes | **** | <0.0001 |
| AIB kill buffer trained vs. AIB kill butanone trained | 0.08763 to 0.7027 | Yes | ** | 0.0021 |
| AIB kill buffer trained vs. AIY kill buffer trained | -0.3730 to 0.2420 | No | ns | >0.9999 |
| AIB kill buffer trained vs. AIY kill butanone trained | -0.06554 to 0.5495 | No | ns | 0.3679 |
| AIB kill buffer trained vs. AIB, AIY kill buffer trained | -0.3745 to 0.2405 | No | ns | >0.9999 |
| AIB kill buffer trained vs. AIB, AIY kill butanone trained | -0.1479 to 0.4672 | No | ns | >0.9999 |
| AIB kill butanone trained vs. AIY kill buffer trained | -0.7682 to -0.1531 | Yes | *** | 0.0001 |
| AIB kill butanone trained vs. AIY kill butanone trained | -0.4607 to 0.1544 | No | ns | >0.9999 |
| AIB kill butanone trained vs. AIB, AIY kill buffer trained | -0.7697 to -0.1546 | Yes | *** | 0.0001 |
| AIB kill butanone trained vs. AIB, AIY kill butanone trained | -0.5430 to 0.07204 | No | ns | 0.4403 |
| AIY kill buffer trained vs. AIY kill butanone trained | -4.164e-005 to 0.6150 | No | ns | 0.0501 |
| AIY kill buffer trained vs. AIB, AIY kill buffer trained | -0.3090 to 0.3060 | No | ns | >0.9999 |
| AIY kill buffer trained vs. AIB, AIY kill butanone trained | -0.08237 to 0.5327 | No | ns | 0.5817 |
| AIY kill butanone trained vs. AIB, AIY kill buffer trained | -0.6165 to -0.001458 | Yes | * | 0.0477 |
| AIY kill butanone trained vs. AIB, AIY kill butanone trained | -0.3899 to 0.2252 | No | ns | >0.9999 |
| AIB, AIY kill buffer trained vs. AIB, AIY kill butanone trained | -0.08087 to 0.5342 | No | ns | 0.5589 |

|  |  |
| --- | --- |
| Within each row, compare columns (simple effects within rows) |  |
| Number of families | 2 |
| Number of comparisons per family | 28 |
| Alpha | 0.05 |

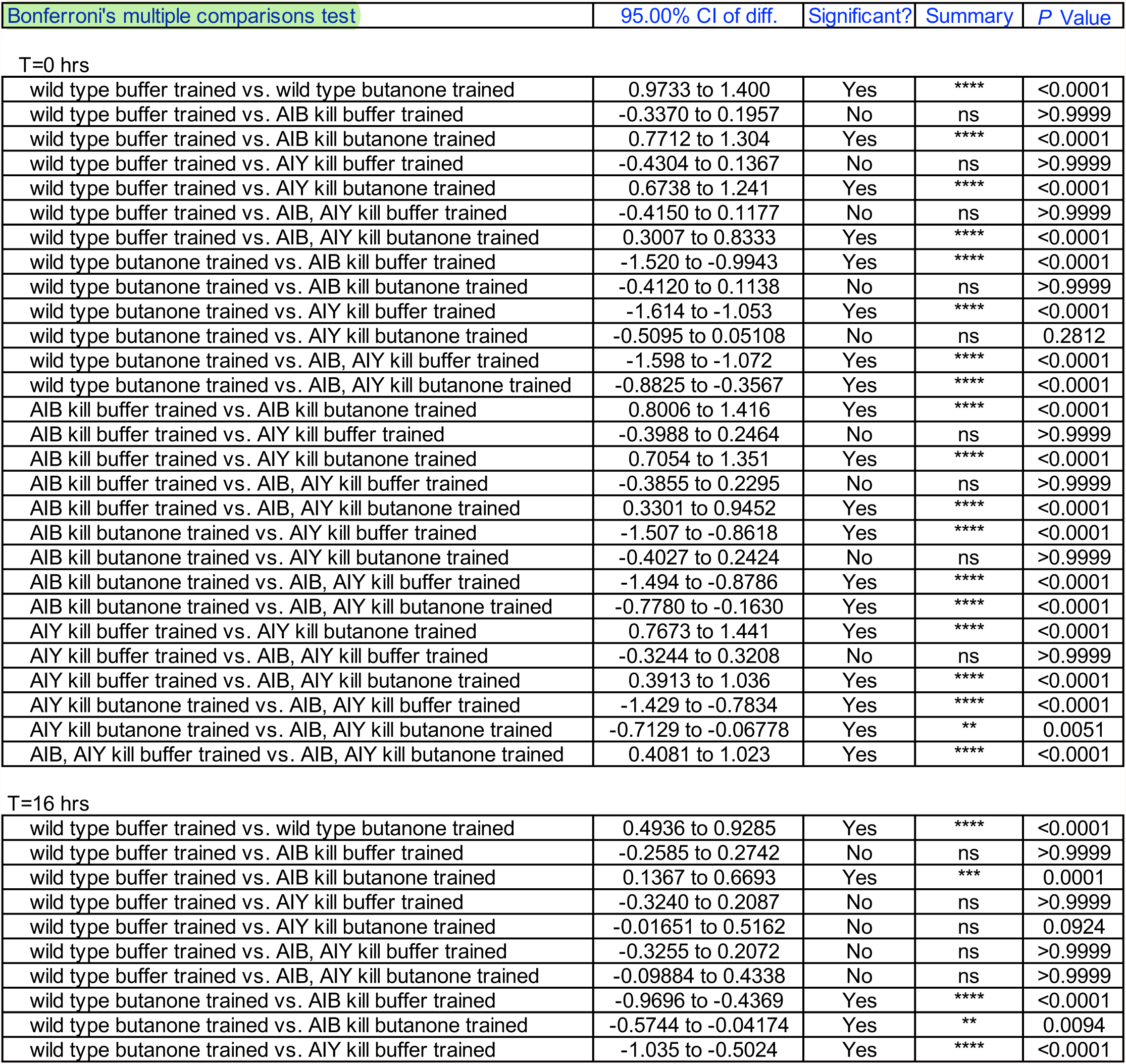
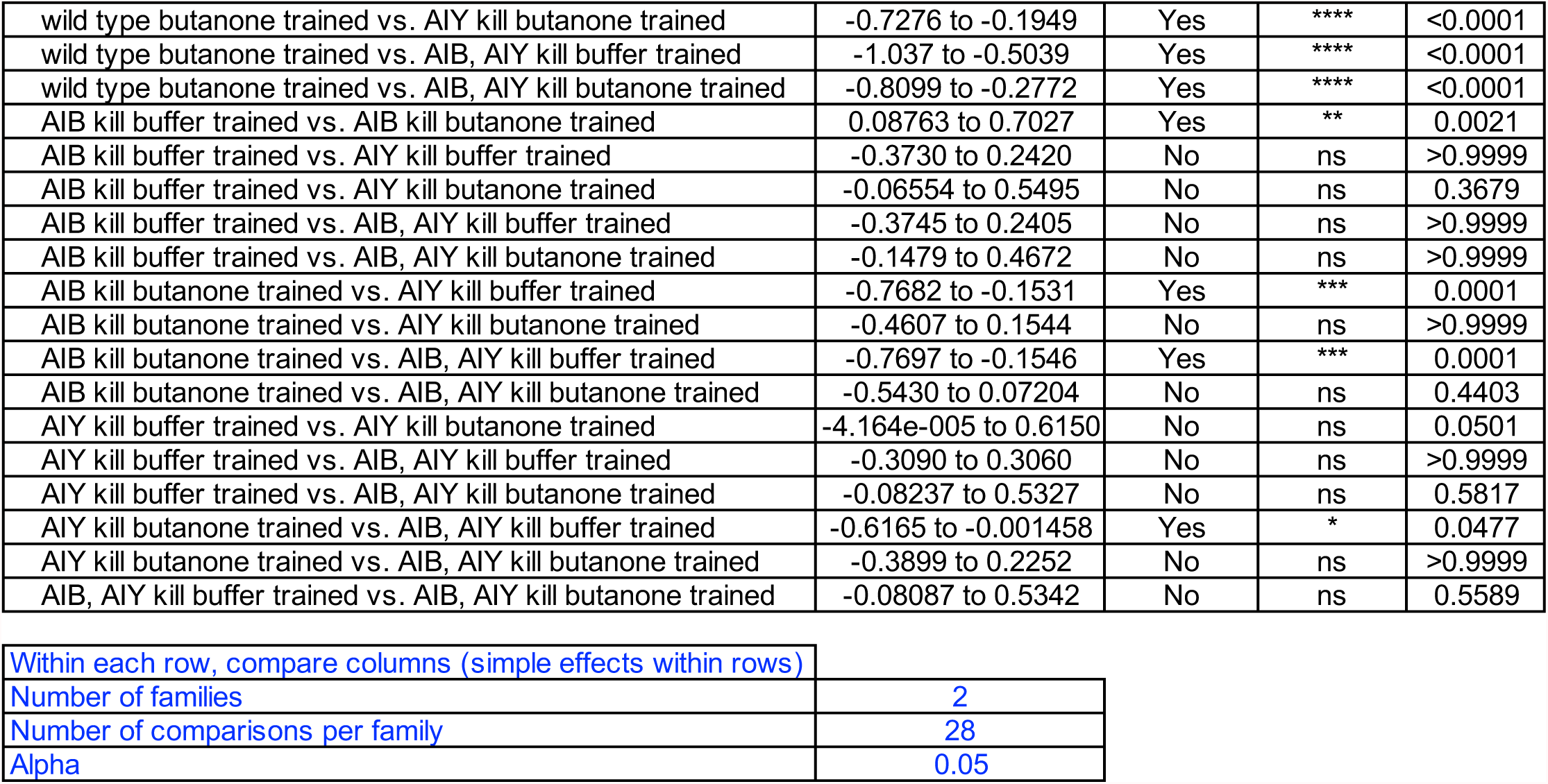

## Notes

https://ucsf.box.com/s/swph47ahmhns33g1ydozxkcukzk1scvr

https://ucsf.box.com/s/80ewoinw5tsdlkgne9peycanq18lsi5o

https://ucsf.box.com/s/mr2it3jyrszo9aqc0xytk8sv4bmqkl9g

https://github.com/LEtoileLab/Sleep_2022.git.

### Competing Interest Statement

The authors have declared no competing interest.

### Summary of Updates

This version has been revised to update Figures 1-4. These new data resulted from the efforts of the following new authors: Angel Garcia,Kateryna Tokalenko, Emily Soohoo, Vanessa Garcia, Sukhdeep Kaur, Malcolm Harris, Fabiola Briseno, Brandon Fung, Andrew Bykov, Hazel Guillen, Decklin Byrd, Emma Odisho and Kevin Daigle.

